# Adeno-Associated Virus Co-Precipitation with Extracellular Vesicles for Genome Editing in Rodent Embryo

**DOI:** 10.64898/2026.05.04.722478

**Authors:** Petr Nickl, Maria Barbiera, Jacopo Zini, Tereza Nickl, Aki Ushiki, Michaela Vaskovicova, Jitka Neburkova, Vojtech Dolejs, Michaela Simova, Jana Balounova, Petr Vyletal, Martina Zivna, Stanislav Kmoch, Zuzana Sumbalova-Koledova, Dominik Filipp, Ondrej Ballek, Veronika Niederlova, Ondrej Stepanek, Nadav Ahituv, Marjo Yliperttula, Radislav Sedlacek

## Abstract

Adeno-associated virus purification by density-gradient ultracentrifugation is labor-intensive and often results in substantial titer loss due to particle aggregation. Here, we present a scalable co-isolation strategy in which AAV is precipitated together with extracellular vesicles secreted by the producer cell line, completely bypassing density-gradient separation. The resulting AAV-EV preparations comprise free AAV, free EVs, and EV-associated AAV. Functionally, AAV-EV vectors (AAV2/1 serotype) support efficient *ex vivo* genome editing across multiple independent *loci* in mouse and rat zygotes, achieving a mean targeting efficiency of approximately 26%. Compared with gradient-purified AAV administered at matched doses, AAV-EV formulations yielded 2.34-fold higher embryo viability while maintaining equivalent transgene copy numbers. By leveraging EVs as a biological matrix, this approach enables ultracentrifugation-free AAV isolation without compromising vector functionality. Overall, AAV-EV represents an accessible and embryo-tolerant platform for rodent genome engineering that aligns with the principles of Replacement, Reduction, and Refinement (3R) principles.

## 1. Introduction

Recombinant adeno-associated virus (AAV) is widely used in gene therapy for its low immunogenicity and long-term expression with minimal genome integration [1]. AAV has also demonstrated high efficiency as a delivery vector for homology-directed repair (HDR) templates in rodent embryos, particularly for serotypes AAV2/1 and AAV2/6 [2–4]. In mammalian production systems, recombinant AAV is commonly generated using a helper-free approach based on triple co-transfection of transgenic HEK293 cells (in this study 293AAV cells) with transfer, Helper, and Rep/Cap plasmids [5] .

Although this approach can yield high viral titers, downstream isolation and purification remain major bottlenecks. Standard methods such as affinity chromatography and density gradient ultracentrifugation are labor-intensive and can contribute to virus aggregation or loss of functional particles [6–8]. Discontinuous density gradient ultracentrifugation is frequently used to separate AAV particles based on buoyant density. The workflow can be technically demanding, requires specialized equipment, and necessitates an additional ultrafiltration step to remove the density medium [8]. The removal of the density medium is particularly critical step, associated with titer loss or virus aggregation, which can result in poor titer retrieval and non-functional virus [9].

Extracellular vesicles (EVs) are naturally secreted proteolipid nanoparticles, including apoptotic bodies, microvesicles, and exosomes, which mediate intercellular communication [10]. Their biocompatibility makes EVs promising vehicles for therapeutic delivery [11, 12]. EVs are typically isolated from conditioned medium by differential centrifugation [13]. It has been reported that AAV capsids can associate on the surface or in the lumen of EVs, resulting in a vector with enhanced transduction efficiency and increased resistance to neutralizing antibodies compared to standard AAV particles [14]. EV association may arise from AAV internalization into the endosomal system during transduction, or they can be also associated during replication stage via the export of mature capsids during apoptosis or exosome-mediated secretion. Additional incorporation can occur during microvesicle budding at the plasma membrane [15–17]. The close interplay between AAV biogenesis and endosomal trafficking enables the generation of EV-associated AAVs [18].

The association of AAVs with EVs offer several advantages over conventional AAV vectors. It combines a transient EV-mediated delivery of RNAs and proteins, and long-term expression mediated by AAV [17]. EV-associated AAVs can be rapidly isolated, display resistance to neutralizing antibodies, are amenable to pseudo-typing for targeted delivery or blood-brain barrier penetration, and support diverse cargo-loading strategies [19].

Advances in EV-associated AAV engineering have focused on modifying vesicle composition and improving production efficiency. Pseudo-typing with vesicular stomatitis virus glycoprotein G (VSV-G) or rabies virus glycoprotein (RVG) can broaden tropism and increases enrichment of EVs in the preparation [14, 19]. Furthermore, an overexpression of the tetraspanin CD9 enhances exosome-associated AAV production [20].

Here, we present a simplified method of AAV isolation using EVs, and further we demonstrate improved *ex vivo* application of the acquired vectors. We refer to this isolation product as AAV-EV, as AAV is the particle of interest and EVs are the biological scaffold mediating the isolation.

AAV-EV is a composite formulation consisting of free AAV, free EV and EV-associated AAVs. Our isolation method streamlines production while maintaining functional performance in transgenic rodent model generation. The workflow consists of four steps:

1) **Continuous Medium Collection:** Following 293AAV triple transfection, conditioned medium is collected and replenished twice every 72 hours, as adapted and modified from Benskey et al., substantially increasing total viral yield [21].
2) **PEG-Mediated Precipitation**: The collected medium is filtered and then subjected to precipitation using polyethylene glycol (PEG) solution, following the ExtraPEG protocol, efficiently concentrating EVs and AAVs from large volumes while preserving their structural and biological integrity [22].
3) **Differential Centrifugation**: Precipitated material is clarified to remove insoluble aggregates, apoptotic bodies, and residual PEG, yielding a final preparation enriched in small EVs (90-200 nm), predominantly exosomes and microvesicles along with AAV particles (summarized in Fig. 1).
4) **Downstream application - *ex vivo* transduction of mouse/rat zygotes:** AAV-EV (AAV2/1 serotype) vectors enable convenient delivery of expression constructs or homology templates simply by adding the AAV-EV vectors directly to the embryo culture medium.

**Figure 1:**
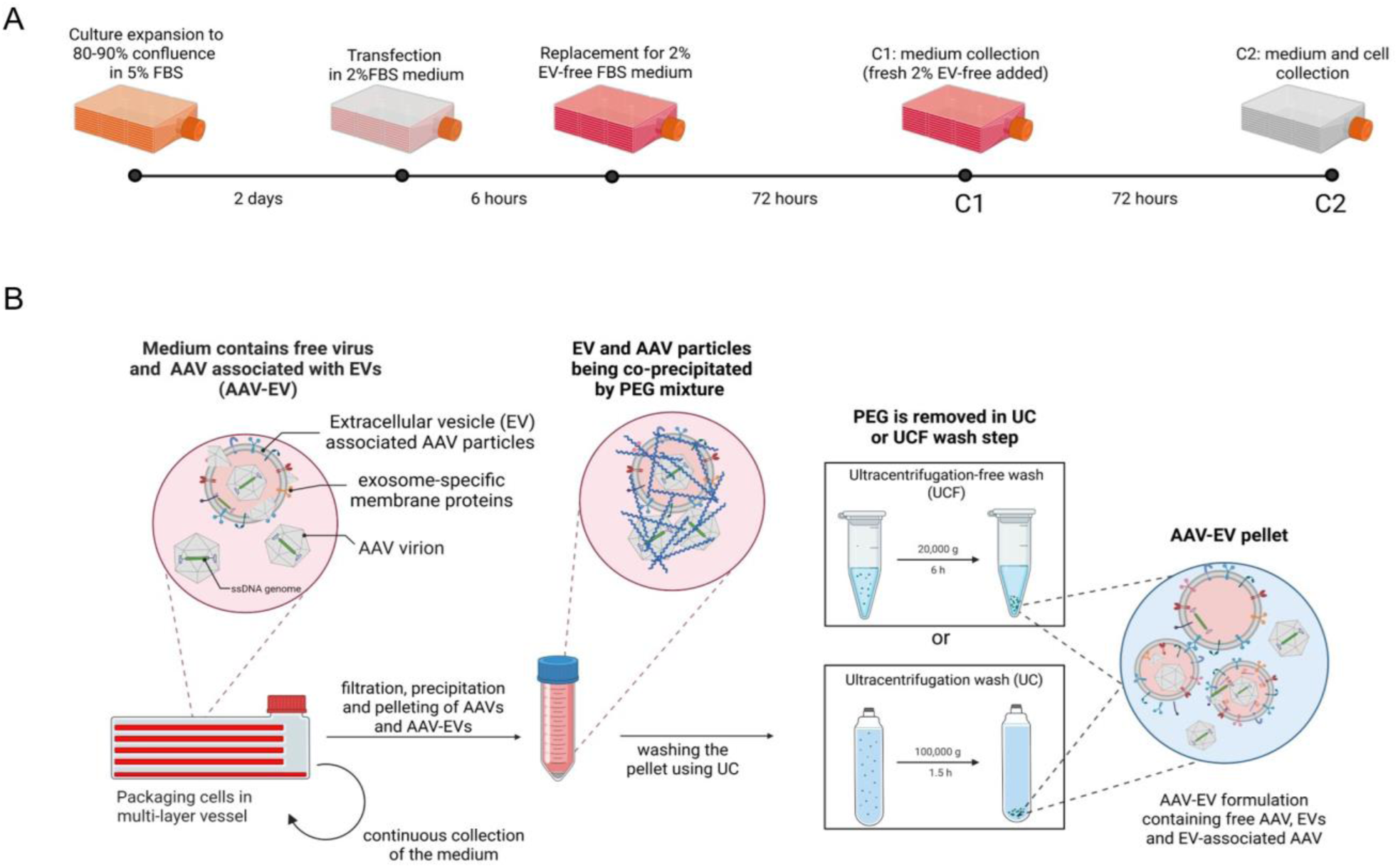
Production and isolation procedure. (A) Schematic description of AAV-EV production and the schedule of collections. (B) Washing step, including either ultracentrifugation (UC) or ultracentrifugation-free (UCF) pelleting of AAV-EVs.

This method bypasses density-gradient ultracentrifugation by using EVs as a biological matrix for direct pelleting of AAV2/1. This EV-based co-isolation also creates an ultracentrifugation-free workflow, improving the scalability and practicality of AAV production for applications such as *ex vivo* treatments and mouse or rat model generation.

## 2. Materials and Methods

### Cell culture

293AAV cell line (AAV-100) was purchased from Cell Biolabs, Inc. The 293AAV cells were cultured in high-glucose Dulbecco’s modified Eagle’s medium-DMEM (Sigma-Aldrich) supplemented with 5% fetal bovine serum-FBS (Gibco, certified, United States) and penicillin (10,000U/mL)/streptomycin (10,000 U/mL) solution (Gibco) in a humidified atmosphere with 5% CO_2_ at 37°C.

### Plasmids and cloning

HDR constructs, such as Actn1 cKO_HDR, Alox5ap-Ccre, Batf3-Cre-eGFP HDR, Lyve1-NCre-HDR, Nes-rtTA3-DTR-IRES-iRFP670-Far5, Dpp4-rtTA3-DTR-H1-mKate2, Ube3a-sl-BioID2_HDR, Alb-CYP3A4-CES1_HDR, Muc1-MUC1_wt, Muc1-MUC1_fs were synthetized by GenScript, and gene fragments were subcloned into AAV-CMVc-Cas9 backbone using *XbaI* and *NotI* restriction sites. AAV-CMVc-Cas9 was a gift from Juan Belmonte (Addgene plasmid # 106431; http://n2t.net/addgene:106431; RRID: Addgene_106431). HDR cassettes up to 2.5kb pAAV-Lck loxP_HDR, Umod C127R and Umod Y180_R188del were synthetized by GenScript, gene fragment was subcloned into pscAAV-Sox2-mStr using *XbaI* sites. pscAAV-Sox2-mStr was a gift from Lin He (Addgene plasmid # 135617; http://n2t.net/addgene:135617; RRID: Addgene_135617). pAAV-RC1 plasmid (VPK-421) and pHelper plasmid (part.no. 340202) were purchased from Cell Biolabs, Inc. PiggyBac system: Hypertransposase (Hyp, pAAV CMV-Hyp) and EGFP transposon (pAAV-CMV-EGFP) plasmids were assembled and provided by Nadav Ahituv and Aki Ushiki. Split-Cre system (NCre/CCre with SpyTag/SpyCatcher conjugation) used to generate pAAV Alox5ap-Ccre and Lyve1-NCre-HDR constructs was provided by Xuyu Zhou [23].

### Standard AAV production

Standard AAV was produced by triple transfection in 293AAV cells seeded in one Corning® HYPERFlask® Cell Culture Vessels (Corning). PEI-mediated transfection (PEI: 24765-1, Polysciences, Inc.) was performed in Dulbecco’s modified Eagle’s medium (Sigma-Aldrich) supplemented with 2% fetal bovine serum (FBS, Gibco, certified, United States) and penicillin (10,000 U/mL)/streptomycin (10,000 U/mL) solution (Gibco). Total amount of plasmid DNA per vessel was 550µg in equimolar ratio 1:1:1. Cells were then incubated overnight. PEI:DNA ratio was 3:1. Next day, the medium was changed for fresh DMEM (Sigma-Aldrich) supplemented with 2% FBS and penicillin/streptomycin solution (Gibco). After 4 days post transfection, the cells and medium were collected and processed as described elsewhere [8]. Standard AAV vectors were stored at -80 °C for long time storage or 4°C when used the next day for downstream experiments. AAV were kept in 0.001% Pluronic-F68 (24040032, Gibco) PBS.

### AAV-EV production and isolation

Production of AAV-EV particles in Corning® HYPERFlask® Cell Culture Vessels (Corning) was initiated by triple transfection in 293AAV cells with AAV Rep/Cap, transfer and pHelper plasmids. In this study, only AAV vectors of AAV2/1 serotype were produced. PEI-mediated transfection was performed in Dulbecco’s modified Eagle’s medium (Sigma-Aldrich) supplemented with 2% fetal bovine serum (Gibco, certified, United States) and penicillin (10,000U/mL)/streptomycin (10,000 U/mL) solution (Gibco). The total amount of plasmid DNA was 550 µg per HyperFlask (for triple transfection and transfection with additional pseudo-typing plasmid, e.g. mEmerald-CD9-10, #54029) in equimolar ratio 1:1:1 (or 1:1:1:1 - CD9 pseudo-typing) and incubated overnight. We used PEI:DNA ratio 3:1 (1650 µg of PEI for 550 µg of DNA). AAV-free EV were generated using equivalent amount of transfer plasmid (as in AAV-EV production) and mixed with PEI in 3:1 PEI:DNA ratio.

Next day, the medium was changed for DMEM (Sigma-Aldrich) supplemented with 2% EV-free FBS (more information below) and penicillin/streptomycin solution (Gibco) and cultured. After 72 hours, the growth medium was collected. The second collection was done in the same period in respect to the previous medium change (72 hours). The last collection included harvest of 293AAV cells from the vessel. The time scheme is graphically depicted in Fig. 1A.

Immediately after collection, harvested medium was processed by filtering through PES vacuum Filter 0.22μm (431097, Corning) to remove cells, larger cellular debris and apoptotic bodies. Filtered medium was mixed with 50% PEG6000 and 375 mM NaCl solution to obtain solution of 10% PEG6000 and final concentration of 75mM NaCl (NaCl in DMEM was not considered).

Solution was mixed gently by inverting the bottle and swirling. Once the solution was properly mixed, samples were stored at 4°C up to 18 hours (overnight).

Next day, medium was centrifuged in a swing-bucket rotor at 3,200 x g (maximum speed) and 4°C for 90 min. Supernatant was discarded and pellet was resuspended in 10 mL of PBS-HT (Trehalose 25 mM; HEPES 25 mM in PBS, no human albumin-HSA) and stored at -80°C. Once all harvests were collected and processed by precipitation, harvests after precipitation (2x10 mL) were pooled and mixed with 45 mL of PBS-HT.

To remove residual cell debris, apoptotic bodies and larger vesicles, the supernatant was centrifuged at 21,000 x g and 4°C for 30 min using fixed-angle rotor (Eppendorf, 5810R). To pellet exosomes, microvesicles and AAV, the supernatant from the previous centrifugation was subjected to 90 min centrifugation at 100,000 x g at 10 °C using Ti70 in an Optima L-90K ultracentrifuge (Beckman Coulter, Indianapolis IN, USA). The AAV-EV pellets of collection 1 and 2 were pooled and resuspended in 0.2-0.5 mL of PBS-HAT (human albumin-HSA 0.2 %; Trehalose 25 mM; HEPES 25 mM in PBS) as described by [24]. Isolates were aliquoted and stored at -80°C.

**EV-free 50% FBS** was prepared by mixing FBS and DMEM medium in 1:1 ratio to make 50% FBS. This solution was ultracentrifuged overnight at 100,000 x g and 4°C, adopted from Saari et al. [25]. Supernatant was collected and sterilized by filtration through 0.22µm syringe filter, then aliquoted and kept at -20°C.

AAV-EV were stored at -80 °C for long time storage or 4°C and used the next day for other experiments. Production of AAV-EVs carrying HDR template was performed with a single medium collection. Average titer yield 1 x 10^12^ GC (genome copies)/mL is often sufficient for downstream treatment of zygotes *ex vivo*.

For **ultracentrifugation-free** (**UCF**, Fig. 1B) protocol, resuspended (in 10mL of PBS-HT) PEG precipitates of each collection (C1 and C2) were pooled and subjected to centrifugation at 21,000 x g and 4°C for 30 min. The supernatant was aliquoted into 1.5 mL tubes (0.5 mL per tube) and centrifuged at 21,000 x g for 6 hours and 10°C. The pellets were resuspended in 10 aliquots by 15 µL of PBS-HAT (in total 150 µL) and stored at -80°C.

**50% Polyethylene glycol 6000, 375mM NaCl solution** was prepared by dissolving Polyethylene Glycol 6000 (Fisher Chemical) and NaCl (S5886, Merck) powder in sterile-deionized water in w/v ratio. Solution was dissolved after constant stirring and elevated temperature (75-90°C) for approx. 4 hours. The solution was filtered through Vacuum Filter 0.22 μm (431097, Corning) and stored at 4 °C.

### Nanoparticle Tracking Analysis (NTA)

Concentrations and size of particles were determined using Nanosight LM-14 instrument equipped with 405 nm (Blue405), 60 mW laser (Nanosight, Salisbury, Great Britain) and sCMOS camera (Hamamatsu Photonics K.K., Hamamatsu, Japan). The camera level was set to 12. The measurement time was 60s in 5 repetitions. The data was processed using NanoSight NTA software v3.0., with detection threshold adjusted to 5. Samples containing AAV were inactivated by 10 min of UV light exposure prior to measurement.

### Liquid chromatography-Mass spectrometry (LC-MS)

Each analyzed formulation was measured in triplicate. LC separation was performed on Dionex Ultimate 3000 nano HPLC system connected with MS instrument. Samples were loaded onto the trap column (C18 PepMap100, 5 μm particle size, 300 μm x 5 mm, Thermo Scientific) for 2 min at 17.5 μL/min. Loading buffer was composed of water, 2% acetonitrile and 0.1% trifluoroacetic acid. Peptides were eluted with Mobile phase B gradient from 4.0% to 35.0% B in 64.0 min. Mobile phase buffer A was composed of water and 0.1% formic acid. Mobile phase B was composed of acetonitrile and 0.1% formic acid. Nano reversed phase column (Aurora Ultimate TS, 25 cm x 75 μm ID, 1.7 μm particle size, Ion Opticks) was used for LC/MS analysis.

Peptide mixture was analyzed on Thermo Scientific Orbitrap Ascend by data dependent approach. Eluting peptide cations were converted to gas-phase ions by electrospray ionization in Positive mode. Spray voltage was set to 1600 V and ion transfer tube temperature to 275 °C. MS1 scans of peptide precursors were analyzed in Orbitrap in range 350-1400 m/z at 120K resolution and with following settings: RF Lens 60%, maximum injection time 251 ms, AGC target 250. Only the precursors with charges states 2-7 and intensity higher than 3000 were selected for fragmentation. Dynamic exclusion duration was set to 60 s with 10 ppm tolerance. Peptide precursors were isolated by Quadrupole with 0.7 m/z isolation window and fragmented by CID with 30% normalized collision energy. Fragment ions were detected in Ion Trap with AGC target set to 100 and maximum injection time 100 ms.

### Statistical analysis

Proteomics data were imported from MaxQuant/Perseus output tables in Excel (Additional file 2). For each comparison (AAV-EV-EGFP vs. EV-EGFP, AAV-EV-HYP vs. EV-HYP, AAV-EV-Sox2mStr vs. EV-Sox2mStr), relevant columns (Protein names, Gene names, Peptides, Student’s T-test Difference, and -Log Student’s T-test p-value) were parsed. Proteins supported by ≤ 1 peptide were excluded. p-values were reconstructed from -log10(p) and adjusted for multiple testing using the Benjamini-Hochberg procedure to obtain FDR-corrected q-values. Proteins were considered significant if |log2FC| ≥ 1 and q ≤ 0.05.

Volcano plots were generated with log2FC on the x-axis and -log10(q) on the y-axis, with thresholds indicated at |log2FC| = 1 and q = 0.05. Significant proteins were colored red (up in AAV-EV) or blue (down in AAV-EV, up in EV), and non-significant (below thresholds) proteins were labeled gray. The 25 proteins with the lowest q-values were highlighted with a red ring. Marker sets (exosome/microvesicle-specific, exosome/microvesicle-negative, and AAV-specific proteins including Hyp-Hypertransposase) were annotated using name-based matching. Labels for these and Top-25 proteins were placed using a non-overlapping algorithm to ensure clarity. All processing and visualization were performed in Python using pandas and matplotlib.

The major EV and AAV proteins were highlighted in the plot, to emphasize their abundance:

exosome/microvesicle-specific markers: CD63 antigen CD63), Tumor susceptibility gene 101 (TSG101), CD9 antigen (CD9), CD81 antigen (CD81), Flotillin-1, Heat shock protein HSP 90-alpha (HSP90α), GAPDH, Programmed cell death 6-interacting protein, Syntenin-1.

exosome/microvesicle-negative markers: Calnexin (CALX), Calreticulin (CALR) Heat shock protein HSP 90-beta (HSP90β), Heat shock 70 kDa protein 1A (HSPA5), histones H2A, H2AFY2 a H3F3A, GM130, Cytochrome C, β-actin AAV-specific proteins: AAV1 VP1, AAV1 VP2, AAV VP3, E1A, E1B, E1B19K, E1B55K, Rep (Rep78/68/55/40), DBP, Hyp.

**EGFP/Rep/DBP expression and copy number in E15.5 embryos and placentas** was determined by quantitative PCR using the ΔΔCt method with Ubb as an endogenous reference. For each sample, Ct values from technical triplicates were averaged. The normalized Ct value (ΔCt) was calculated as the difference between the mean target gene Ct and the mean Ubb Ct for the same sample. A genomic DNA or cDNA (in case of RNA expression analysis) reference sample with a defined copy number was used as the calibrator/positive RNA reference, and ΔΔCt was calculated as the difference between the sample ΔCt and the reference ΔCt. Relative copy number/expression was then calculated as 2^−ΔΔCt2^, assuming equal and near 100% PCR efficiencies for target genes and Ubb assays. Error propagation was performed based on the standard deviation of Ct values.

#### Statistical analysis of PiggyBac experiments

Copy-number analyses used per-sample “Average Copy Number” values from Additional file 5; positive/negative controls (PC/NC) were excluded. Embryo-placenta pairs were matched by shared numeric keys in sample IDs (e.g., 15E_61 ↔ 15P_61). At the matched dose, embryo copy-number distributions for AAV-EV versus AAV were compared using the Wilcoxon-Mann-Whitney test (two-sided). Differences across embryo cohorts (AAV-EV matched, AAV matched, AAV-EV high dose) were assessed by a Kruskal-Wallis test. Within-animal placenta-embryo differences were tested using a paired Wilcoxon signed-rank test with the directional alternative *embryo > placenta*. For each nonparametric test, we report the exact p-value (or normal approximation when ties/zeros precluded an exact calculation) and a rank-based effect size: rank-biserial correlation for between-group Mann-Whitney and matched-pairs rank-biserial for paired Wilcoxon. Viability at the matched dose (AAV vs AAV-EV) was evaluated by Fisher’s exact test on the counts reported in the study (AAV 16/158; AAV-EV 36/152); we additionally provide relative risk (RR) with a Wald 95% confidence interval on the log scale. All analyses were performed in Python (pandas/scipy), reproducing the cohort means/medians shown in the Supplementary tables (Additional files 5 and 6).

### Zygote preparation

Mouse: 5-8 weeks old female C57BL/6NCrl mice (Crl:SD, Charles River) were super-ovulated by intraperitoneal application of 5 IU/female of pregnant mare serum gonadotropin (Calbiochem, Millipore, 367222), and 46-48 h later, 5 IU/female human chorion gonadotropin (Calbiochem, Millipore, 230734). Super-ovulated females were mated one-to-one with 10-25 weeks old C57BL/6NCrl males to produce one-cell zygotes at 0.5 days post-coitum. Zygotes were collected and washed. In short, zygotes were harvested from the ampulla of euthanized females, washed in hyaluronidase/M2 solution (Millipore, MR-051-F) to remove cumulus cells, washed four times in M2 media (Zenith, ZFM2-100) supplemented with 4 mg/mL bovine serum albumin (BSA, Sigma, A3311), and washed again four times in M2 + BSA media, eventually zygotes were transferred to KSOM + AA media (KCl-enriched simplex optimization medium with amino acid supplement, Zenith Biotech, ZEKS-050).

Rat: 6-8 weeks old female Sprague Dawley rats (Charles River) were super-ovulated by intraperitoneal application of 300 IU/kg of pregnant mare serum gonadotropin (Calbiochem, Millipore, 367222), and 46-48 h later, 200 IU/kg human chorion gonadotropin (Calbiochem, Millipore, 230734). Super-ovulated females were mated one-to-one with 10-25 weeks old SD males. Next day, females with a vaginal plug were selected for one-cell zygotes extraction in the morning upcoming day. Zygotes were collected and washed as follows. In short, zygotes were harvested from the ampulla of euthanized females, washed in hyaluronidase/M2 solution (Millipore, MR-051-F) to remove cumulus cells, washed four times in M2 media (Zenith, ZFM2-100) supplemented with 4 mg/mL bovine serum albumin (BSA, Sigma, A3311), and washed again four times in M2 + BSA media, eventually zygotes were transferred to KSOM + AA media (Zenith Biotech, ZEKS-050).

### Zygote treatment with AAV/AAV-EV vectors

Both mouse and rat zygotes were cultured in 50 µL droplets of KSOM + AA media (Zenith Biotech, ZEKS-050) containing corresponding AAV-EV vector dose per 50 µL drop (in high, medium or low virus dose, 2.5 x 10^10^ , 5 x 10^9^ and 2.5 x 10^9^ GC, respectively) in 35 x 10 mm culture dishes (CellStar Greiner Bio-One, 627160) at 37 °C with 95% humidity and 5% CO_2_ for 4 hours prior to RNP electroporation. AAV-EV-treated embryos were electroporated with assembled RNPs as previously described [2]. Briefly, embryos were transferred to 10 µL of Opti-MEM reduced serum media (Thermo Fisher, 31985062) with assembled Cas9/sgRNA RNPs. Embryos in RNP mixture were transferred to a 20 mm length platinum plate electrode on glass slide, 1 mm gap, 1.5 mm height (Bulldog-Bio, CUY501P1-1.5) and electroporated (poring: 40V, pulse length - 3.5ms, interval - 50ms, number of pulses - 4, decay - 10%, + polarity; transfer: 5 V wave, pulse length - 50ms, interval - 50ms, number of pulses - 5, decay - 40%, ± polarity) using a NEPA21 (Nepagene). Zygotes were recovered from the cuvette by flushing three times with 100 µL of KSOM + AA media, then transferred back into the culture droplets containing AAV-EV for a total incubation length of 24 hours. The following day, embryos were transferred to fresh KSOM + AA media overlaid with mineral oil until analysis or oviduct transfer. For generation of live mice, treated zygotes, which successfully developed into two-cell embryos, were washed by three flushes in 100 µL of KSOM + AA media and surgically transferred into the oviducts of pseudo-pregnant ICR:CD1 females (Charles River, Strain 022) or SD rat females (Charles River), using up to 18 embryos per oviduct. Used gRNAs are summarized in Additional file 8 table.

### Zygote treatment with PiggyBac system

Zygotes were cultured in 30 µL droplets of KSOM + AA media (KCl-enriched simplex optimization medium with amino acid supplement, Zenith Biotech, ZEKS-050) containing corresponding AAV-EV or AAV vectors titer in 35 x 10 mm culture dishes (CellStar Greiner Bio-One, 627160) at 37 °C with 95% humidity and 5% CO_2_ for 24 hours. Treatment with PiggyBac system consists of Hypertransposase and EGFP transposon. Embryos were always treated with both constructs in either AAV-EV or AAV form. For the treatment to compare AAV-EV and AAV, the following doses per 30 µL droplet were used - Hyp: 3 x 10^9^ GC; EGFP transposon: 1.5 x 10^10^ GC. The high-dose AAV-EV treatment was performed with the following titer per 30µL droplet - EGFP transposon: 1.5 x 10^11^ GC; Hyp: 3 x 10^10^ GC. The following day, embryos were transferred to fresh KSOM + AA media overlaid with mineral oil until oviduct transfer. For generation of live mice/embryos, treated zygotes, that successfully developed into two-cell embryos, were washed by three flushes in 100 µL of KSOM + AA media and surgically transferred into the oviducts of pseudo-pregnant ICR:CD1 females (Charles River, Strain 022), using up to 18 embryos per oviduct.

### Fluorescence imaging of blastocysts and E15.5 embryos

Blastocysts were captured with Zeiss Axio Zoom V.16 microscope (2000 ms exposure), using a 560-585 nm excitation wavelength and a 600-690 nm emission wavelength, at a total magnification of 12.5x. E15.5 embryos were captured in PBS with an exposure time of 180 ms, using a 480 nm excitation wavelength and a 509 nm emission wavelength, at a total magnification of 11.2x. All images were processed in ZEN 3.0 (blue edition) software.

### Genotyping

Mice born after AAV and AAV-EV zygote treatment were genotyped 4 weeks after birth.

Tissue samples (tail/ear) of all mentioned strains were lysed in lysis buffer (100mM Tris HCl pH 8, 200mM NaCl, 5mM EDTA, 10% SDS) with proteinase K (R75282, P-LAB) overnight at 55 °C, then inactivated at 95 °C for 15 min. Lysates were further processed by phenol: chloroform based isolation using Phenol/Chloroform/Isoamylalcohol reagent (A156.2, ROTI) following the manufacturer’s protocol.

PCR reaction was performed using DreamTaq™ Green Buffer (10X, Thermo Scientific), DreamTaq DNA Polymerase (5 U/µL, Thermo Scientific) and Deoxynucleotide Set, 100 mM (Sigma Aldrich). Genotyping was performed with the primers listed in Fig. S4, to confirm a correct genomic insertion.

Genotyping of animals treated with Lck cKO: loxP, Umod C127R and Umod Y180_R188del HDR AAV-EV vectors detection was performed using digestion of PCR product with a corresponding restriction enzyme (Fig. Additional file 8 figure C, L, M)

### DNA and RNA isolation from E15.5 embryos

ICR:CD1 surrogate females were euthanized by CO_2_ inhalation and cervical dislocation to collect E15.5 embryos. The embryos were halved along sagittal axis, where one half of the embryo was used for DNA isolation, and the second half for RNA isolation and reverse transcription.

DNA isolation was performed by lysis in lysis buffer (50mM Tris HCl pH 8, 1mM EDTA, 0,5% Triton X-100) with proteinase K (R75282, P-LAB) overnight at 55 °C. The lysate was treated by RNase A (Sigma-Aldrich R4875-1G) and processed by phenol: chloroform-based isolation using Phenol/Chloroform/Isoamylalcohol reagent (A156.2, ROTI) following the manufacturer’s protocol. DNA was further used for qPCR and copy number variation analysis using DNA from homozygote Rosa26-VFRL-EGFP (MuX) reporter mouse as a positive reference.

RNA isolation was performed using TRI Reagent® (Sigma Aldrich). Isolated RNA was treated with *DNaseI* (Life Technologies, AM2222). RNA was further reverse-transcribed into cDNA using reverse transcriptase (M-MLV Reverse Transcriptase, Promega, M1705) and used for qRT-PCR.

### qPCR and qRT-PCR

AAV titer was quantified by real-time qPCR with LightCycler® 480 SYBR Green I Master Mix, primers amplifying *AAV2 ITR* sequence: forward 5’-GGAACCCCTAGTGATGGAGTT, reverse 5’-CGGCCTCAGTGAGCGA [26] and related software (Roche). Before each quantification, DNA samples were isolated and purified using High Pure Viral Nucleic Acid Kit (Roche).

The qPCR and qRT-PCR reaction was performed with LightCycler® 480 SYBR Green I Master Mix and gene-specific primers amplifying *EGFP* sequence in CMV-EGFP transposon: forward 5’-ACGACGGCAACTACAAGACC, reverse 5’-TGTAGTTGTACTCCAGCTTGTGC, and reference gene Ubiquitin B (Ubb) primers: forward 5’-ATGTGAAGGCCAAGATCCAG, reverse 5’-TAATAGCCACCCCTCAGACG for normalization by Pfaffl [27]. For replication assay, the following primers were used to detect DNA/RNA of: AAV Rep2 gene - forward: 5’-TCACCAAGCAGGAAGTCAAAG, reverse: 5’-CCCGTTTGGGCTCACTTATATC, Adenoviral DNA binding protein (DBP): forward 5’-CACTGGGTCGTCTTCATTCA, reverse: 5’-CGCTACAAATGGTGGGTTTC. pAAV-RC1 plasmid (VPK-421, CellBiolabs) and pHelper plasmid (part.no. 340202, CellBiolabs) were used as standards quantification of DNA/RNA presence in embryo and as positive controls for the primers.

### Continuous density gradient (CDG) separation

The density gradient was prepared with the diffusion method described by Saari et al. [25] . Briefly, the gradient was prepared in a 13.5 mL ultracentrifuge tube (344085, Beckman Coulter) by overlaying 4 mL of 45 % iodixanol with 4 mL of 25 % iodixanol and filling the rest of the tube with DPBS-2 mM MgCl_2_. The tubes were sealed with parafilm, carefully tipped to a horizontal position and incubated for one hour at RT. After the incubation, the tubes were chilled on ice and 1.5 mL were removed from the top to make room for sample loading. Each sample was separated and analyzed in triplicate. The samples (1 x 10^10^ GC per tube) mixed with 60% iodixanol (07820, Stemcell technologies) were then loaded through the gradient to the bottom of the tube using a syringe with long needle and ultracentrifuged at 200 000 x g at 4°C for 3 hours using rotor SW41Ti (Beckman Coulter). After the run 1 mL fractions were collected from the top of the tubes for further analysis. Density of individual fractions was measured by OptiPrep Application Sheet V05 (https://diagnostic.serumwerk.com/wp-content/uploads/2021/05/V05-Serumwerk.pdf). AAV genomes were quantified in each fraction using AAV2 ITR specific primers (Additional file 4).

### Ethics statement

Mice were bred in our specific pathogen-free facility (Institute of Molecular Genetics of the Czech Academy of Sciences; IMG). This study was conducted in accordance with the ARRIVE guidelines and the laws of the Czech Republic. Animal protocols (93/2020 and 101/2020) were approved by the Resort Professional Commission for Approval of Projects of Experiments on Animals of the Czech Academy of Sciences, Czech Republic.

All mice were euthanized using carbon dioxide (CO_2_) inhalation followed by cervical dislocation. CO_2_ inhalation was performed at a flow rate that displaced 30% of the chamber volume per minute to minimize distress.

## 3. Results

### Co-Isolation of AAV Vectors for AAV-EV Formulations

The validation of AAV isolation protocol by AAV producer cell line for EV production was performed in a multilayer vessel. The protocol is based on continuous collection of growth medium combined with PEG precipitation. That yields a mixture of free and EV-associated AAVs, referred here as the AAV-EVs. This approach enabled the isolation of AAVs from growth medium without density gradient separation. The obtained formulations using ultracentrifugation-based protocol are marked “UC”. The collection scheme is depicted in Fig. 1A and wash step, in Fig. 1B.

The AAV-EV vectors were pseudo-typed by co-transfecting CD9-mEmerald during production as a reference for enrichment of individual (C1 and C2) and pooled collections after UC isolation. CD9-mEmerald enabled exclusive labeling of exosomes and microvesicles. A green pellet indicated enrichment by labeled exosomes and microvesicle in the formulation (Additional file 2).

To assess the robustness of EV-based AAV isolation, we subjected PEG-precipitated samples from two pooled collections (Fig. 1A) to differential centrifugation, concluding with a 6-hour spin at 21,000 x g at 10 °C (Fig. 1B). We termed this alternative approach ultracentrifugation-free protocol (UCF). This procedure eliminated the need for standard ultracentrifugation while maintaining pelleting efficiency and AAV yield. This method has produced AAV-EV-Sox2-mStr and AAV-EV-Alb-CYP3A4-CES1, and AAV-EV-Umod C127R targeting vectors. The production workflow is schematically illustrated in Fig. 1A and B, with double collection of growth medium performed at 72-hours intervals.

The formulations used for delivery of homology-directed repair template (HDR) and expression constructs (EXP) *ex vivo* were not pseudo-typed with CD9mE to avoid the risk of random integration of carryover CD9mE plasmid or its fragments. The key characteristics of all AAV-EV formulations used in this study, including AAV titer, EV particle count, and nanoparticle size parameters (mode and mean size), are summarized in Table.1. Nanoparticle tracking analysis (NTA) showed that AAV-EV formulations were predominantly composed of 100-150 nm particles, a size range consistent with exosomes and small microvesicles.

The differences and complexity of protein compositions were analyzed by Liquid chromatography-Mass spectrometry (LC-MS) using the three main formulations. These were the HDR vector Sox2-mStrawberry targets *Sox2* gene, which replaces stop codon with P2A-mStrawberry sequence, and allows tracking of *Sox2* expression (Fig. 2A). The other two expression vectors carry the EGFP transposon and Hypertransposase (Hyp), Fig. 2B and C. For each construct, the protein composition of AAV-EV co-isolated vectors was compared with that of EV preparations generated in the absence of AAV packaging plasmids, produced using the corresponding transfer plasmids alone (pAAV-Sox2-mStr, pAAV-CMV-EGFP, and pAAV-CMV-Hyp).

**Figure 2:**
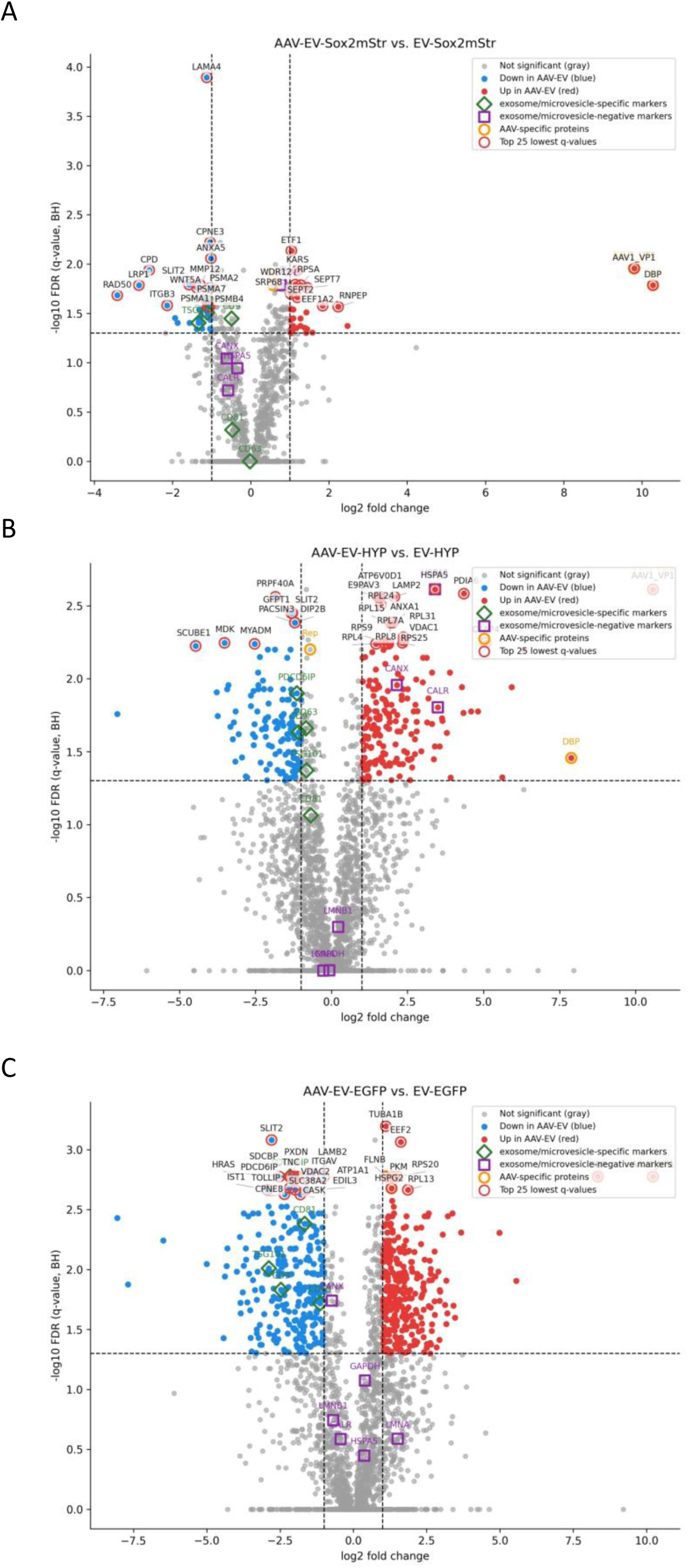
Comparative proteomic analysis of AAV-EV and EV vectors. Negative (left) side of X axis in the plot represents proteins abundant in EV reference. Positive (right) side of X axis in the plot represents proteins abundant in AAV-EV vectors. Data points above Y axis threshold represent proteins with q ≤ 0.05 (FDR-adjusted). (A) Comparative proteomic analysis of AAV-EV (AAV+) and EV (AAV-) carrying HDR template (Sox2-mStr). (B) Comparative proteomic analysis of AAV-EV (AAV+) and EV (AAV-) carrying EGFP transposon. (C) Comparative proteomic analysis of AAV-EV (AAV+) and EV (AAV-) carrying Hypertransposase. *AAV+ = adeno-associated virus positive AAV- = adeno-associated virus negative, FDR*-*adjusted p*-*value: expected false*-*positive proportion among significant findings, False Discovery Rate (FDR)*

LC-MS profiling revealed a modest enrichment of canonical exosome markers (Syntenin-1/SDCBP, ALIX/PDCD6IP, CD9, TSG101) in virus-free EV preparations, confirming their vesicular identity. In contrast, core exosome/microvesicle markers (TSG101, CD9, FLOT1, CD81) were moderately reduced in AAV-EV samples relative to EV-only controls, indicating that AAV production alters EV protein composition and favors incorporation of viral components (Fig. 2A-C). Enrichment of EV-negative or apoptotic-body-associated markers was formulation-dependent and most pronounced in AAV-EV-Hyp preparations (e.g., Calnexin, Calreticulin), whereas Sox2-mStrawberry and EGFP transposon vectors showed only modest or non-significant changes at the predefined thresholds.

Packaging-related proteins (AAV Rep78/68/55/40; adenoviral DBP/E1B) were detectable in AAV-EV proteomes, consistent with helper-free vector production; however, neither Rep nor DBP DNA or RNA was detected in E15.5 embryos (Additional file 7), arguing against *in vivo* viral replication under these conditions. Together, these data indicate a compositional shift in AAV-EV formulations characterized by enrichment of viral proteins at the expense of exosome-associated markers, motivating further evaluation of AAV-EV properties with respect to safety and functional performance in downstream applications.

Cryo-electron microscopy (Cryo-EM) of AAV-EV and AAV samples confirmed the presence of AAV virions (∼25 nm) and larger microvesicles (80-400 nm, Additional file 3). To evaluate AAV and EV associations, AAV-EV formulations were fractionated using iodixanol density gradients (Fig. 3), which separated the particles by their density. AAV-EV preparations obtained by ultracentrifugation (UC) and ultracentrifugation-free (UCF) pelleting exhibited comparable viral titers, EV counts, and particle size distributions (Table.1). Both formulations showed enrichment of AAV genomes in 1.04-1.21 g/cm^3^ fractions, suggesting partial association with the vesicles. Quantitatively, 16.45 % (UC) and 33.55% (UCF) of total AAV genomes were localized in these fractions (Fig. 3). All the tested isolates show the majority of AAV genomes in 1.23-1.29 g/cm^3^ (40-50% iodixanol) fractions, corresponding to free AAV virions or small, dense vesicles. These findings indicated that while most AAV particles remain unassociated, a measurable proportion is retained in lower-density fractions due to their vesicular associations. Notably, unassociated AAVs were co-pelleted with EVs, supporting a role for EVs as a physical scaffold facilitating AAV recovery during the pelleting process. The necessity of separating EVs from AAV, or isolating a strictly EV-associated AAV population, was tested using PiggyBac transposon system *ex vivo*.

**Figure 3:**
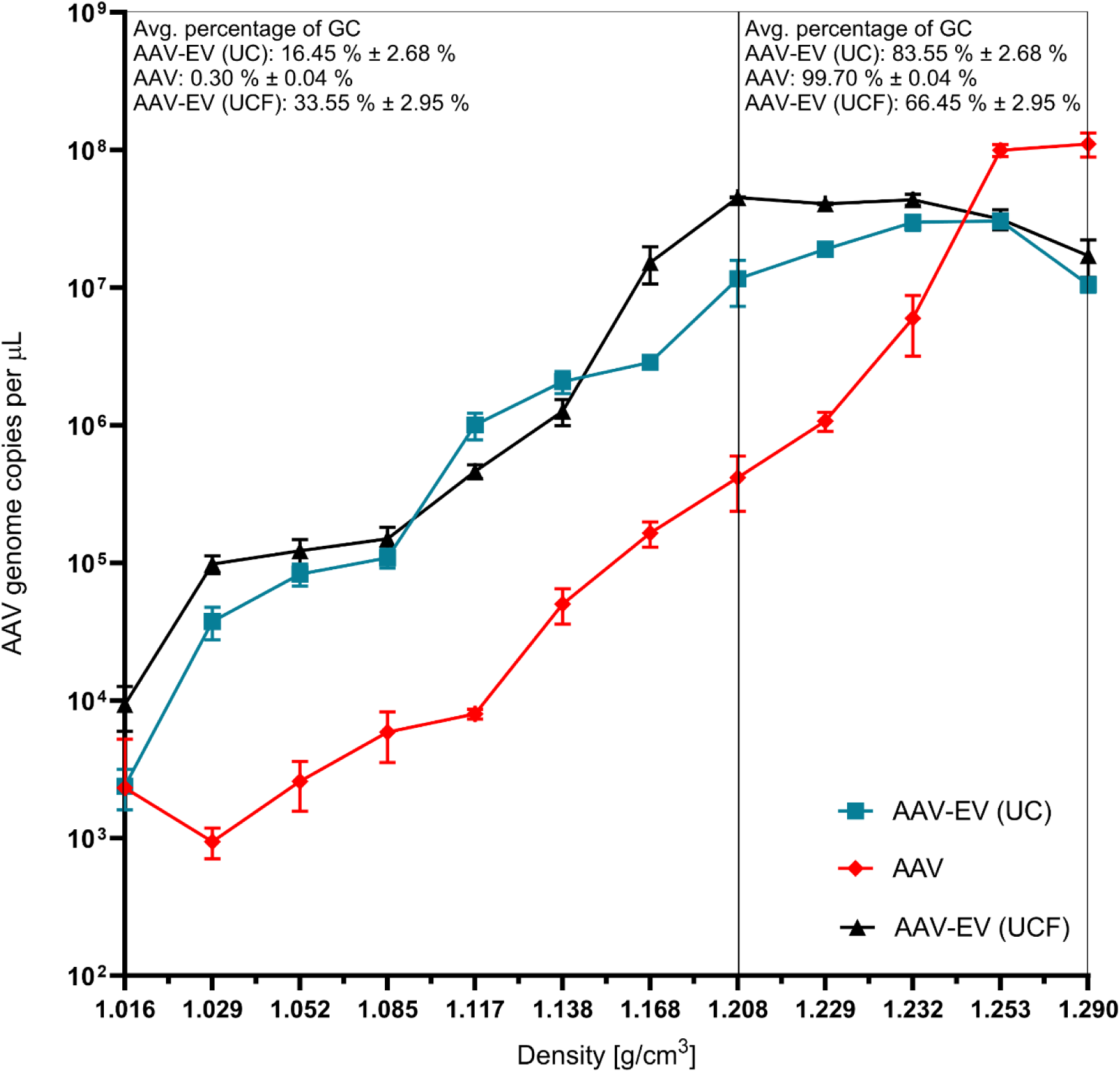
Quantification of AAV genomes in distinct fractions of iodixanol CDG of AAV and AAV-EV of serotype AAV2/1. CDG separates particles based on density: free AAVs in 1.21-1.29 g/cm^3^ (right side), EVs in 1.08-1.19 g/cm^3^ (left side). Fractions ≥1.21 g/cm^3^ contain free-AAV and high density EVs.

### AAV-EV-Mediated PiggyBac Delivery Enables Efficient Integration without Loss of Embryonic Viability and Viral Replication Risk

Delivery potential of AAV-EV and AAV vectors was tested by employing PiggyBac (PB) transposon system to deliver the constitutive CMV-EGFP reporter (EGFP transposon) into the mouse genome at the highest possible copy number. The constitutive system was utilized to evaluate AAV-EV and AAV delivery efficiency and toxicity. The efficiency was subsequently assessed by number of integration events of EGFP transposon per genome and number of viable embryos. Additionally, we assessed EGFP copy number in the placenta, hypothesizing that embryo might sequester some portion of the vector into this extra-embryonic protective tissue (Fig. 4A and B).

**Figure 4:**
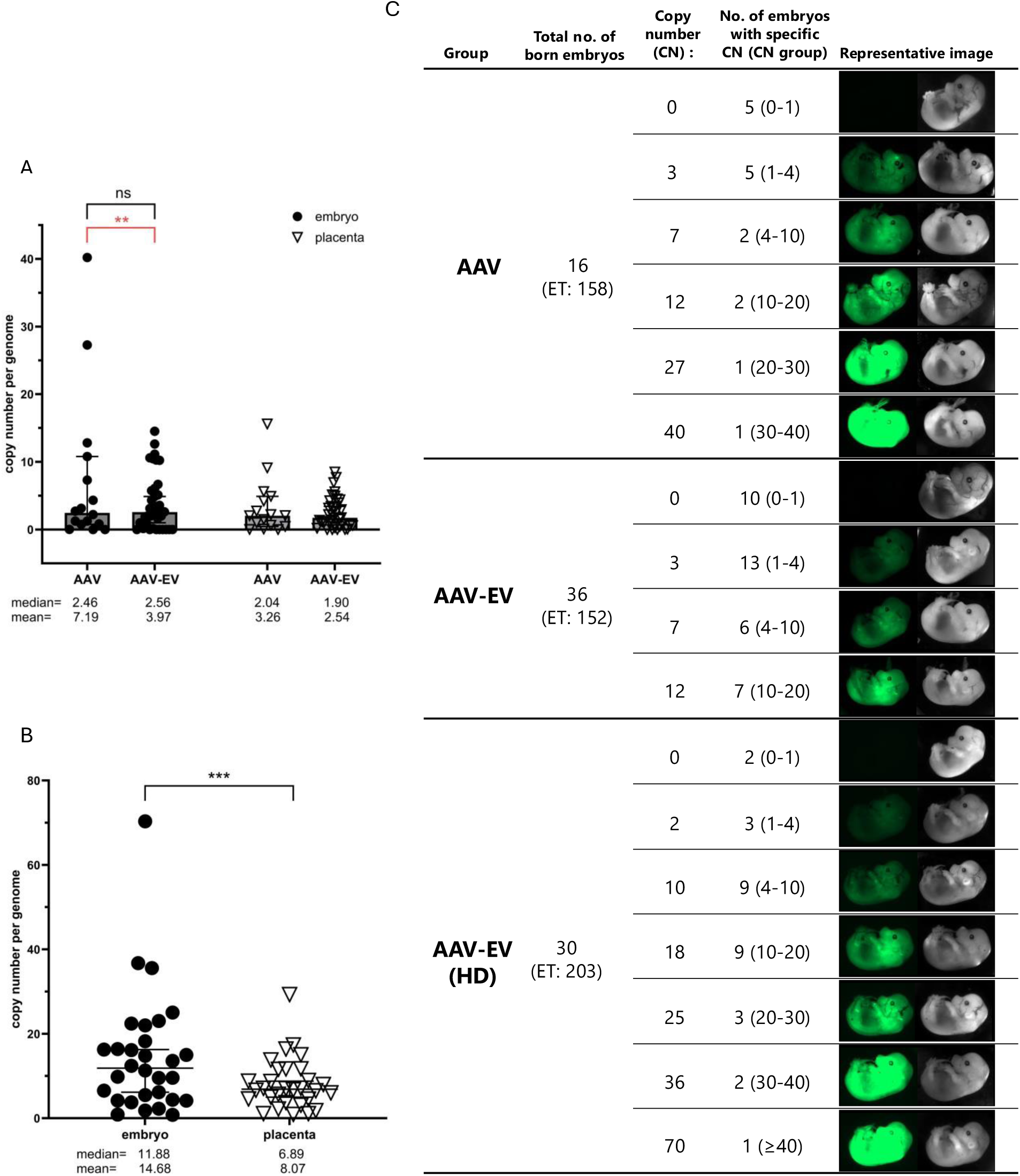
Delivery efficiency and toxicity of PiggyBac using AAV versus AAV-EV. (A)Copies of the EGFP transposon per genome in embryos and placentas after *ex vivo* zygote treatment with AAV or AAV-EV PiggyBac vectors at matched titers (Hyp: 3 x 10^9^ GC; EGFP transposon: 1.5 x 10^10^ GC), together with counts of viable embryos. Viability (red) (AAV-EV vs AAV): Fisher p = 0.0021; RR = 2.34 [95% CI 1.36-4.03] (36/152 vs 16/158), EGFP copy number distribution (between-group embryo comparison, black): Mann-Whitney U = 266.5, p = 0.687; r_rb = 0.075. (B) High-dose AAV-EV cohort (EGFP transposon: 1.5 x 10^11^ GC; Hyp: 3 x 10^10^ GC): copies per genome in embryo and placenta and corresponding viability. Paired Wilcoxon (embryo > placenta): p = 1.34 x 10^-5^; r_rb = 0.81 (C) Relationship between EGFP fluorescence and copies per genome (E15.5) with the distribution of embryos across predefined copy-number (CN) groups (0-1, 1-4, 4-10, 10-20, 20-30, 30-40, ≥70). Abbreviations: CNA, copy-number analysis; ET, total transferred 2-cell embryos; HD, high dose. Statistical tests are described in Methods.

At the matched dose, AAV-EV and AAV yielded comparable embryo EGFP copy-number distributions (medians 2.56 and 2.46, respectively), with both medians exceeding 1 (Fig. 4A). Within each vector condition, embryo copies exceeded placental copies (paired Wilcoxon: AAV-EV: W = 446.0, p = 0.00155, r_rb = 0.590; AAV: W = 109.0, p = 0.0168, r_rb = 0.603), consistent with higher transduction in embryos than matched placentas at this dose (Fig. 4A). In the same matched-dose experiments, viability favored AAV-EV over AAV (36/152 vs 16/158). Escalating to high-dose AAV-EV further increased copy numbers (embryos: mean 14.68, median 11.88; placentas: mean 8.07, median 6.89), with a large embryo-over-placenta shift (W = 421.0, p = 1.34 x 10^-5^, r_rb = 0.811) while maintaining viability (Fig. 4B).

Across cohorts (AAV-EV matched, AAV matched, AAV-EV high-dose), copy-number distributions differed (Kruskal-Wallis H = 20.29, p = 3.93 x 10^-5^), consistent with a dose-related increase in the high-dose AAV-EV group (Fig. 4A-C, Additional file 5). Full test specifications (viability: Fisher; between-group copies: Wilcoxon/Mann-Whitney; cohort trend: Kruskal-Wallis; placenta-embryo: paired Wilcoxon) and all effect sizes and exact p-values are provided in Methods and Additional file To determine whether the DNA-binding protein and AAV Rep proteins originated from active expression of carryover molecules, total DNA and RNA were isolated from E15.5 embryos treated with high-dose AAV-EV carrying PB vectors (Additional file 7). Samples were analyzed by qPCR and qRT-PCR to quantify integrated EGFP transposon copies, EGFP mRNA expression, and the presence of Rep and DBP sequences from the helper-free packaging system. No evidence of vector replication or detectable DNA/RNA signals for Rep and DBP was found in highly EGFP-positive embryos (Additional file 7), indicating these proteins do not persist in treated animals.

AAV-EV vectors demonstrated efficient PiggyBac-mediated transgene integration while preserving embryonic viability, outperforming conventional AAV in toxicity profiles. High-dose AAV-EV treatments achieved elevated copy numbers per genome without compromising survival, highlighting their potential for advanced genetic engineering. No evidence of residual viral replication was detected in treated embryos, confirming the safety of this approach.

### AAV-EV Enables Efficient Delivery of HDR Templates *Ex Vivo*

To evaluate the capacity of AAV-EV vectors for precise genome editing, we applied them to deliver Homology Directed Repair (HDR) templates into mouse and rat zygotes. This approach aimed to test whether AAV-EV formulations can support efficient site-specific integration while maintaining embryo viability, a critical requirement for generating transgenic models. Fourteen HDR constructs targeting diverse *loci* in mouse (10 constructs) and rats (4 constructs) were packaged as either single-stranded or self-complementary AAV genomes and administered in combination with Cas9/gRNA ribonucleoprotein (RNP) electroporation. The following section details the integration outcomes, dose-dependent effects on viability, and comparative performance of ultracentrifugation-based and ultracentrifugation-free AAV-EV preparations.

All tested HDR constructs were successfully integrated into the target *loci*. Animals were classified as transgene-positive when insertions were confirmed at both 5′ and 3′ ends. All thirteen strains exhibited site-specific integration. Nine ssAAV and three scAAV UC-produced vectors were used to generate twelve independent transgenic lines. Viral titers for zygote treatment were optimized to balance transgenesis efficiency and embryo viability. Three doses were tested: 2.5 x 10^10^ GC (high), 5 x 10^9^ GC (moderate), and 2.5 x 10^9^ GC (low) per medium drop. High titers frequently reduced embryo survival, occasionally resulting in complete lethality, indicating dose-dependent toxicity. Low-titer treatments demonstrated the highest viability rate (number of born animals to transferred embryos, Table 2) - 14.3 %. The moderate dose treatment provided the most consistent outcomes, in terms of transgene-positive animals out of total born animals with an average of 25.9 % (Table 2). In addition, AAV-EV demonstrated an efficient integration in the rat genome, resulting in production of SD-Umod^em1(DelY180-R188)Ccpcz^ SD-Muc1^em1(MUC1fs)Ccpcz^ and *SD-*_Muc1em1(MUC1wt)Ccpcz_ *_lines_*.

**Table 1:**
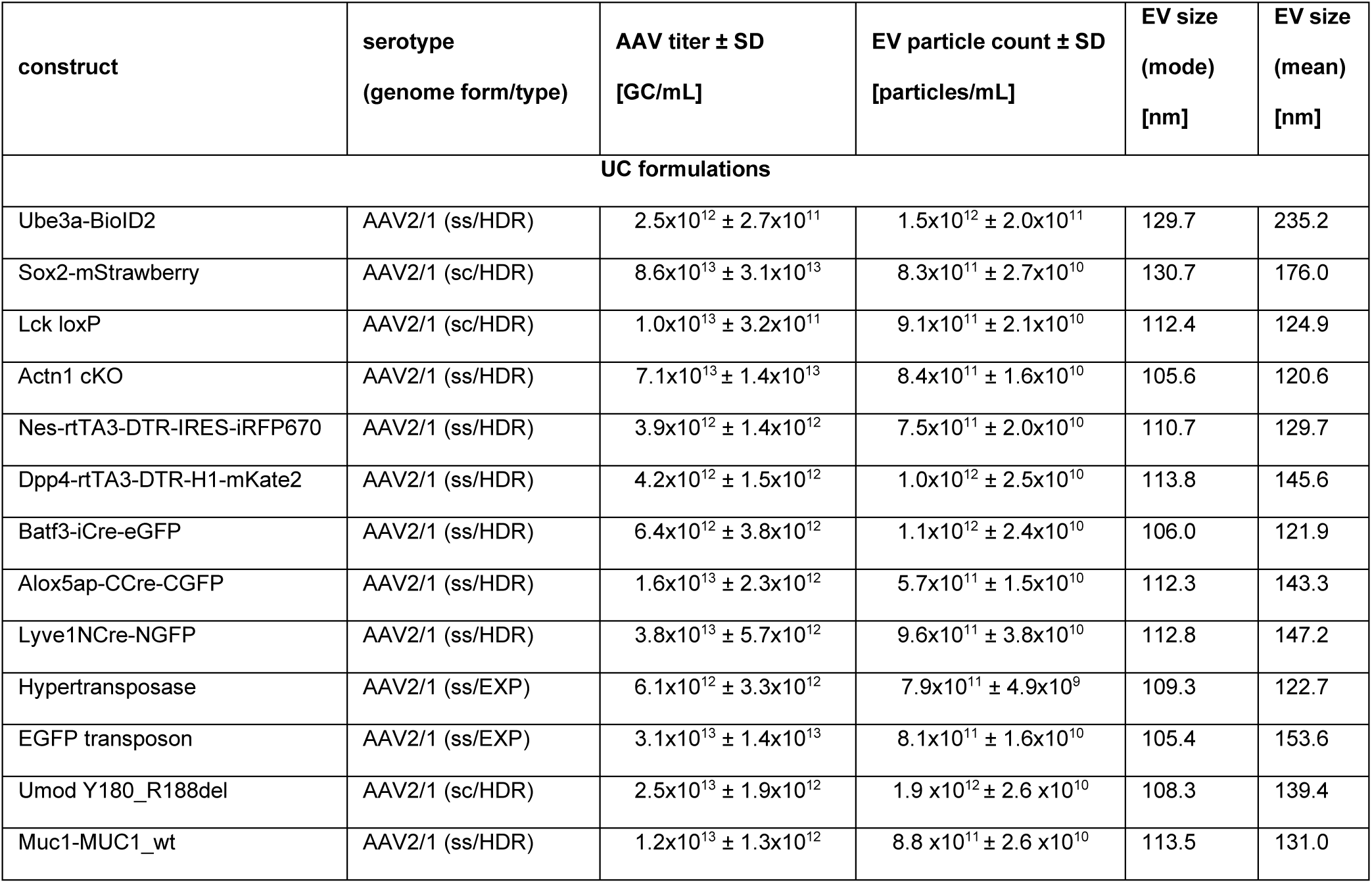
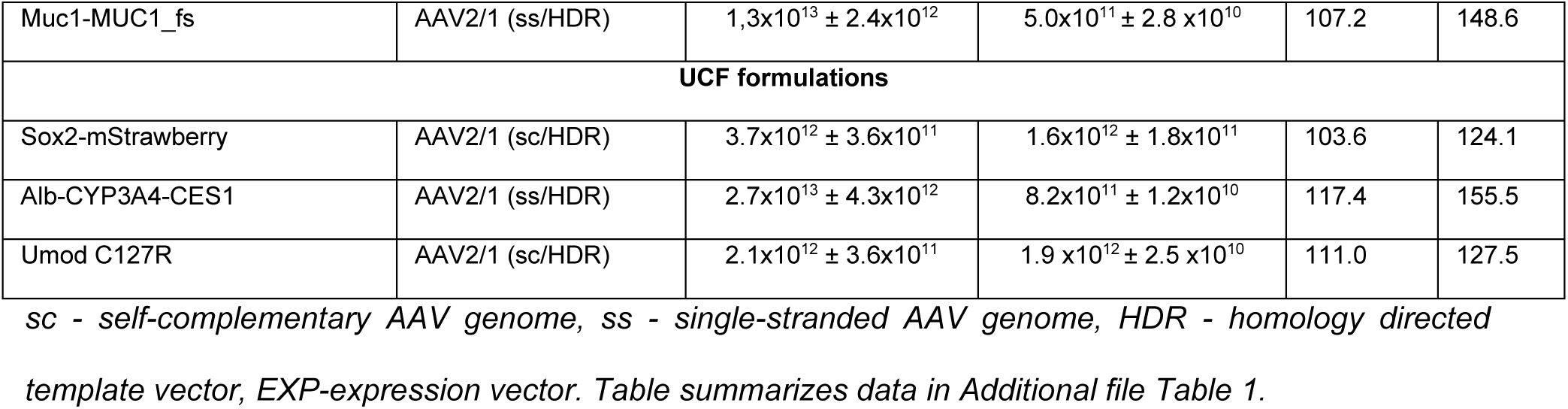
Quantitative characterization of individual AAV-EV formulations obtained by pooling collection 1 (C1) and collection 2 (C2)

**Table 2:** Efficiency of targeted insertion using AAV-EV vectors in mouse.

| UC formulation treatment |  |  |  |  |  |  |  |  |  |  |  |  |  |  |  |
| --- | --- | --- | --- | --- | --- | --- | --- | --- | --- | --- | --- | --- | --- | --- | --- |
| Transgenic construct delivered by AAV-EV | No. of transferred embryos |  |  | No. of born animals |  |  | No. of positive animals |  |  | Targeting efficiency |  |  | Viability |  |  |
| Viral titer per µl (high/medium/ low) | high | med | low | high | med | low | high | med | low | high | med | low | high | med | low |
| Ube3a-BioID2 | 182 | 197 | 172 | 0 | 19 | 16 | 0 | 8 | 6 | 0,00 | 42,10 | 37,50 | 0,00 | 9,60 | 9,30 |
| Sox2-mStrawberry | 64 | 64 | 75 | 3 | 0 | 13 | 1 | 0 | 6 | 33,30 | 0,00 | 46,20 | 4,70 | 0,00 | 17,30 |
| Lck cKO (loxP) | 142 | 36 | 164 | 8 | 0 | 32 | 1 | 0 | 6 | 12,50 | 0,00 | 18,80 | 5,60 | 0,00 | 19,50 |
| Actn1 cKO | 127 | 101 | 115 | 5 | 3 | 4 | 3 | 2 | 2 | 60,00 | 66,70 | 50,00 | 3,90 | 2,90 | 3,80 |
| Nes-rtTA3-DTR-IRES-iRFP670-Far5 | 57 | 78 | 86 | 5 | 6 | 7 | 0 | 2 | 1 | 0,00 | 33,30 | 14,30 | 8,80 | 7,70 | 8,10 |
| Dpp4-rtTA3-DTR-H1-mKate2 | 60 | 68 | 85 | 0 | 4 | 9 | 0 | 2 | 0 | 0,00 | 50,00 | 0,00 | 6,70 | 5,90 | 14,10 |
| Batf3-iCre-eGFP | 54 | 58 | 54 | 3 | 0 | 9 | 1 | 0 | 1 | 33,30 | 0,00 | 11,10 | 5,60 | 0,00 | 18,50 |
| Alox5ap-CCre-CGFP | 38 | 34 | 28 | 9 | 7 | 5 | 1 | 2 | 1 | 11,10 | 28,60 | 20,00 | 23,70 | 20,60 | 17,90 |
| Lyve1-NCre-NGFP | 16 | 58 | 39 | 4 | 16 | 8 | 2 | 2 | 1 | 50,00 | 12,50 | 12,50 | 25,00 | 27,60 | 15,70 |
|  |  |  |  |  |  |  | average (%) |  |  | 22,3 | 25,9 | 23,2 | 9,3 | 8,3 | 14,3 |
| UCF formulation treatment |  |  |  |  |  |  |  |  |  |  |  |  |  |  |  |
| Sox2-mStrawberry | 35 | 54 | 75 | 4 | 12 | 15 | 0 | 2 | 2 | 0 | 16,6 | 13,3 | 11,4 | 22,2 | 20 |
| Alb-CYP3A4-CES1 | 32 | 34 | 35 | 1 | 5 | 2 | 0 | 1 | 0 | 0 | 20 | 0 | 3,1 | 14,7 | 5,8 |

One of the examples is the Sox2-mStrawberry HDR template, which targets *Sox2* locus (Fig. 5A). Correct insertion of the template enables the tracking of *Sox2* expression in blastocyst, specifically in inner cell mass (Fig. 5B). AAV-EV demonstrates successful delivery and insertion of the P2A-mStrawberry cassette into the *Sox2* locus in blastocyst (Fig. 5B and C), and generation of viable C57BL/6NCrl-Sox2^em1(P2A-mStrawberry)Ccpcz^ transgenic mouse line (Fig. 5C).

**Figure 5:**
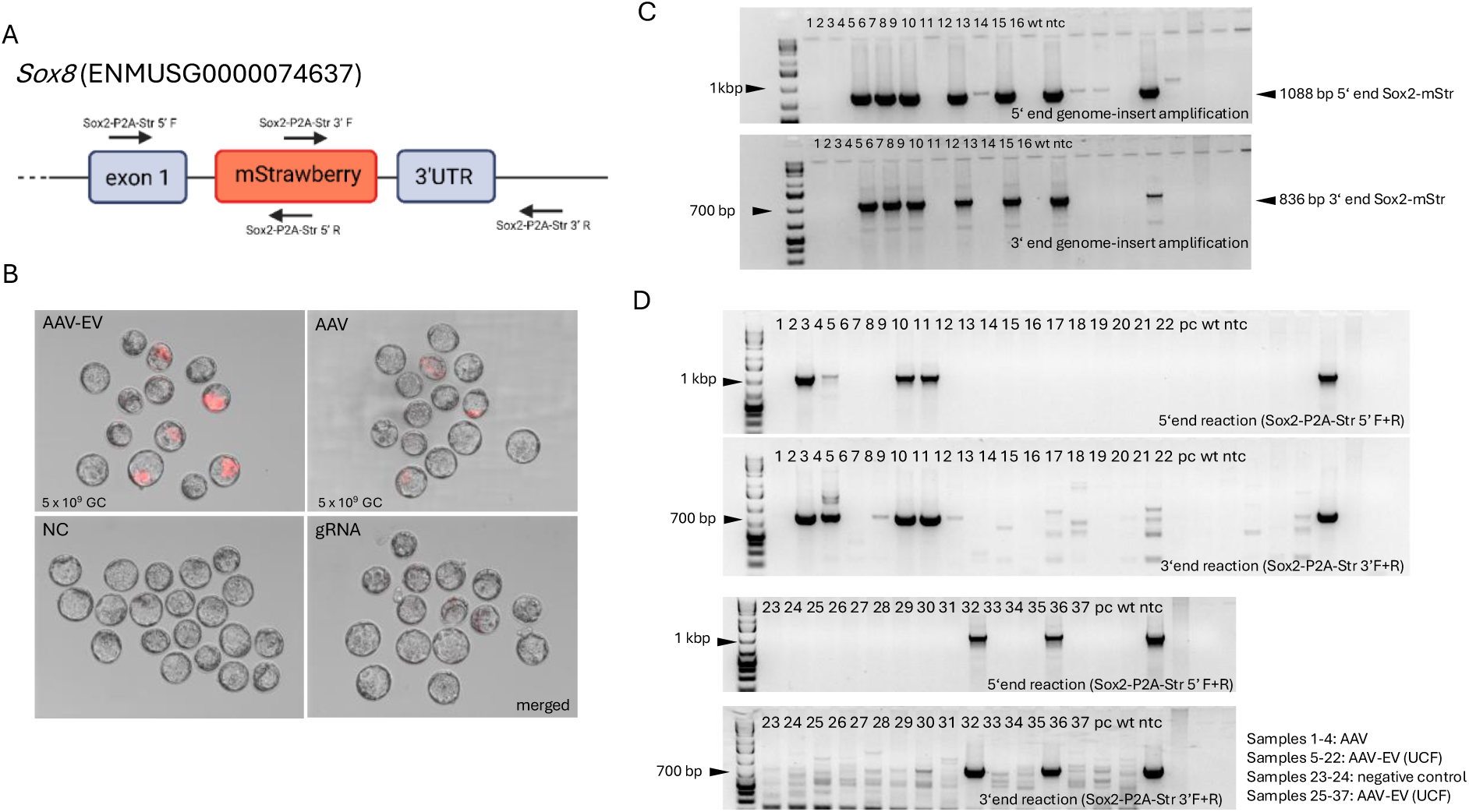
Gene targeting *ex vivo* using AAV-EV vectors. (A) Schematic description of *Sox2* locus and fusion of mStrawberry fluorescent protein via P2A peptide. (B) Blastocyst imaging after AAV-EV and AAV vectors treatment to deliver the Sox2-mStrawberry transgene into the mouse genome using 5 x 10^9^ GC dose with subsequent RNP electroporation. gRNA refers to samples electroporated only with Cas9/gRNA RNP targeting 3’ end of *Sox2* gene. NC-negative control. (C) Agarose gel image: Specific amplification of the genomic-transgene junction at both the 5’ and 3’ ends of the transgene, confirming site-specific insertion. (D) Detection of genome-transgene junction of Sox2-mStrawberry cassette inserted in *Sox2* locus, mice 1-4 were generated by treatment with AAV-Sox2-mStr and 5-22 mice were generated using AAV-EV (UCF) formulation.

UCF-derived AAV-EV formulations were also evaluated for HDR template delivery into zygotes, successfully generating C57BL/6NCrl-Sox2^em1(P2A-mStrawberry)Ccpcz^ (Fig. 5D, Additional file 8 figure J), C57BL/6NCrl-Alb^em6(Cyp3a4,P2A,Ces1)Ccpcz^ mouse transgenic lines (Additional file 8 figure K), and SD-Umod^em1(C127R)Ccpcz^ (Additional file 8 figure L) rat mutant line. These results demonstrated that UCF isolation enables production of a functional AAV-EV vector suitable for *ex vivo* treatment. Tables 2 and 3 summarize viral dose, number of transferred embryos, born and transgene-positive animals, illustrating the relationship between dose, targeting efficiency, and viability. Additional genotyping and allele distribution data are provided in Additional file 8 figure.

**Table 3:** Efficiency of targeted insertion using AAV-EV vectors in rat.

| UC formulation treatment |  |  |  |  |  |  |  |  |  |  |
| --- | --- | --- | --- | --- | --- | --- | --- | --- | --- | --- |
| Transgenic construct delivered by AAV-EV | No. of transferred embryos |  | No. of born animals |  | No. of positive animals |  | Targeting efficiency |  | Viability |  |
| Viral titer per μl (high/medium/low) | med | low | med | low | med | low | med | low | med | low |
| Umod Y180_R188del | 36 | 36 | 15 | 2 | 3 | 0 | 20 | 0 | 41.6 | 0 |
| Muc1-MUC1_wt | - | 75 | - | 17 | - | 2 | - | 11.7 | - | 22.6 |
| Muc1-MUC1_fs | 52 | 111 | 6 | 14 | 0 | 1 | 0 | 7.1 | 11.5 | 12.6 |
| UCF formulation treatment |  |  |  |  |  |  |  |  |  |  |
| Umod C127R | 39 | 17 | 0 | 6 | 0 | 1 | 0 | 16.6 | 0 | 35.3 |

AAV-EV vectors proved highly effective for HDR-mediated genome editing in mouse and rat zygotes, enabling precise site-specific integration across multiple *loci* while maintaining embryo viability. Dose-optimization was critical, as high titers increased toxicity, whereas moderate doses provided the best balance between efficiency and survival. Both UC- and UCF-derived AAV-EV formulations successfully generated transgenic lines, confirming the versatility of this approach. These results highlighted AAV-EV as a promising platform for producing genetically engineered models.

## 4. Discussion

This study establishes an EV-based co-isolation strategy for adeno-associated virus (AAV) vectors that simplifies production while preserving functional performance in highly sensitive systems such as rodent embryos. By combining continuous medium collection with PEG-mediated precipitation and EV-assisted pelleting, we generate composite AAV-EV formulations comprising free AAV, free extracellular vesicles (EVs), and EV-associated AAV particles. Importantly, these preparations bypass density-gradient ultracentrifugation while maintaining delivery efficiency and significantly improving embryo tolerance compared with gradient-purified AAV.

Purification of recombinant AAV is a critical determinant of vector performance, toxicity, and scalability [25]. Discontinuous iodixanol gradients, while widely used, are labor-intensive, technically demanding, and associated with particle aggregation and titer loss, particularly during density-medium removal [8, 9]. AAV-EV approaches build on prior observations that AAV capsids can associate with microvesicles and exosomes during production, generating vectors that display altered uptake, immune shielding, and improved tolerability [14, 16, 19, 20]. Despite these advantages, adoption of EV-associated AAV has been limited by low effective yields, dependence on ultracentrifugation, and scarce validation in sensitive *in vivo* or *ex vivo* contexts. Our work addresses these limitations by demonstrating a scalable, gradient-free workflow (Fig. 1) that produces functional AAV vectors suitable for genome engineering in mouse and rat embryos.

A central feature of this approach is that AAV-EV represents a composite formulation rather than a purified EV-associated AAV subpopulation. Continuous iodixanol gradient analyses confirmed that only a fraction of total AAV genomes co-fractionate with low-density vesicular compartments (1.04-1.21 g/cm³), while the majority localize to higher-density fractions corresponding to free AAV or dense vesicles (Fig. 3) [28, 29]. Importantly, complete physical separation of EV-associated and free AAV was not required to achieve functional benefit. Both ultracentrifugation-based (UC) and ultracentrifugation-free (UCF) preparations supported efficient delivery of HDR templates and expression constructs (Fig. 4 and 5), indicating that EVs primarily function as a biological matrix that modulates vector presentation rather than as obligate carriers.

The UCF protocol represents a key practical advance. Prolonged centrifugation at moderate speed (21,000 x g) enables recovery of EVs and EV-associated AAVs without high-speed ultracentrifugation or density gradients, consistent with established sedimentation behavior of extracellular vesicles [30]. UCF-derived AAV-EVs exhibited titers, particle size distributions (Tab.1), and functional performance comparable to UC preparations and successfully supported HDR-mediated gene targeting in both mouse and rat embryos (Tab.2). By eliminating the need for ultracentrifuges and density-medium handling, this workflow substantially lowers the technical barrier for laboratories engaged in embryo genome engineering.

Proteomic profiling revealed that viral production alters EV composition, resulting in relative depletion of canonical exosome markers such as TSG101, Syntenin-1, ALIX, and CD9, together with enrichment of viral and packaging-related proteins within AAV-EV formulations (Fig. 2). These shifts were formulation-dependent and consistent with previously reported virus-driven remodeling of EV cargo [31]. Despite detection of helper-related proteins at the proteomic level, no Rep or adenoviral DNA-binding protein (DBP) DNA or RNA was detected in treated embryos, arguing against vector replication or persistence *in vivo* under the conditions tested (Additional file 7).

Functional evaluation using the PiggyBac transposon system demonstrated a major advantage of AAV-EV formulations: improved embryo viability at matched and elevated doses without loss of delivery efficiency (Fig. 4). While conventional AAV achieved comparable transgene copy numbers, it did so at the cost of reduced survival (Fig. 4), consistent with prior studies showing dose-, payload-, and promoter-dependent AAV toxicity in early embryos [9, 32–35]. In contrast, AAV-EV vectors supported high-dose administration with sustained developmental competence and increased integration capacity. This expanded dosing window is particularly advantageous for applications requiring high levels of genomic integration, such as transposon-based systems [36, 37].

The improved tolerability of AAV-EV formulations likely reflects qualitative differences in vector presentation rather than reduced effective dose. EVs may buffer AAV particles, limiting capsid aggregation and local overexposure, which are recognized contributors to AAV-induced toxicity during purification and concentration [9, 38]. Partial EV association may also diversify uptake pathways and intracellular trafficking, reducing cellular stress in early embryos, as previously suggested for EV-associated AAV vectors in other systems [14, 16, 39]. Together, these properties provide a plausible mechanistic framework for the observed increase in embryo viability without compromising delivery efficiency.

Extending beyond expression systems, we demonstrated efficient HDR-mediated genome editing across fourteen independent *loci* in mouse and rat embryos using AAV-EV vectors (Tab.2 and 3, Fig. S4). Moderate doses consistently provided the optimal balance between targeting efficiency and viability, yielding an average of approximately 26% correctly targeted offspring across constructs, in line with previously reported AAV-based embryo editing strategies [2–4, 32, 34]. Both UC- and UCF-derived AAV-EV formulations supported generation of viable germline-transmitting animal models, underscoring the robustness and versatility of the platform.

Several limitations should be noted. AAV-EV preparations are heterogeneous, and the precise contribution of EV-associated versus free AAV particles to functional outcomes remains unresolved. The mechanistic basis of reduced toxicity is inferred but not directly tested, and broader *in vivo* applications beyond embryos were not explored here. Further refinement of EV-associated subpopulations and mechanistic studies of uptake and trafficking will be important next steps.

In summary, this work establishes EV-based AAV co-isolation as a practical, scalable, and embryo-tolerant alternative to conventional AAV purification. The approach delivers reproducible titers, improves embryo viability at matched or elevated doses, and supports efficient HDR- and transposon-based genome engineering in rodents. By eliminating density-gradient ultracentrifugation and enabling ultracentrifugation-free workflows, AAV-EV represents an accessible platform that advances the toolkit for rodent genome engineering while aligning with 3R principles.

## 6. Author Contributions

P.N. conceived the experiments, cloning work conducted by P.N. and A.U., production of AAVs conducted by P.N. and M.B., animal work planned and performed by P.N., T.N. and M.V. Characterization of produced formulations was performed by P.N., M.B., T.N., J.Z, J.N. and M.V. The results analysis by P.N., M.B. T.N. and J.Z. Writing manuscript, review, editing; P.N., T.N, M.B, J.Z, V.D., M.V, M.Y., R.S. Funding acquisition, supervision, administration; M.Y, N.A. and R.S. Funding and conceptualization of rodent transgenic and disease models were contributed by P.N., M.S., J.B., P.V., M.Z., S.K., Z.K-S., D.F., O.B., V.N., and O.S. All authors reviewed the manuscript.

## Acknowledgments

This work was supported by the EDUFI Fellowship granted by the Finnish National Agency for Education (EDUFI). We thank the staff of Transgenic and Archiving Module, Czech Centre for Phenogenomics for the assistance during the experiments. Graphical abstracts and diagrams were created with BioRender.com. LC-MS analyses were performed in Laboratory of Mass Spectrometry at Biocev research center, Faculty of Science, Charles University. The authors acknowledge Imaging Methods Core Facility at BIOCEV, supported by the MEYS CR (LM2023050 Czech-BioImaging) and their support with cryo-EM data acquisition & assistance. We thank Xuyu Zhou for generously sharing the Split-Cre constructs based on the SpyTag/SpyCatcher system with J. Balounova. Graphical schemes in this publications were generated with Biorender.

## 8. Availability of data and materials

The datasets supporting the conclusions of this article are included within the article and its additional files.

Additional file 1: figure (.pdf) - Images of AAV-EV and AAV-EV-CD9mE (taggedCD9withmEmerald) pellets from individual collections (C1 and C2) and pooled (P) collections after centrifugation at 100,000×g for 90min.

Additional file 2: table (.xls) - Comparative mass spectrometry analysis dataset (data depicted in plots of Figure 2)

Additional file 3: figure (.pdf) - CryoEM structural analysis of AAV-EV and AAV formulations

Additional file 4: table (.xls) - Absolute quantification of genome copies in individual fractions from a continuous iodixanol gradient (UC_UCF_AAV_GC_quant) and corresponding iodixanol density measurements (data depicted in Figure 3).

Additional file 5: table (.xls) – qPCR copy number analysis of E15.5 embryos and placentas within treatment groups relative to the Rosa26-VFRL-EGFP (MuX) reporter line (EGFP heterozygous control, single-copy reference), data depicted in Figure 4.

Additional file 6: table (.xls.) – Summary of statistical analyses for PiggyBac vector treatments (data depicted in Figure 4)

Additional file 7: tables and figure (.doc) – Normalized expression and copy number analysis of EGFP, Rep, and DBP, with EGFP signal detection in embryos.

Additional file 8: figures and tables (.pdf) Genotyping of Transgenic Rodent Lines by Agarose Gel Electrophoresis and primer sequence (Results Summarized in Tables 2 and 3)

Additional file 9: tables (.xls) and reports (.pdf) - Absolute genome copy quantification in AAV-EV and nanoparticle analysis (particle count, size, quality, and size distribution)

Additional file 10: table (.pptx) – complementary file with Table 2

## Reference

1. Naso MF, Tomkowicz B, Perry WL, Strohl WR. Adeno-Associated Virus (AAV) as a Vector for Gene Therapy. Biodrugs. 2017;31:317. 10.1007/S40259-017-0234-5.

2. Chen S, Sun S, Moonen D, Lee C, Lee AY-F, Schaffer D V., et al. CRISPR-READI: Efficient Generation of Knockin Mice by CRISPR RNP Electroporation and AAV Donor Infection. Cell Rep. 2019;27:3780–3789.e4. 10.1016/j.celrep.2019.05.103.

3. N M, E M, H S, M K, A O, F S, et al. Intra-embryo Gene Cassette Knockin by CRISPR/Cas9-Mediated Genome Editing with Adeno-Associated Viral Vector. iScience. 2018;9:286–97. 10.1016/J.ISCI.2018.10.030.

4. Davis DJ, McNew JF, Maresca-Fichter H, Chen K, Telugu BP, Bryda EC. Efficient DNA knock-in using AAV-mediated delivery with 2-cell embryo CRISPR-Cas9 electroporation. Front Genome Ed. 2023;5:1256451. 10.3389/FGEED.2023.1256451/BIBTEX.

5. Crosson SM, Dib P, Smith JK, Zolotukhin S. Helper-free Production of Laboratory Grade AAV and Purification by Iodixanol Density Gradient Centrifugation. Mol Ther Methods Clin Dev. 2018;10:1–7. 10.1016/J.OMTM.2018.05.001.

6. Florea M, Nicolaou F, Pacouret S, Zinn EM, Sanmiguel J, Andres-Mateos E, et al. High-efficiency purification of divergent AAV serotypes using AAVX affinity chromatography. Mol Ther Methods Clin Dev. 2023;28:146. 10.1016/J.OMTM.2022.12.009.

7. Khanal O, Kumar V, Jin M. Adeno-associated viral capsid stability on anion exchange chromatography column and its impact on empty and full capsid separation. Mol Ther Methods Clin Dev. 2023;31. 10.1016/j.omtm.2023.101112.

8. Zolotukhin S, Byrne BJ, Mason E, Zolotukhin I, Potter M, Chesnut K, et al. Recombinant adeno-associated virus purification using novel methods improves infectious titer and yield. Gene Ther. 1999;6:973–85. 10.1038/SJ.GT.3300938.

9. Wright JF, Le T, Prado J, Bahr-Davidson J, Smith PH, Zhen Z, et al. Identification of factors that contribute to recombinant AAV2 particle aggregation and methods to prevent its occurrence during vector purification and formulation. Mol Ther. 2005;12:171–8. 10.1016/J.YMTHE.2005.02.021.

10. Kalra H, Drummen GPC, Mathivanan S. Focus on Extracellular Vesicles: Introducing the Next Small Big Thing. International Journal of Molecular Sciences 2016, Vol 17, Page 170. 2016;17:170. 10.3390/IJMS17020170.

11. Raposo G, Stoorvogel W. Extracellular vesicles: exosomes, microvesicles, and friends. J Cell Biol. 2013;200:373–83. 10.1083/JCB.201211138.

12. Abels ER, Breakefield XO. Introduction to Extracellular Vesicles: Biogenesis, RNA Cargo Selection, Content, Release, and Uptake. 10.1007/s10571-016-0366-z.

13. Maguire CA, Balaj L, Sivaraman S, Crommentuijn MHW, Ericsson M, Mincheva-Nilsson L, et al. Microvesicle-associated AAV vector as a novel gene delivery system. Molecular Therapy. 2012;20:960–71. 10.1038/mt.2011.303.

14. Deshetty UM, Sil S, Buch S. Gene therapy for the heart: encapsulated viruses to the rescue. Extracell Vesicles Circ Nucleic Acids 2024;5:114–8. 2024;5:114-8. 10.20517/EVCNA.2023.70.

15. György B, Maguire CA. Extracellular vesicles: nature’s nanoparticles for improving gene transfer with adeno-associated virus vectors. Wiley Interdiscip Rev Nanomed Nanobiotechnol. 2018;10:e1488. 10.1002/WNAN.1488.

16. Wang BZ, Luo LJ, Vunjak-Novakovic G. RNA and Protein Delivery by Cell-Secreted and Bioengineered Extracellular Vesicles. Adv Healthc Mater. 2022;11:e2101557. 10.1002/ADHM.202101557.

17. Wang Y, Fu Q, Park SY, Lee YS, Park SY, Lee DY, et al. Decoding cellular mechanism of recombinant adeno-associated virus (rAAV) and engineering host-cell factories toward intensified viral vector manufacturing. Biotechnol Adv. 2024;71:108322. 10.1016/J.BIOTECHADV.2024.108322.

18. György B, Fitzpatrick Z, Crommentuijn MHW, Mu D, Maguire CA. Naturally enveloped AAV vectors for shielding neutralizing antibodies and robust gene delivery invivo. Biomaterials. 2014;35:7598–609. 10.1016/j.biomaterials.2014.05.032.

19. Schiller LT, Lemus-Diaz N, Ferreira RR, Böker KO, Gruber J. Enhanced Production of Exosome-Associated AAV by Overexpression of the Tetraspanin CD9. 2018. 10.1016/j.omtm.2018.03.008.

20. Benskey MJ, Sandoval IM, Manfredsson FP. Continuous collection of adeno-associated virus from producer cell medium significantly increases total viral yield. Hum Gene Ther Methods. 2016;27:32–45. 10.1089/hgtb.2015.117.

21. Rider MA, Hurwitz SN, Meckes DG. ExtraPEG: A polyethylene glycol-based method for enrichment of extracellular vesicles. Sci Rep. 2016;6. 10.1038/srep23978.

22. Wei X, Zhang J, Cui J, Xu W, Zhou X, Ma J. A Split-Cre system designed to detect simultaneous expression of two genes based on SpyTag/SpyCatcher conjugation and Split-GFP dimerization. Journal of Biological Chemistry. 2021;297:101119. 10.1016/j.jbc.2021.101119.

23. Görgens A, Corso G, Hagey DW, Jawad Wiklander R, Gustafsson MO, Felldin U, et al. Identification of storage conditions stabilizing extracellular vesicles preparations. J Extracell Vesicles. 2022;11:e12238. 10.1002/JEV2.12238.

24. Zini J, Saari H, Ciana P, Viitala T, Lõhmus A, Saarinen J, et al. Infrared and Raman spectroscopy for purity assessment of extracellular vesicles. European Journal of Pharmaceutical Sciences. 2022;172:106135. 10.1016/J.EJPS.2022.106135.

25. Aurnhammer C, Haase M, Muether N, Hausl M, Rauschhuber C, Huber I, et al. Universal real-time PCR for the detection and quantification of adeno-associated virus serotype 2-derived inverted terminal repeat sequences. Hum Gene Ther Methods. 2012;23:18–28. 10.1089/hgtb.2011.034.

26. Pfaffl MW. A new mathematical model for relative quantification in real-time RT-PCR. Nucleic Acids Res. 2001;29:e45. 10.1093/NAR/29.9.E45.

27. Onódi Z, Pelyhe C, Nagy CT, Brenner GB, Almási L, Kittel Á, et al. Isolation of high-purity extracellular vesicles by the combination of iodixanol density gradient ultracentrifugation and bind-elute chromatography from blood plasma. Front Physiol. 2018;9 OCT:1479. 10.3389/FPHYS.2018.01479/FULL.

28. Graham J, Ford T, Rickwood D. The Preparation of Subcellular Organelles from Mouse Liver in Self-Generated Gradients of Iodixanol. Anal Biochem. 1994;220:367–73. 10.1006/ABIO.1994.1351.

29. Livshts MA, Khomyakova E, Evtushenko EG, Lazarev VN, Kulemin NA, Semina SE, et al. Isolation of exosomes by differential centrifugation: Theoretical analysis of a commonly used protocol. Scientific Reports 2015 5:1. 2015;5:1-14. 10.1038/srep17319.

30. Li X, La Salvia S, Liang Y, Adamiak M, Kohlbrenner E, Jeong D, et al. Extracellular Vesicle-Encapsulated Adeno-Associated Viruses for Therapeutic Gene Delivery to the Heart. Circulation. 2023;148:405–25. 10.1161/CIRCULATIONAHA.122.063759/SUPPL_FILE/COTR148_05.PDF.

31. Nickl P, Jenickova I, Elias J, Kasparek P, Barinka C, Kopkanova J, et al. Multistep allelic conversion in mouse pre-implantation embryos by AAV vectors. Scientific Reports 2024 14:1. 2024;14:1–13. 10.1038/s41598-024-70853-1.

32. Romeo C, Chen SH, Goulding E, Van Gorder L, Schwartz M, Walker M, et al. AAV Diffuses across Zona Pellucida for Effortless Gene Delivery to Fertilized Eggs. Biochem Biophys Res Commun. 2020;526:85. 10.1016/J.BBRC.2020.03.026.

33. Duddy G, Courtis K, Horwood J, Olsen J, Horsler H, Hodgson T, et al. Donor template delivery by recombinant adeno-associated virus for the production of knock-in mice. BMC Biol. 2024;22:1–12. 10.1186/S12915-024-01834-Z/FIGURES/6.

34. Imre G, Takács B, Czipa E, Drubi AB, Jaksa G, Latinovics D, et al. Prolonged activity of the transposase helper may raise safety concerns during DNA transposon-based gene therapy. Mol Ther Methods Clin Dev. 2023;29:145–59. 10.1016/J.OMTM.2023.03.003/ASSET/7ADAD607-F149-4F1D-9F6E-BB7DA8EE59EE/MAIN.ASSETS/GR3.JPG.

35. Cabanes-Creus M, Liao SHY, Gale Navarro R, Knight M, Nazareth D, Lau NS, et al. Harnessing whole human liver ex situ normothermic perfusion for preclinical AAV vector evaluation. Nature Communications 2024 15:1. 2024;15:1876-. 10.1038/s41467-024-46194-y.

36. Keng CT, Guo K, Liu YC, Shen KY, Lim DS, Lovatt M, et al. Multiplex viral tropism assay in complex cell populations with single-cell resolution. Gene Therapy 2022 29:9. 2022;29:555–65. 10.1038/s41434-022-00360-3.

37. Vyas P, Balakier H, Librach CL. Ultrastructural identification of CD9 positive extracellular vesicles released from human embryos and transported through the zona pellucida. Syst Biol Reprod Med. 2019;65:273–80. 10.1080/19396368.2019.1619858.

38. Hudry E, Martin C, Gandhi S, György B, Scheffer DI, Mu D, et al. Exosome-associated AAV vector as a robust and convenient neuroscience tool. Gene Ther. 2016;23:380–92. 10.1038/gt.2016.11.

