## Additional file 1 for "Adeno-Associated Virus Co-Precipitation with Extracellular Vesicles for Genome Editing in Rodent Embryo"

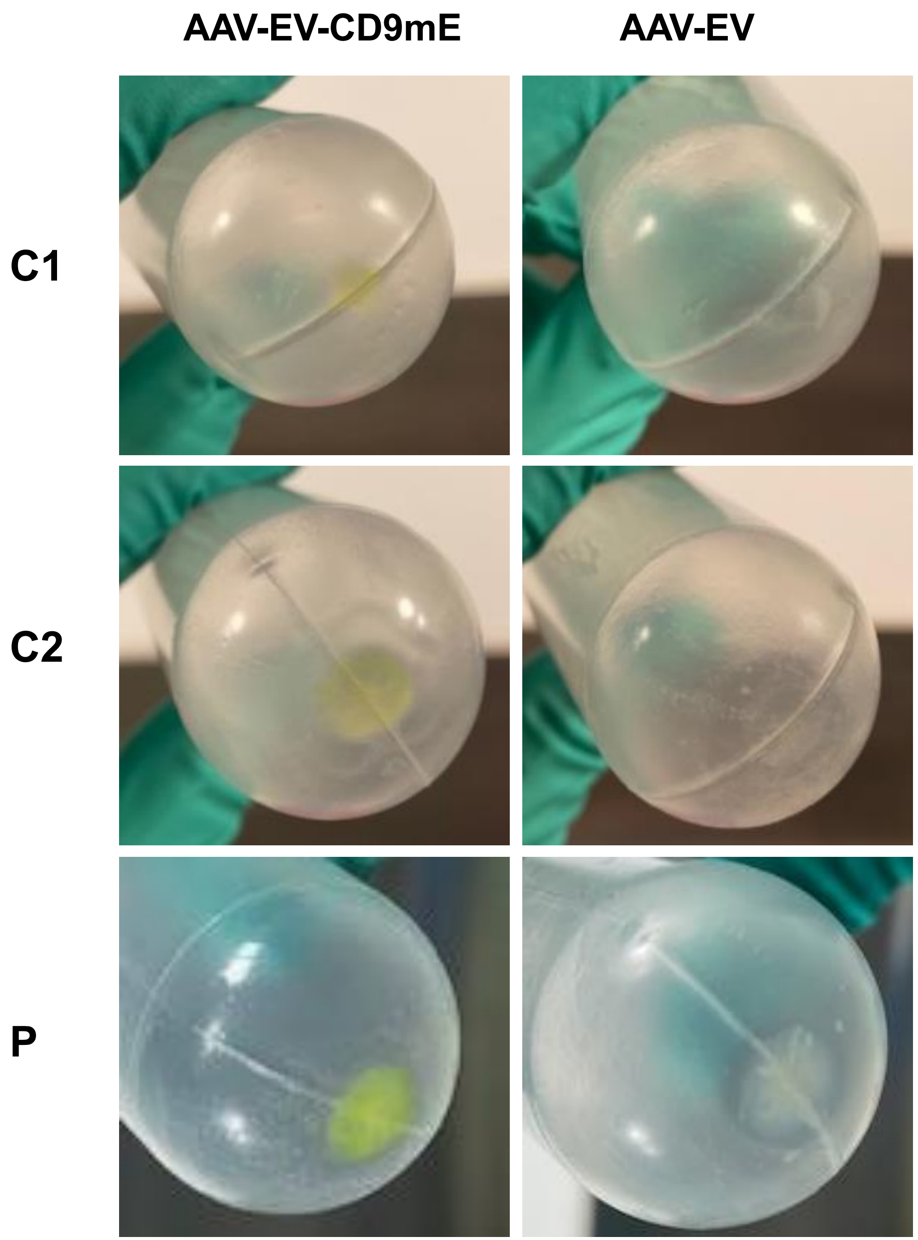


**Additional file 1 figure:** Images of AAV-EV and AAV-EV-CD9mE (tagged CD9 with mEmerald) pellets from individual collections (C1 and C2) and pooled (P) collections after centrifugation at 100,000×g for 90 min.
