## Additional file 3 for "Adeno-Associated Virus Co-Precipitation with Extracellular Vesicles for Genome Editing in Rodent Embryo"

A

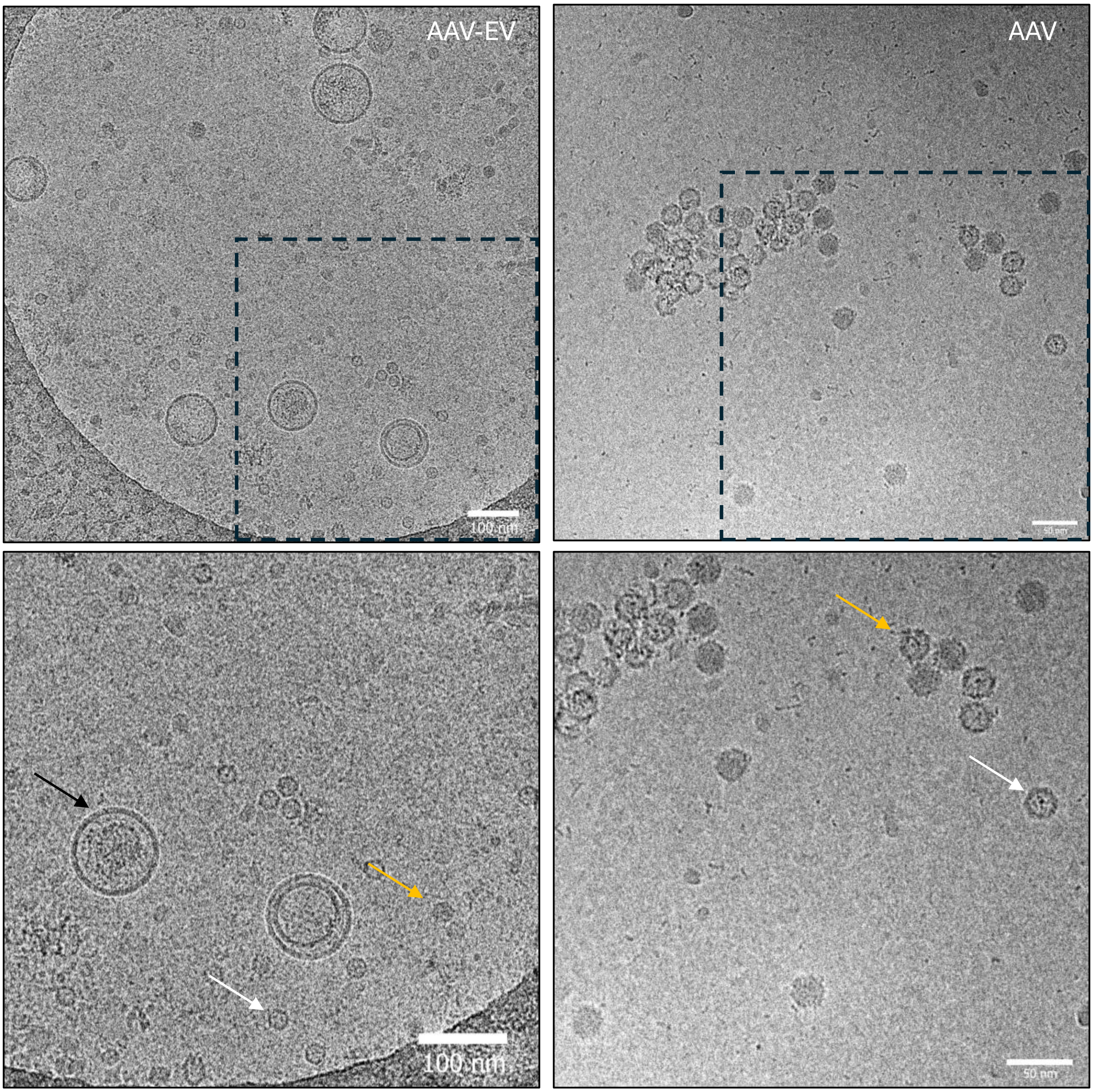

B

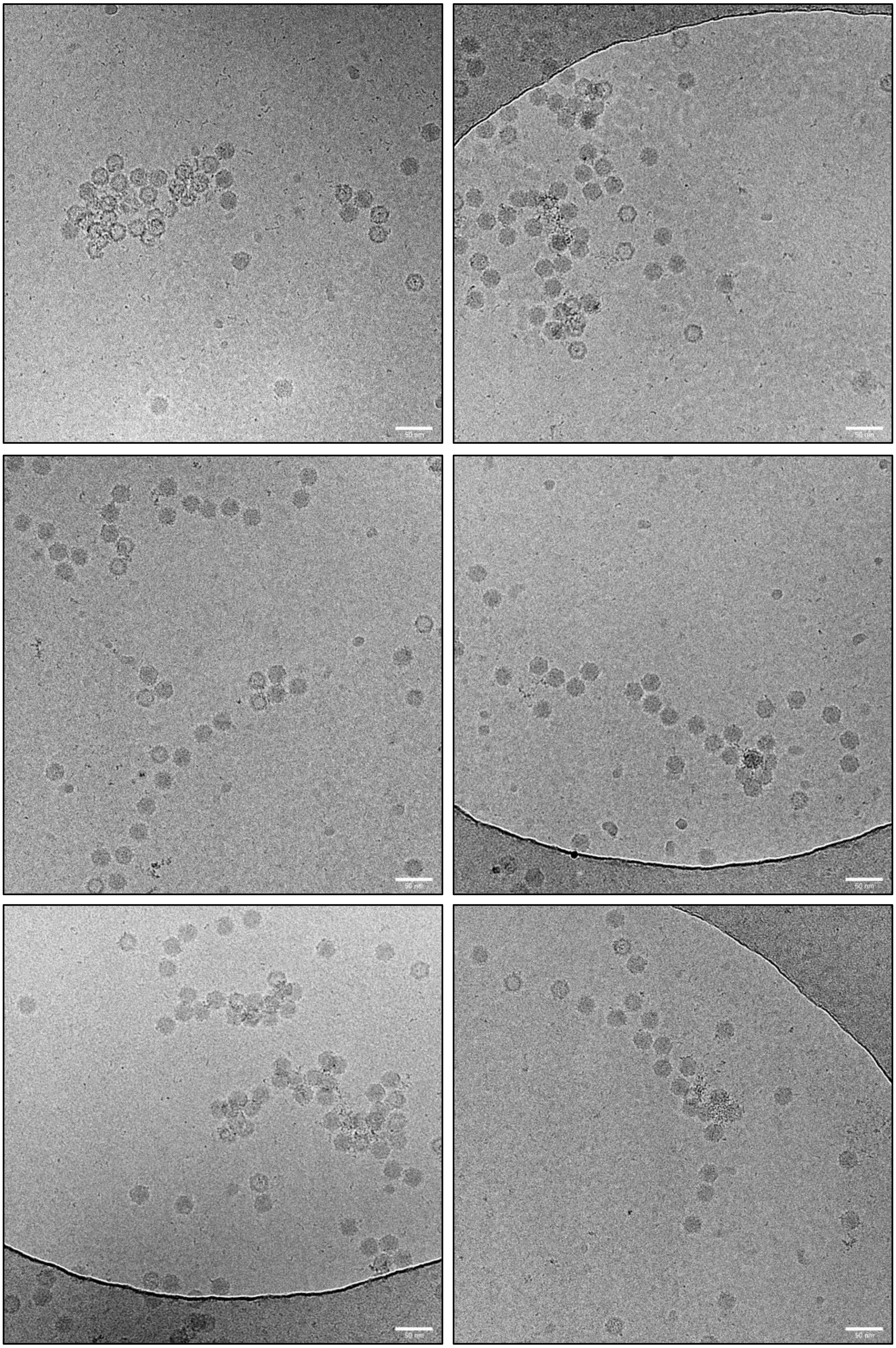

B

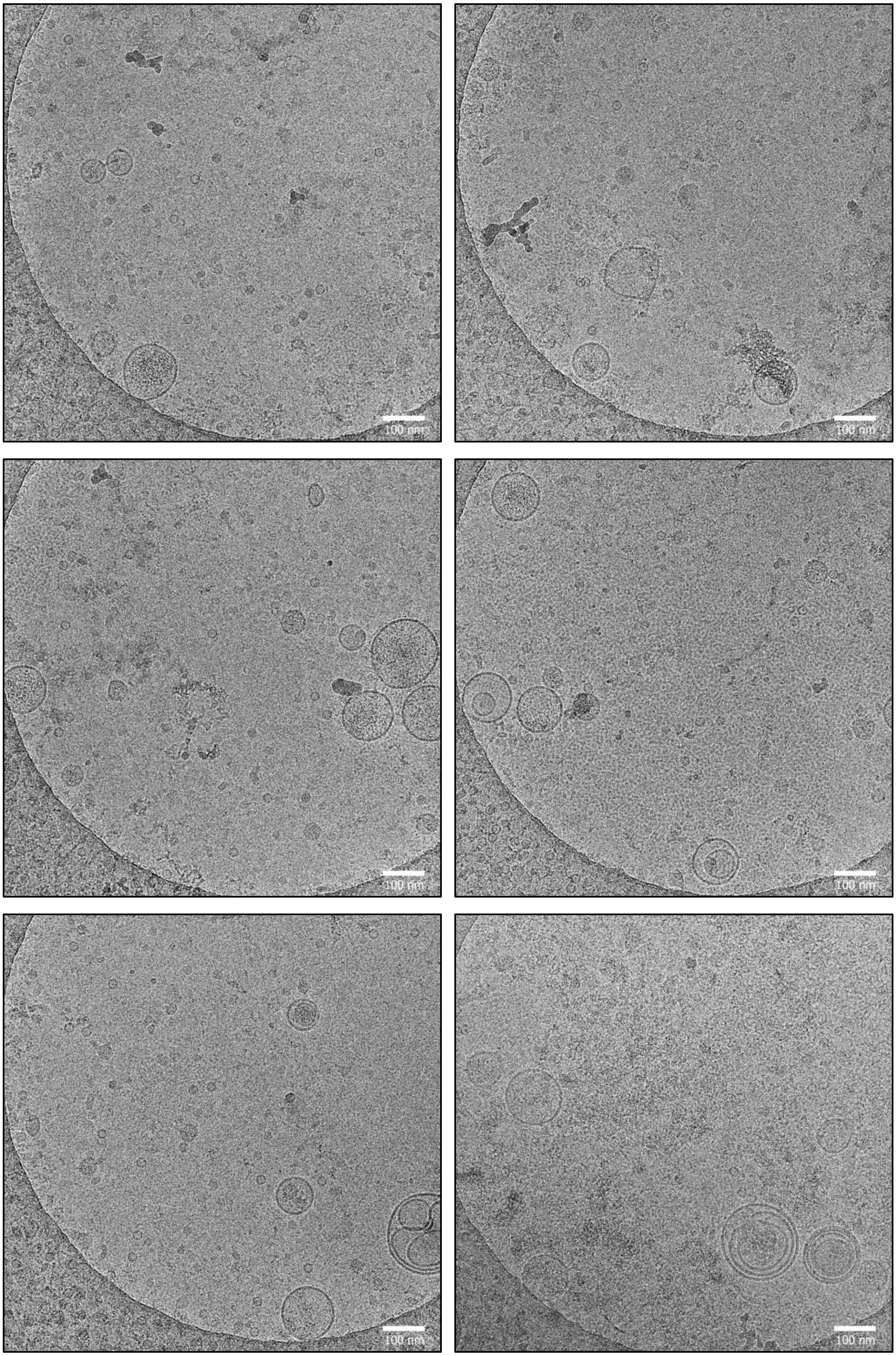


**Additional file 3 figure: CryoEM structural analysis of formulations** (A) AAV-EV and AAVs formulations; empty AAV capsid (white arrow), full AAV capsid (yellow arrow), vesicle (black arrow), (B) Additional images of AAV formulations. (C) Additional images of AAV-EV formulations.
