## Additional file 7 for "Adeno-Associated Virus Co-Precipitation with Extracellular Vesicles for Genome Editing in Rodent Embryo"

|  | **Embryo** | **Average Ct values** | **Ct values SD** | **Average Ct values** | **Ct values SD** | **Average Ct values** | **Average Ct values** | **Average Ct values** | **Average Ct values** |
| --- | --- | --- | --- | --- | --- | --- | --- | --- | --- |
|  | **Detection of:** | **EGFP (DNA)** | **EGFP (DNA)** | **EGFP (RNA)** | **EGFP (RNA)** | **Rep (DNA)** | **Rep (RNA)** | **DBP (DNA)** | **DBP (RNA)** |
|  | positive control | 24,42 | 0,06 | 20,37 | 0,00 | x | x | x | x |
|  | negative control | 36,56 | 0,64 | 31,95 | 0,49 | >36 | >40 | >40 | >40 |
| **AAV** | E336 | 22,59 | 0,24 | 22,58 | 0,26 | >36 | >40 | >40 | >40 |
|  | E337 | 34,38 | 0,24 | 22,96 | 0,49 | >36 | >40 | >40 | >40 |
|  | E339 | 25,64 | 0,15 | 22,99 | 0,44 | >36 | >40 | >40 | >40 |
|  | E340 | 21,65 | 0,22 | 19,58 | 0,24 | >36 | >40 | >40 | >40 |
|  | E341 | 22,92 | 0,08 | 16,13 | 0,32 | >36 | >40 | >40 | >40 |
| **AAV-EV** | E348 | 24,16 | 0,10 | 21,16 | 1,02 | >36 | >40 | >40 | >40 |
|  | E349 | 22,87 | 0,05 | 22,04 | 0,50 | >36 | >40 | >40 | >40 |
|  | E350 | 25,11 | 0,20 | 25,93 | 0,45 | >36 | >40 | >40 | >40 |
|  | E351 | 22,79 | 0,04 | 22,22 | 0,47 | >36 | >40 | >40 | >40 |
|  | E352 | 36,12 | 1,05 | 27,00 | 1,10 | >36 | >40 | >40 | >40 |
|  | E353 | 22,02 | 0,04 | 15,64 | 0,30 | >36 | >40 | >40 | >40 |
|  | E354 | 22,52 | 0,21 | 15,37 | 0,46 | >36 | >40 | >40 | >40 |
|  | E355 | 21,77 | 0,07 | 15,79 | 0,23 | >36 | >40 | >40 | >40 |
|  | E356 | 24,68 | 0,11 | 17,96 | 0,48 | >36 | >40 | >40 | >40 |

Table: Cycle threshold (Ct) values of target genes EGFP, Rep, and DBP analyzed from DNA and cDNA samples.

Table: Standard dilution series used for reference in Rep copy number analysis

| **No. of molecules** | **Ct1** | **Ct1** | **Ct1** | **Mean Ct** |
| --- | --- | --- | --- | --- |
| 10^8 | 7,81 | 7,77 | 7,76 | 7,78 |
| 10^7 | 12,12 | 12,27 | 12,44 | 12,27 |
| 10^6 | 16,24 | 16,07 | 16,06 | 16,12 |
| 10^5 | 20,81 | 20,77 | 20,75 | 20,77 |
| 10^4 | 25,75 | 25,68 | 25,68 | 25,70 |
| 10^3 | 29,33 | 29,26 | 29,23 | 29,27 |

Table: Standard dilution series used for reference in DBP copy number analysis

| **No. of molecules** | **Ct1** | **Ct1** | **Ct1** | **Mean Ct** |
| --- | --- | --- | --- | --- |
| 10^8 | 8,28 | 7,81 | 8,11 | 8,06 |
| 10^7 | 12,49 | 12,59 | 12,23 | 12,43 |
| 10^6 | 16,82 | 16,57 | 16,58 | 16,65 |
| 10^5 | 21,51 | 21,21 | 21,28 | 21,33 |
| 10^4 | 26,62 | 26,02 | 26,2 | 26,28 |
| 10^3 | 29,77 | 29,92 | 29,91 | 29,86 |

|  | Embryo | Average Ct values | Ct values SD | Average Ct values | Ct values SD | Average Ct values | Ct values SD | Average Ct values | Ct values SD | Average Ct values | Ct values SD | Average Ct values | Ct values SD |
| --- | --- | --- | --- | --- | --- | --- | --- | --- | --- | --- | --- | --- | --- |
|  | Detection of: | EGFP/Ubb (DNA) | EGFP/Ubb (DNA) | EGFP/Ubb (RNA) | EGFP/Ubb (RNA) | Rep/Ubb (DNA) | Rep/Ubb (DNA) | Rep/Ubb (RNA) | Rep/Ubb (RNA) | DBP/Ubb (DNA) | DBP/Ubb (DNA) | DBP/Ubb (RNA) | DBP/Ubb (RNA) |
|  | positive control | 20,16 | 0,03 | 15,05 | 0,43 | x | x | x | x | x | x | x | x |
|  | negative control | 20,69 | 0,15 | 16,34 | 0,21 | 21,29 | 0,30 | 16,68 | 0,23 | 21,42 | 0,13 | 16,58 | 0,11 |
| AAV | E336 | 19,88 | 0,07 | 18,14 | 0,15 | 20,27 | 0,39 | 19,83 | 0,17 | 20,14 | 0,08 | 19,86 | 0,07 |
|  | E337 | 20,83 | 0,03 | 13,99 | 0,36 | 21,51 | 0,18 | 15,76 | 0,10 | 21,57 | 0,18 | 15,72 | 0,01 |
|  | E339 | 20,42 | 0,04 | 14,63 | 0,08 | 21,03 | 0,29 | 16,13 | 0,31 | 21,33 | 0,12 | 16,12 | 0,22 |
|  | E340 | 20,09 | 0,12 | 16,12 | 0,17 | 20,32 | 0,21 | 17,74 | 0,11 | 20,52 | 0,07 | 17,69 | 0,30 |
|  | E341 | 20,78 | 0,15 | 14,05 | 0,11 | 21,30 | 0,35 | 15,14 | 0,43 | 21,68 | 0,38 | 15,14 | 0,31 |
| AAV-EV | E348 | 20,50 | 0,05 | 14,69 | 0,07 | 21,29 | 0,40 | 17,14 | 0,43 | 21,28 | 0,52 | 16,88 | 0,28 |
|  | E349 | 20,66 | 0,02 | 13,94 | 0,35 | 21,44 | 0,35 | 15,37 | 0,44 | 21,55 | 0,38 | 15,20 | 0,28 |
|  | E350 | 20,88 | 0,13 | 14,43 | 0,11 | 21,67 | 0,16 | 16,24 | 0,52 | 21,93 | 0,60 | 15,96 | 0,14 |
|  | E351 | 20,59 | 0,12 | 15,51 | 0,12 | 21,06 | 0,40 | 16,89 | 0,16 | 21,34 | 0,29 | 16,56 | 0,44 |
|  | E352 | 20,43 | 0,08 | 14,09 | 0,18 | 20,91 | 0,29 | 15,71 | 0,11 | 21,25 | 0,19 | 15,58 | 0,10 |
|  | E353 | 20,76 | 0,09 | 14,30 | 0,03 | 21,30 | 0,30 | 16,06 | 0,06 | 21,67 | 0,18 | 15,85 | 0,12 |
|  | E354 | 20,48 | 0,13 | 13,69 | 0,02 | 21,01 | 0,22 | 15,69 | 0,10 | 21,09 | 0,19 | 15,56 | 0,04 |
|  | E355 | 20,51 | 0,09 | 14,58 | 0,12 | 21,11 | 0,23 | 16,36 | 0,17 | 21,25 | 0,27 | 16,24 | 0,02 |
|  | E356 | 20,55 | 0,04 | 14,15 | 0,24 | 21,41 | 0,12 | 15,88 | 0,14 | 21,22 | 0,25 | 15,77 | 0,04 |

Table: Cycle threshold (Ct) values of the Ubb reference gene in DNA and cDNA samples for relative quantification analysis.

Table: Comparison of copy numbers in analyzed samples with the Rosa26‑VFRL‑EGFP (MuX) reporter (heterozygous, single‑copy reference)

|  | Embryo | Average Ct values | SD | dCt | SD | Copy number | SD |
| --- | --- | --- | --- | --- | --- | --- | --- |
|  | Detection of: | EGFP/Ubb (DNA) | EGFP/Ubb (DNA) | EGFP/Ubb (DNA) | EGFP/Ubb (DNA) | EGFP/Ubb (DNA) | EGFP/Ubb (DNA) |
|  | positive control | 24.42 | 0.06 | 4.26 | 0.07 | 1.0 | 0.05 |
|  | negative control | 36.56 | 0.64 | 15.87 | 0.66 | 0.0 | 0.00 |
| AAV | E336 | 22.59 | 0.24 | 2.71 | 0.25 | 2.9 | 0.55 |
|  | E337 | 34.38 | 0.24 | 13.55 | 0.24 | 0.0 | 0.00 |
|  | E339 | 25.64 | 0.15 | 5.22 | 0.16 | 0.5 | 0.06 |
|  | E340 | 21.65 | 0.22 | 1.56 | 0.25 | 6.5 | 1.23 |
|  | E341 | 22.92 | 0.08 | 2.14 | 0.17 | 4.3 | 0.54 |
| AAV-EV | E348 | 24.16 | 0.1 | 3.66 | 0.11 | 1.5 | 0.12 |
|  | E349 | 22.87 | 0.05 | 2.21 | 0.05 | 4.1 | 0.16 |
|  | E350 | 25.11 | 0.2 | 4.23 | 0.24 | 1.0 | 0.18 |
|  | E351 | 22.79 | 0.04 | 2.2 | 0.13 | 4.2 | 0.38 |
|  | E352 | 36.12 | 1.05 | 15.69 | 1.05 | 0.0 | 0.00 |
|  | E353 | 22.02 | 0.04 | 1.26 | 0.10 | 8.0 | 0.57 |
|  | E354 | 22.52 | 0.21 | 2.04 | 0.25 | 4.7 | 0.87 |
|  | E355 | 21.77 | 0.07 | 1.26 | 0.11 | 8.0 | 0.66 |
|  | E356 | 24.68 | 0.11 | 4.13 | 0.12 | 1.1 | 0.09 |

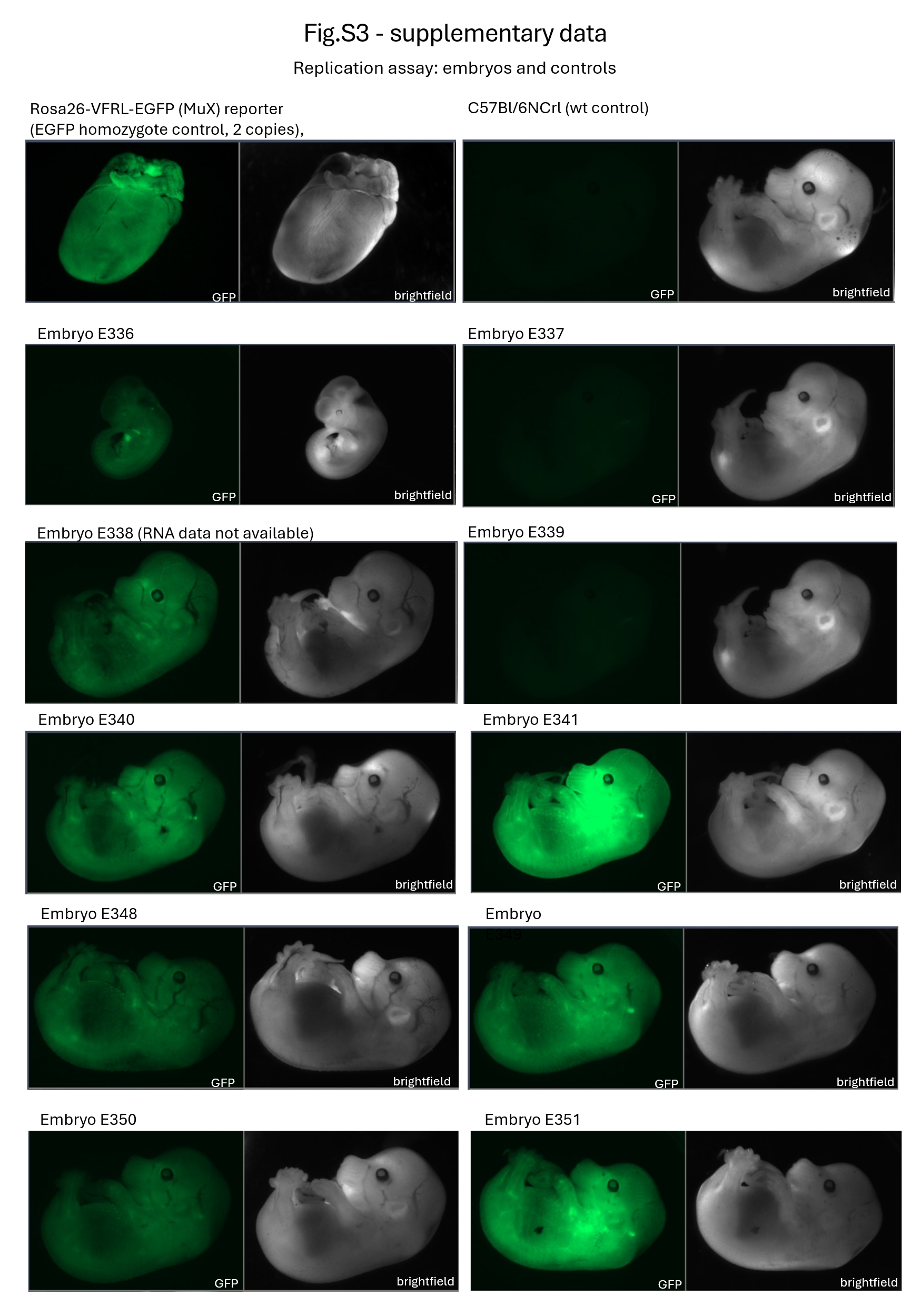

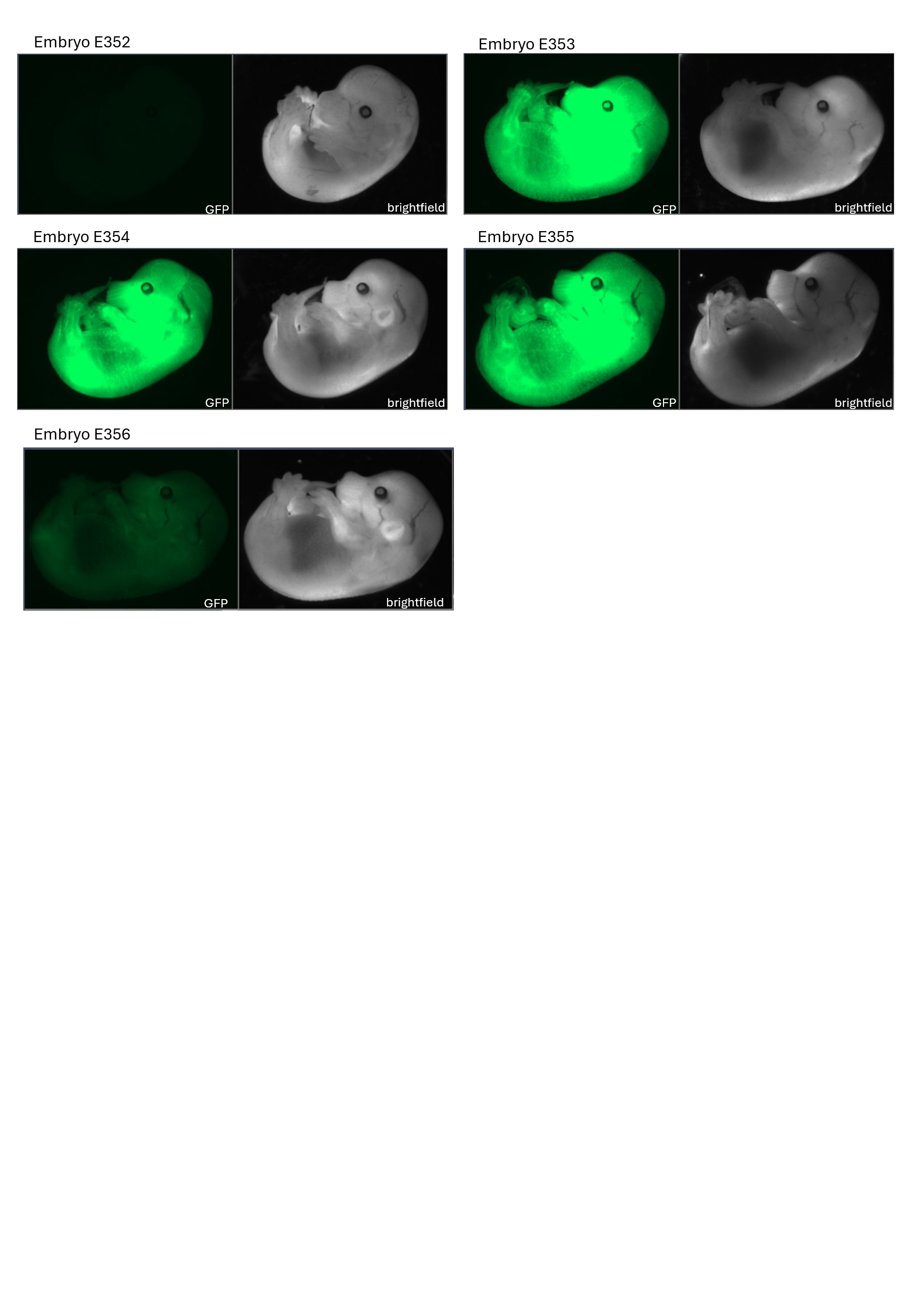

**Additional File 7, Figure: EGFP fluorescence in treated and control E15.5 embryos, assessed for replication potential of AAV‑EV compared with standard AAV vectors.**

Rosa26-VFRL-EGFP (MuX) reporter mouse line was used as homozygote control, 2 EGFP copies per genome.

**Table: Primers used for copy number analysis and replication assay**

| **Genes of interest** | **sequence** | **product** |
| --- | --- | --- |
| Rep_F1 | TCACCAAGCAGGAAGTCAAAG | 144 bp |
| Rep_R1 | CCCGTTTGGGCTCACTTATATC |  |
| Ubb_F | ATGTGAAGGCCAAGATCCAG | 160 bp |
| Ubb_R | TAATAGCCACCCCTCAGACG |  |
| DBP_F1 | CACTGGGTCGTCTTCATTCA | 98 bp |
| DBP_R1 | CGCTACAAATGGTGGGTTTC |  |
| EGFP_F1 | ACGACGGCAACTACAAGACC | 132 bp |
| EGFP_R1 | TGTAGTTGTACTCCAGCTTGTGC |  |
