## Additional file 8 for "Adeno-Associated Virus Co-Precipitation with Extracellular Vesicles for Genome Editing in Rodent Embryo"

#### A) C57BL/6NCrl-Ube3a$^{em2(BioID2)Ccpcz}$

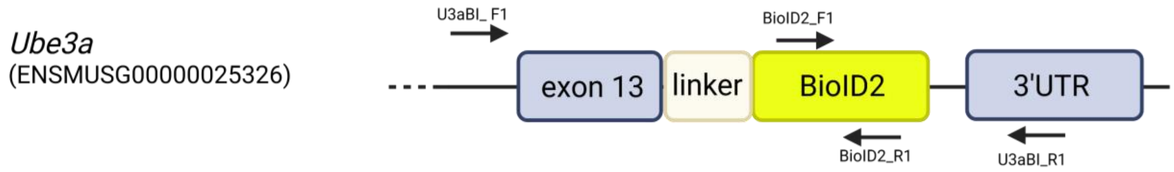


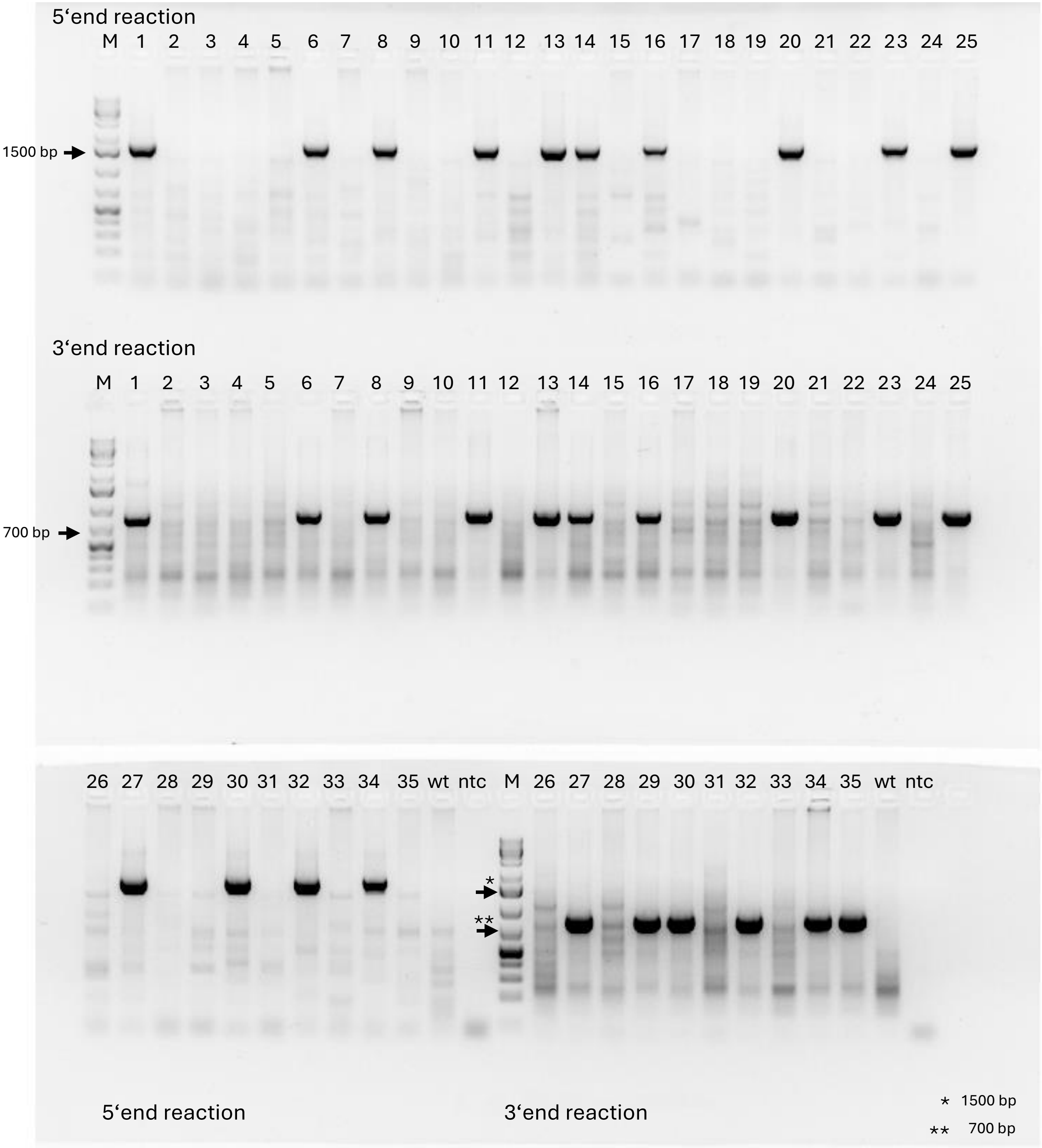


| 5‘reaction | Primers sequence (5‘>3‘) | Product size (bp) |
|---|---|---|
| U3aBI_F1 | CCATATTCCATACACGCAAGCAG | 1672 |
| BioID2_R1 | TCGCCCAGGTACAGCATC | |
| **3‘reaction** | **Primers sequence (5‘>3‘)** | **Product size (bp)** |
| U3aBI_R1 | CCAATGAAGAAGGGAGGCAC | 863 |
| BioID2_F1 | ACCCAGGAGAGACTGAAGG | |

| Low dose |
|---|
| 1* |
| 2 |
| 3 |
| 4 |
| 5 |
| 6* |
| 7 |
| 8* |
| 9 |
| 29 |
| 30* |
| 31 |
| 32* |
| 33 |
| 34* |
| 35 |
| **Medium dose** |
| 10 |
| 11* |
| 12 |
| 13* |
| 14* |
| 15 |
| 16* |
| 17 |
| 18 |
| 19 |
| 20* |
| 21 |
| 22 |
| 23* |
| 24 |
| 25* |
| 26 |
| 27* |
| 28 |

### B) C57BL/6NCrl-Sox2$^{em1(P2A-mStrawberry)Ccpcz}$

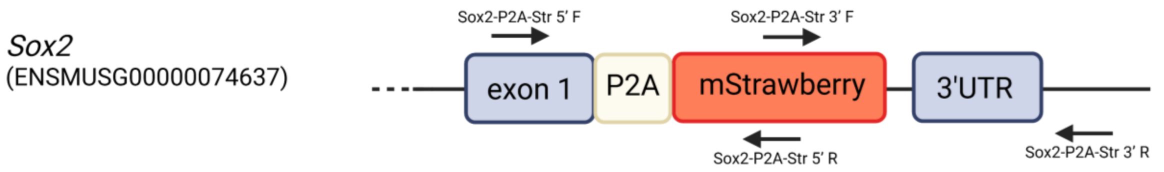


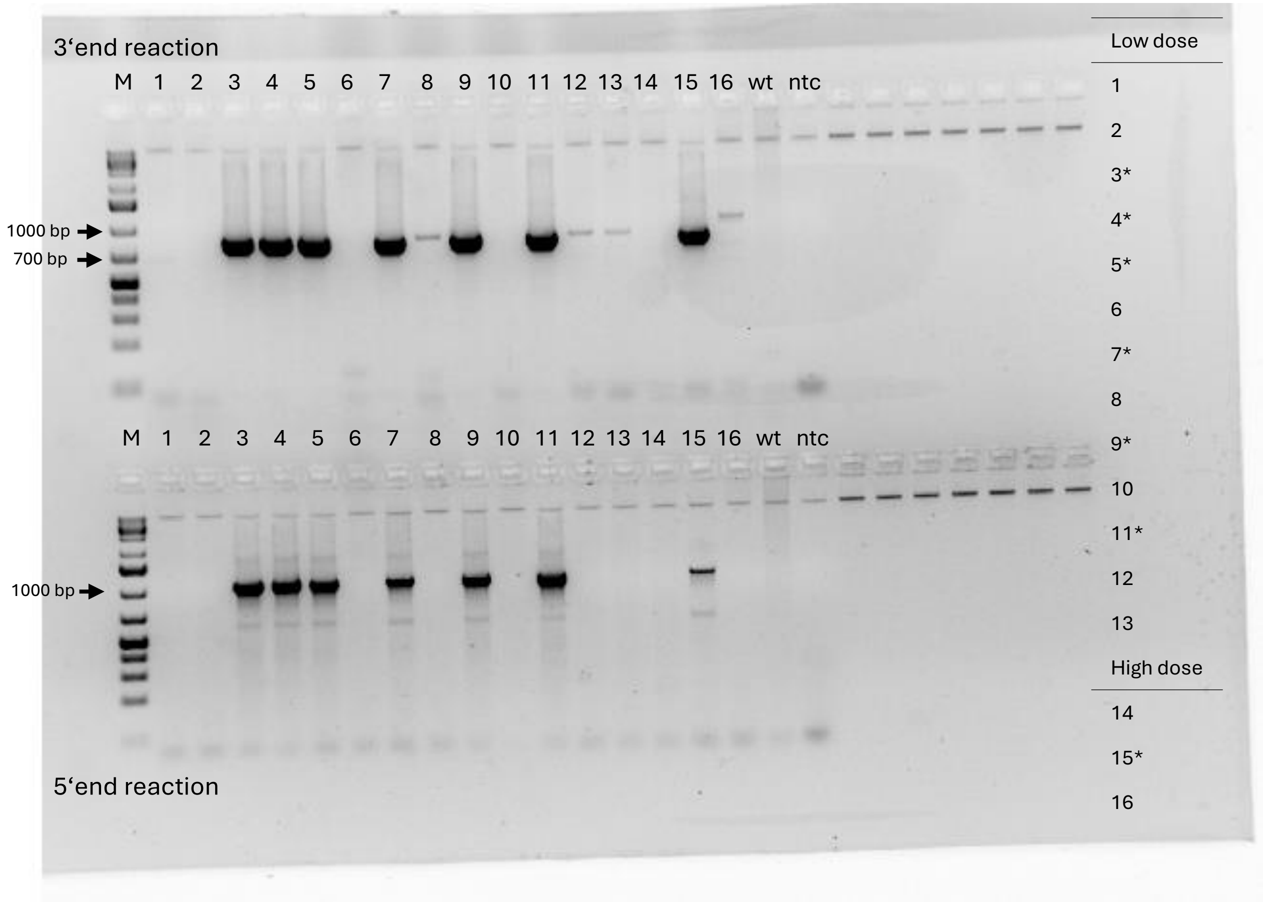


| 3‘reaction | Primers sequence (5‘>3‘) | Product size (bp) |
|---|---|---|
| Sox2-P2A-Str 3' F | GCAGAAGAAGACCATGGGCT | 836 |
| Sox2-P2A-Str 3' R | CCCAGCAAGAACCCTTTCCT | |
| **5‘reaction** | **Primers sequence (5‘>3‘)** | **Product size (bp)** |
| Sox2-P2A-Str 5' F | TAAGTACACGCTTCCCGGAG | 1088 |
| Sox2-P2A-Str 5' R | AGCCCATGGTCTTCTTCTGC | |

#### C) C57BL/6NCrl-Lck$^{em2Ccpcz}$ (loxP insertion)

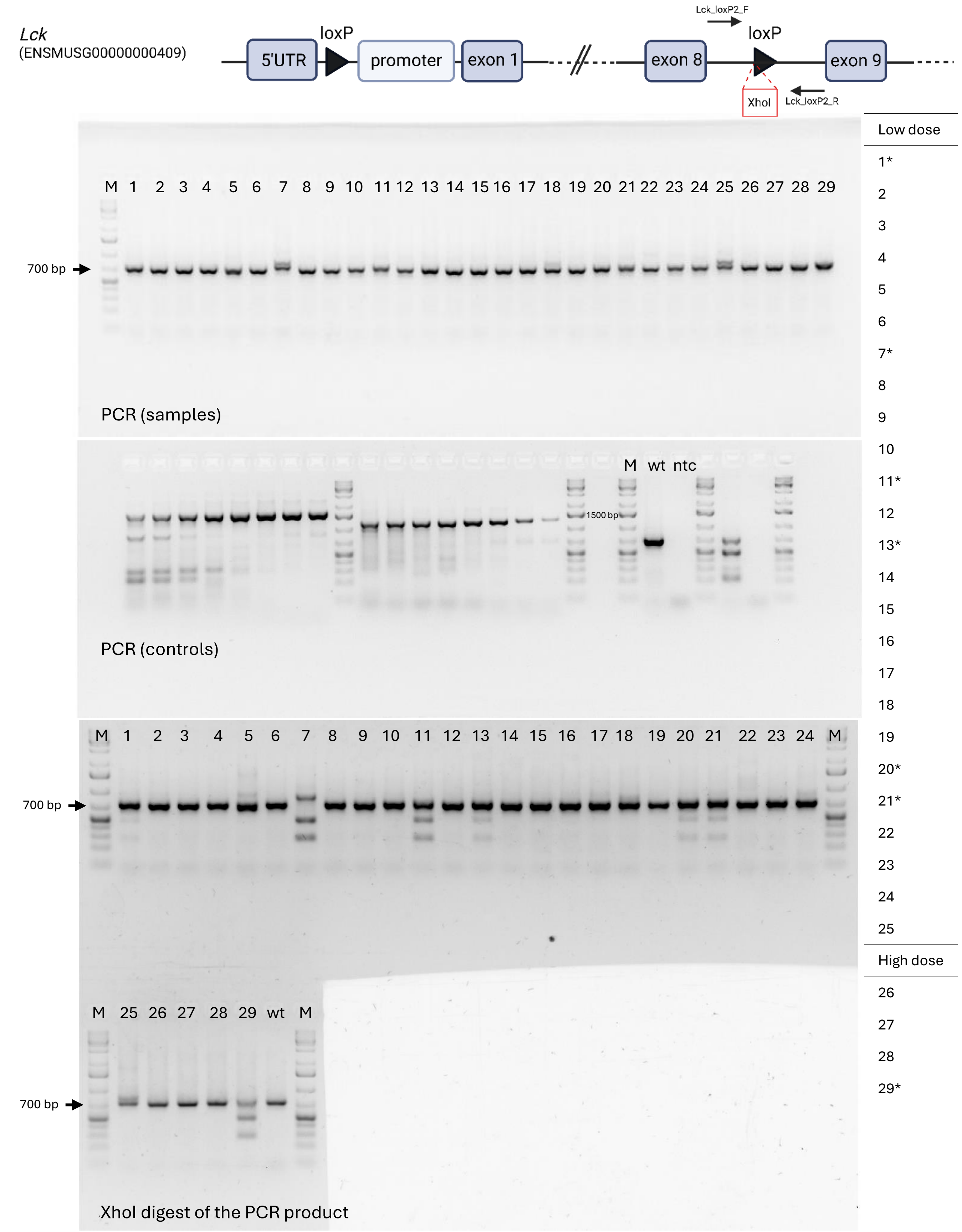


| PCR reaction | Primers sequence (5'>3') | Product size (bp) |
|---|---|---|
| Lck_loxP2_F | TTGCTGACAAGCCTGATGAGC | 713 (wt), 753 (+loxP) |
| Lck_loxP2_R | TGGGAACATCCCTAGGTCACAA | |
| **XhoI digestion** | WT product – no digestion, Lck cKO loxP2 – 274 + 479 bp fragments | |

D) C57BL/6NCrl-Actn1$^{em1Ccpcz}$

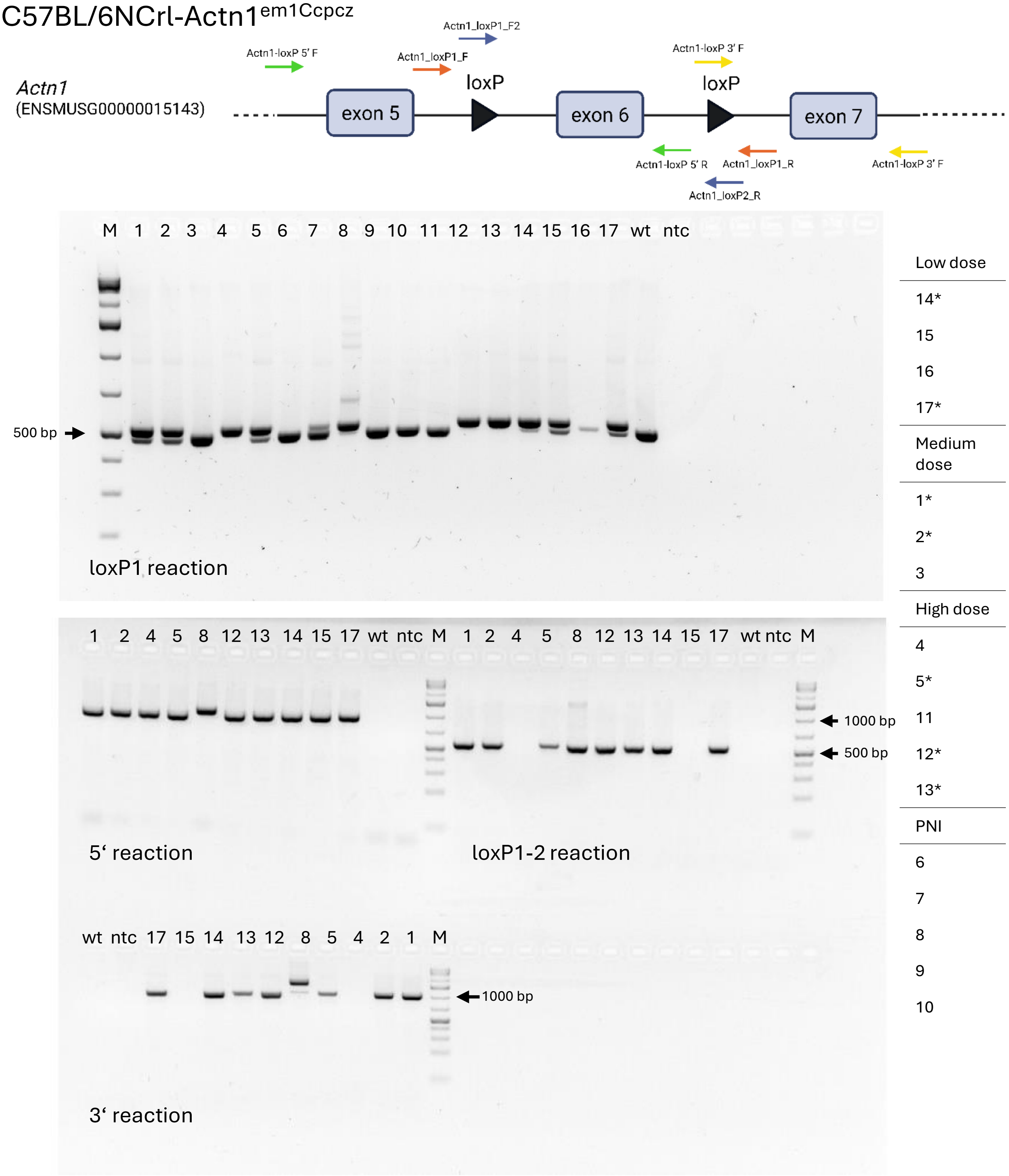


| **loxP1 reaction** | **Primers sequence (5'>3')** | **Product size (bp)** |
|---|---|---|
| Actn1_loxP1_F | ACACTGTCAAGACCCCCAAG | 420 (wt)/524 (loxP1) |
| Actn1_loxP1_R | TGTAGCTGGTCCATAAGGCCAA | |
| **5' reaction** | **Primers sequence (5'>3')** | **Product size (bp)** |
| Actn1-loxP 5' F | AGTTCACAGATCTGAACACACG | 874 |
| Actn1-loxP 5' R | ATTGAGCAAGAGAGGACAGC | |
| **3' reaction** | **Primers sequence (5'>3')** | **Product size (bp)** |
| Actn1-loxP 3' F | TGTTACCCAGCGATAACTTCG | 1060 |
| Actn1-loxP 3' F | ACATATGAGACAGTCTGTGAC | |
| **loxP1-2 reaction** | **Primers sequence (5'>3')** | **Product size (bp)** |
| Actn1_loxP1_F2 | GCTTTAGTGAACGATAACTTCG | 559 |
| Actn1_loxP2_R | CGGGATGTGCAATAACTTCGT | |

E) C57BL/6NCrl-Nes$^{em1(rtTA3,DTR,iRFP670)Ccpcz}$

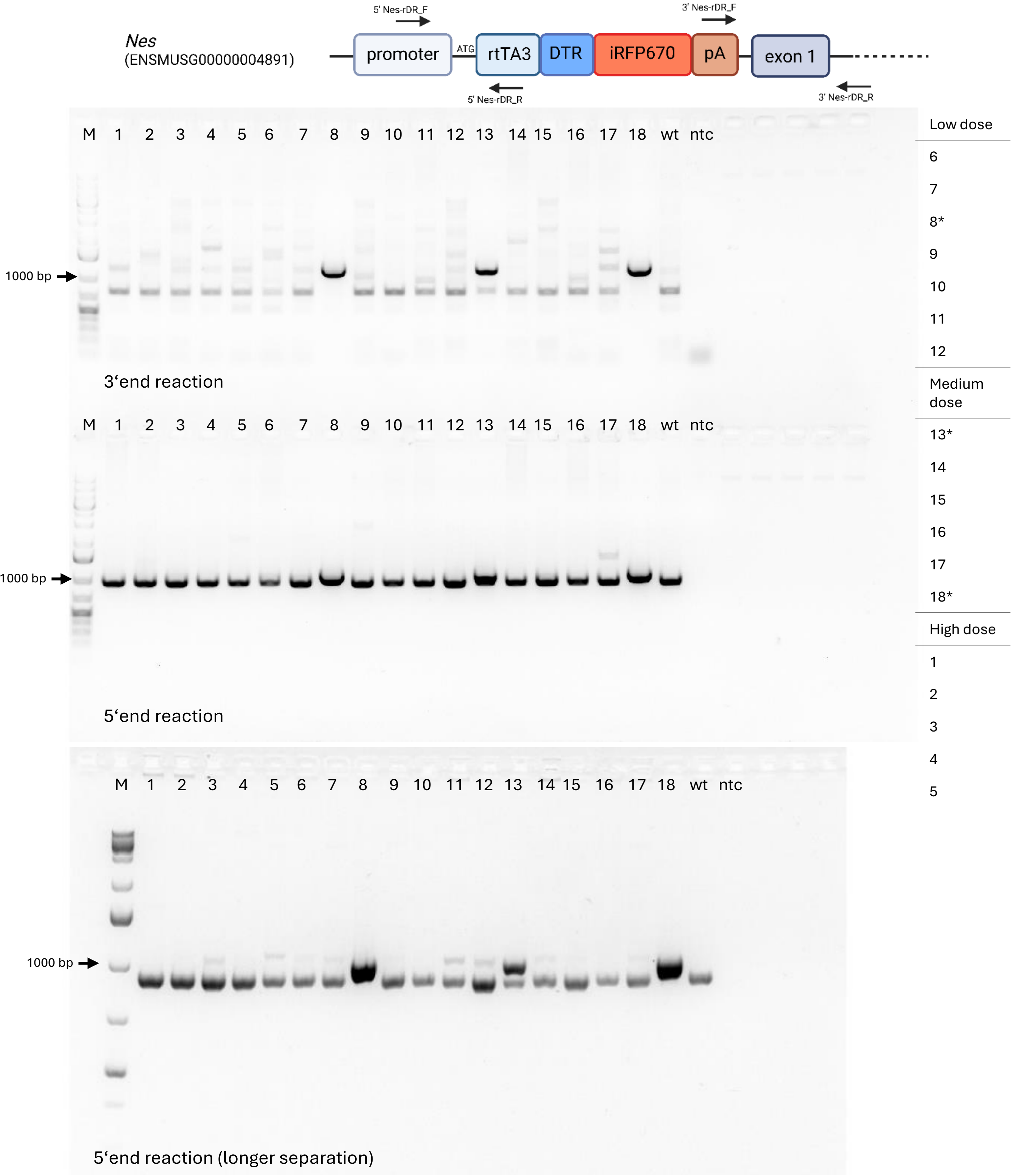


| 3‘reaction | Primers sequence (5‘>3‘) | Product size (bp) |
|---|---|---|
| 3' Nes-rDR_F | TGCGGGGTCTATTGGGAAC | 1154 |
| 3' Nes-rDR_R | CTGAAGGTCTCTTGCCATCCT | |
| **5‘reaction** | **Primers sequence (5‘>3‘)** | **Product size (bp)** |
| 5' Nes-rDR_F | AGCCGCGTAACTTCTTCACT | 1026 |
| 5' Nes-rDR_R | AACTCCCAGCTTTTGAGCGA | |

#### F) C57BL/6NCrl-Dpp4$^{em1(rtTA3,DTR,mKate)Ccpcz}$

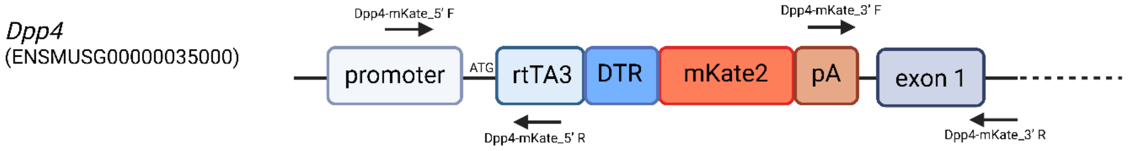


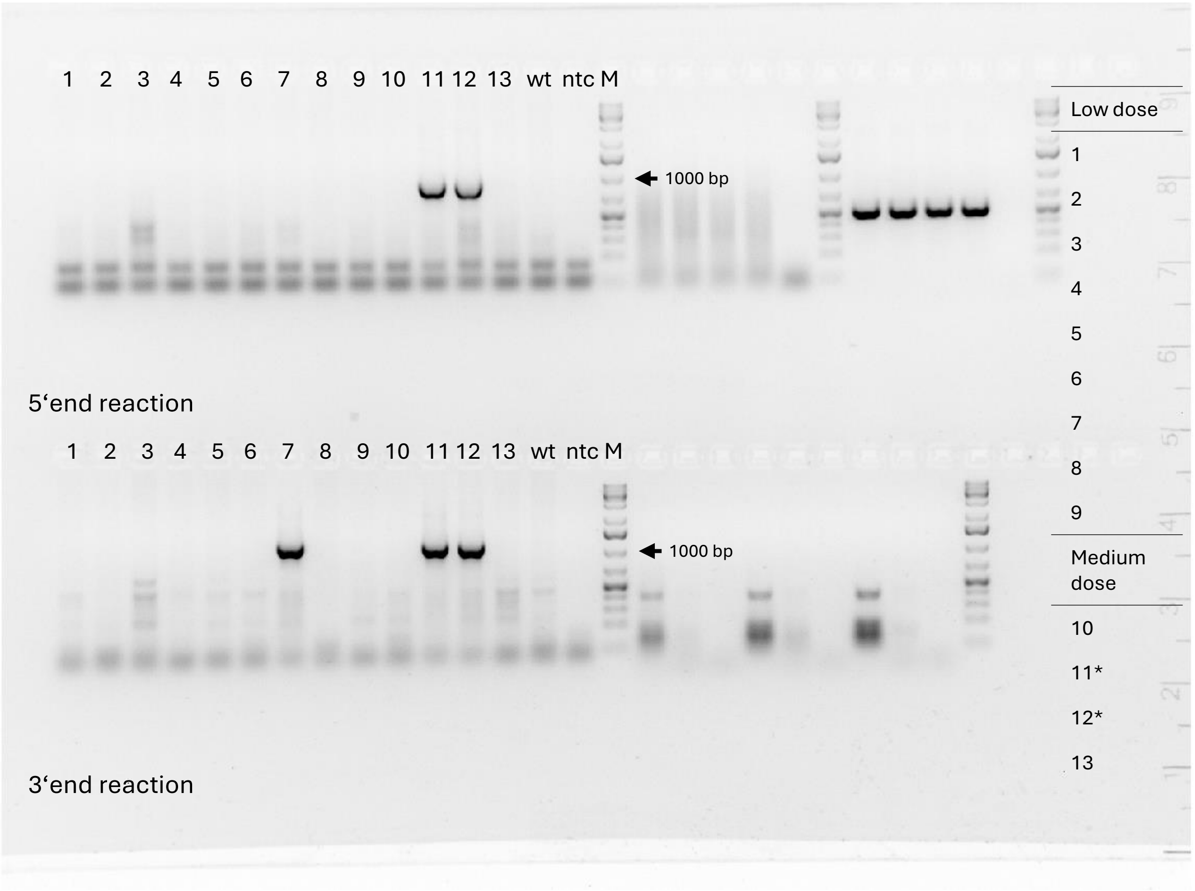


| 3'reaction | Primers sequence (5'>3') | Product size (bp) |
|---|---|---|
| Dpp4-mKate_3' F | CTGCGGGGTCTATTGGGAAC | 1213 |
| Dpp4-mKate_3' R | TGGGTCTTCAAAAGCTGCCT | |
| **5'reaction** | **Primers sequence (5'>3')** | **Product size (bp)** |
| Dpp4-mKate_5' F | CCTCTCAGGGAAGGGGACAAG | 938 |
| Dpp4-mKate_5' R | GAAGTGGGGGCATAGAATCGG | |

#### G) C57BL/6NCrl-Batf3$^{em1(iCre,eGFP)Ccpcz}$

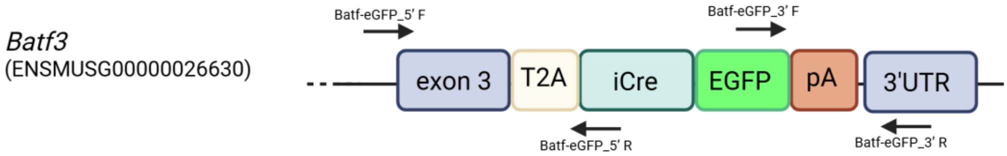


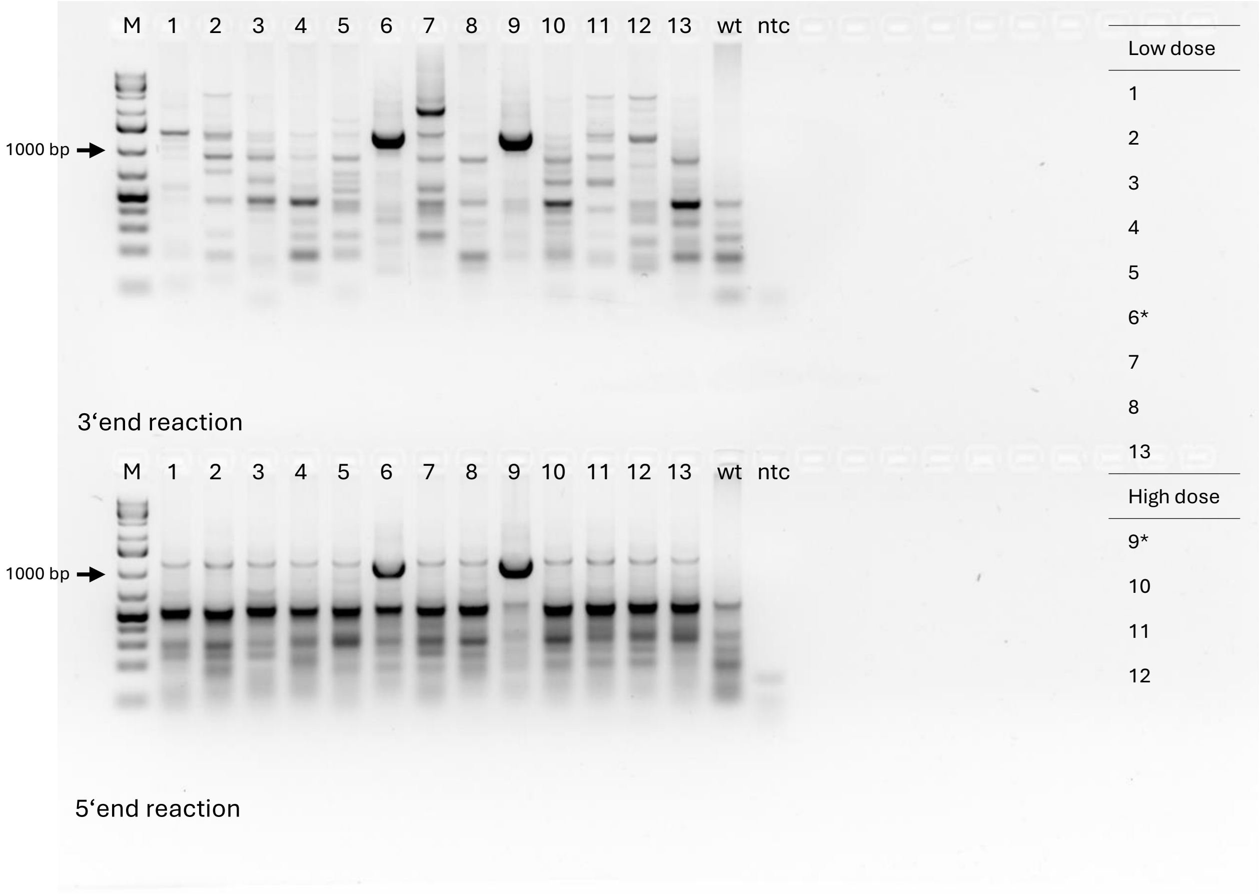


| 3'reaction | Primers sequence (5'>3') | Product size (bp) |
|---|---|---|
| Batf-eGFP_3' F | CACATGGTCCTGCTGGAGTT | 1350 |
| Batf-eGFP_3' R | TGCAAGAAGAATGGGCACCT | |
| **5'reaction** | **Primers sequence (5'>3')** | **Product size (bp)** |
| Batf-eGFP_5' F | GAACTTGTAAGGGCGAGGGA | 1103 |
| Batf-eGFP_5' R | ATCAGAGGTGGCATCCACAG | |

#### H) C57BL/6NCrl-Alox5ap$^{em1(CCre, CGFP)Ccpcz}$

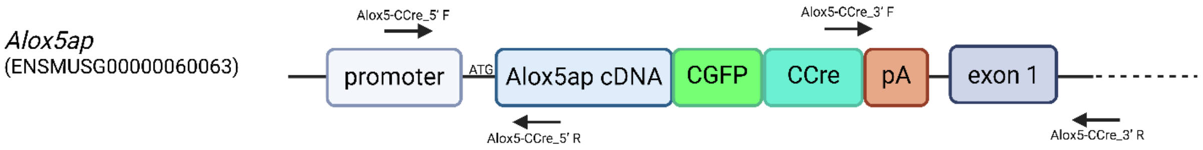


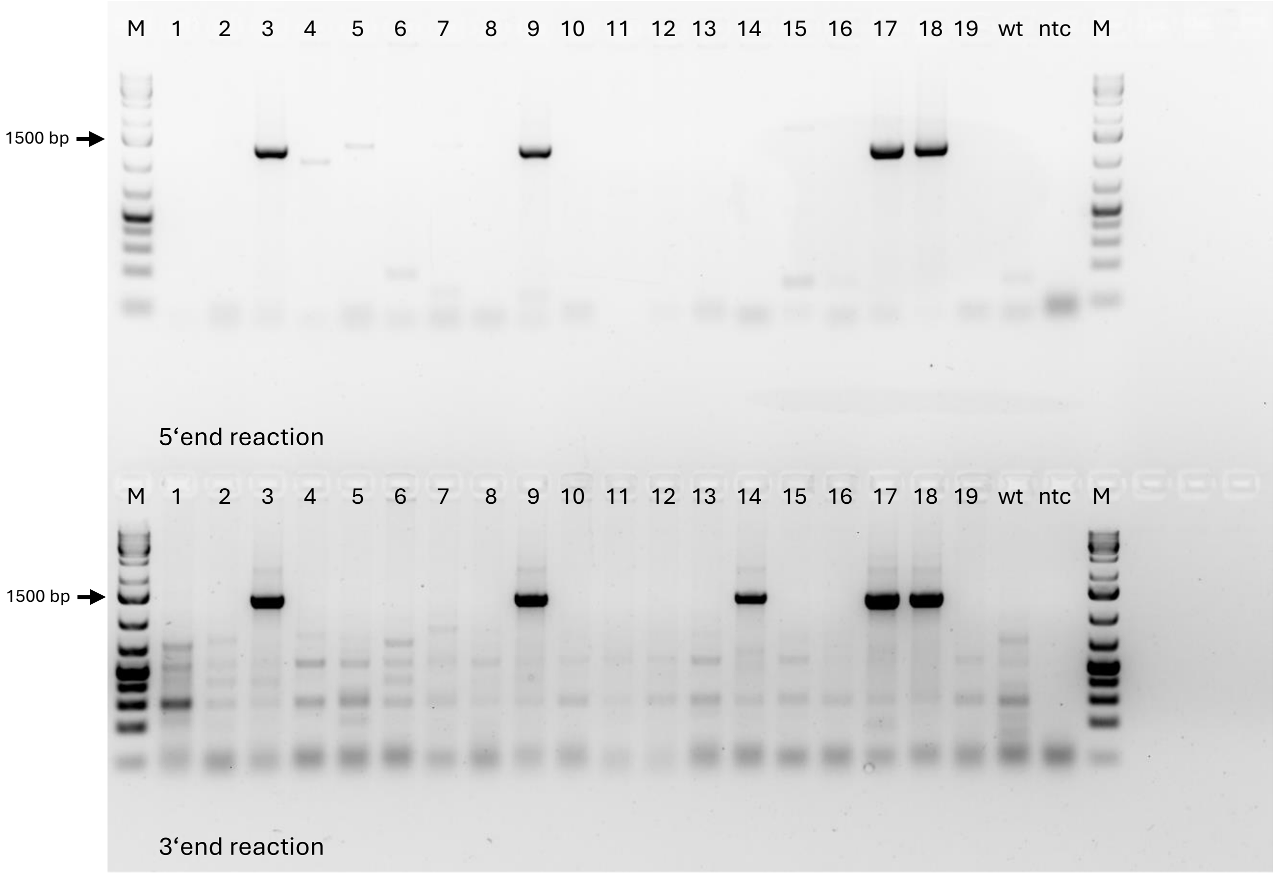


| Low dose |
|---|
| 1 |
| 2 |
| 3* |
| 4 |
| 5 |
| Medium dose |
| 15 |
| 14 |
| 15 |
| 16 |
| 17* |
| 18* |
| 19 |
| High dose |
| 6 |
| 7 |
| 8 |
| 9* |
| 10 |
| 11 |
| 12 |
| 13 |
| 14 |

| **3'reaction** | **Primers sequence (5'>3')** | **Product size (bp)** |
|---|---|---|
| Alox5-CCre_3' F | CTCGAGGATGGGGACTGATG | 1358 |
| Alox5-CCre_3' R | ACTGTGTCAAAGGGCTAGGA | |
| **5'reaction** | **Primers sequence (5'>3')** | **Product size (bp)** |
| Alox5-CCre_5' F | CCTGCCCCTCTTCCTGTAAT | 1340 |
| Alox5-CCre_5' R | GGCAAAGAACGCATTCTGGA | |

### I) C57BL/6NCrl-Lyve$^{em1(NCre,\ NGFP)Ccpcz}$

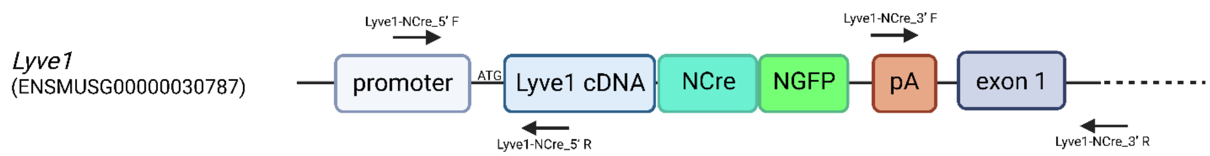


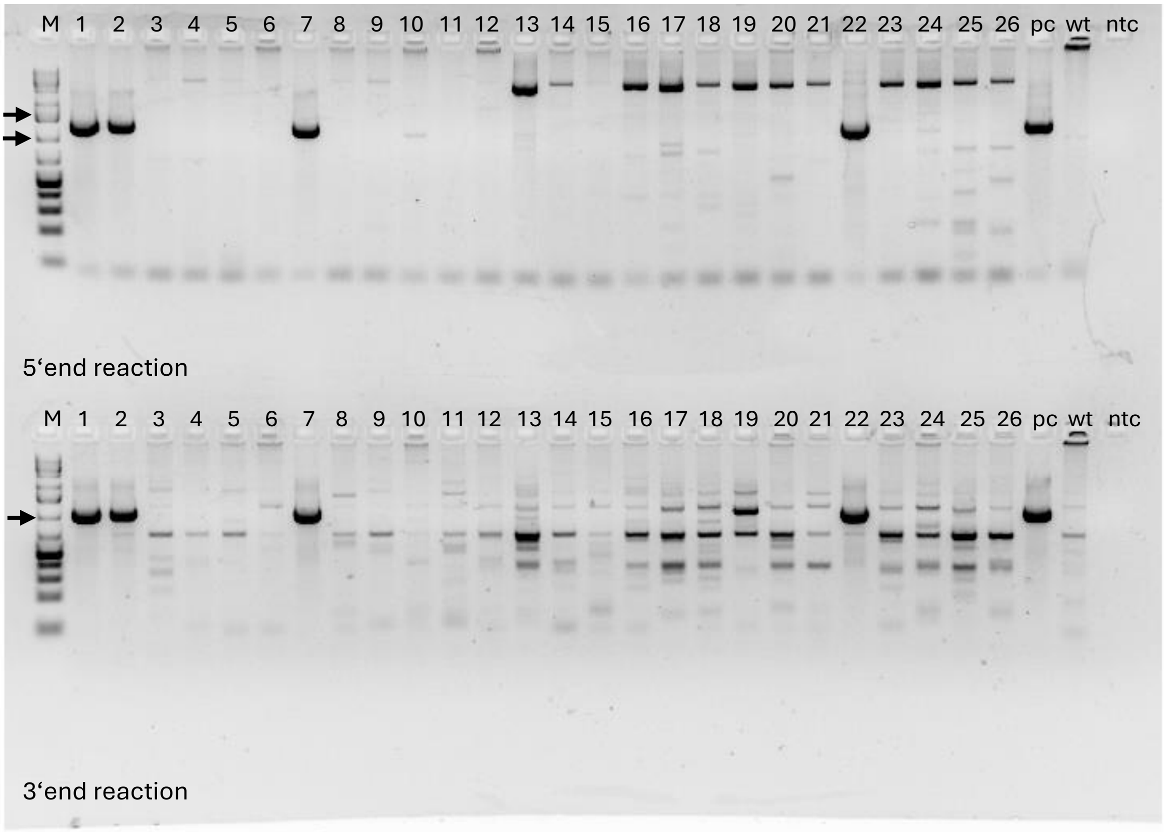


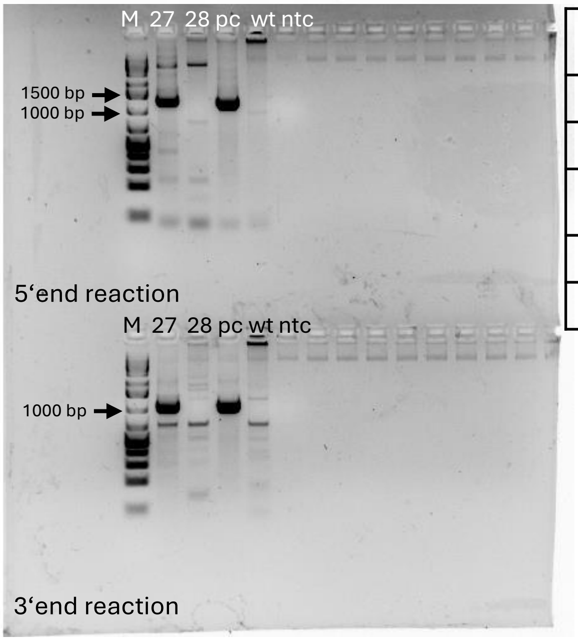


| 3reaction | Primers sequence (5‘>3‘) | Product size (bp) |
|---|---|---|
| Lyve1-NCre_3’ F | TGGTAGAGACGGGGTTTCAC | 1061 |
| Lyve1-NCre_3’ R | TGGCATTGCTAAGAGGACCA | |
| **5‘reaction** | **Primers sequence (5‘>3‘)** | **Product size (bp)** |
| Lyve1-NCre_5’ F | CTCCAGGCGTTACTCCAGAG | 1297 |
| Lyve1-NCre_5’ R | GCCTCGTTGGCTTCTGTGA | |

Low dose

10
11
12
13
14
15
27*
28

High dose

1*
2*
3
4

Medium dose

5
6
7*
8
9
16
17
18
19
20
21
22*
23
24
25
26

#### J) C57BL/6NCrl-Sox2$^{em1(P2A\text{-}mStrawberry)Ccpcz}$

*Sox2*
(ENSMUSG00000074637)

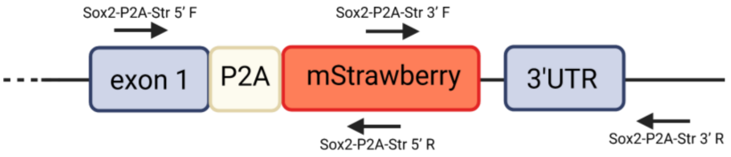


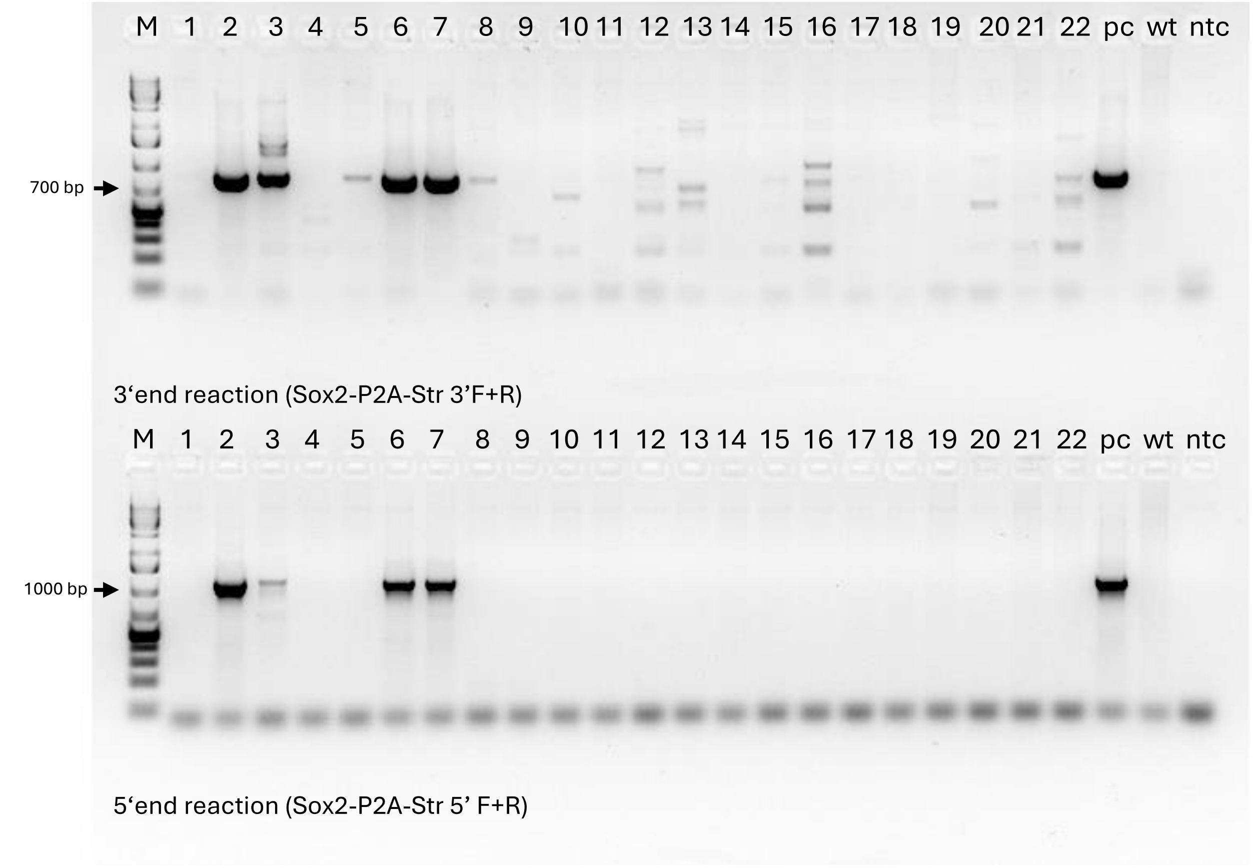


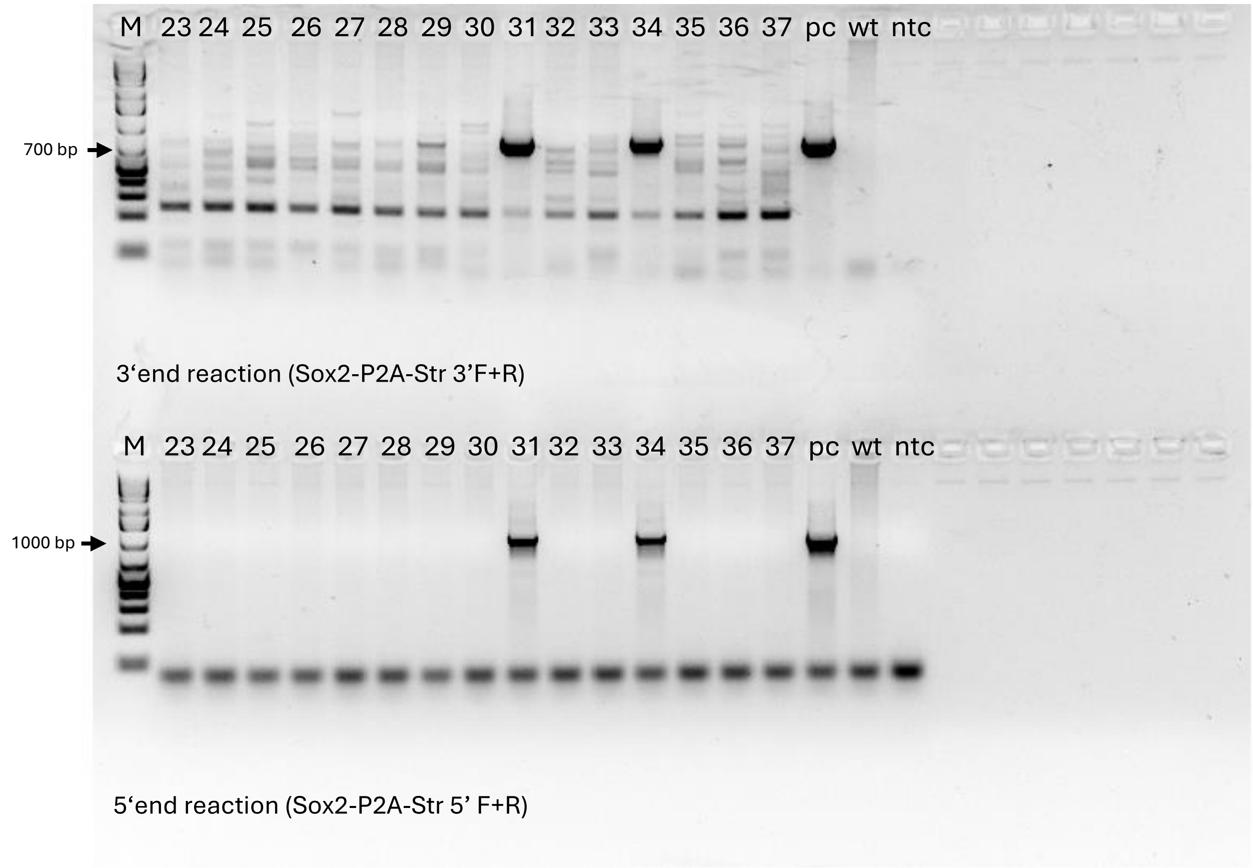


| |
|---|
| AAV Medium dose |
| 1 |
| 2 |
| 3* |
| 4* |
| AAV-EV Low dose |
| 5 |
| 6* |
| 7* |
| 8 |
| 9 |
| 10 |
| 11 |
| 12 |
| 13 |
| 14 |
| 15 |
| 16 |
| 35 |
| 36 |
| 37 |
| AAV-EV Medium dose |
| 17 |
| 18 |
| 19 |
| 20 |
| 21 |
| 22 |
| 29 |
| 30 |
| 31* |
| 32 |
| 33 |
| 34* |
| AAV-EV High dose |
| 25 |
| 26 |
| 27 |
| 28 |

#### K) C57BL/6NCrl-Alb$^{em6(Cyp3a4,P2A,Ces1)Ccpcz}$

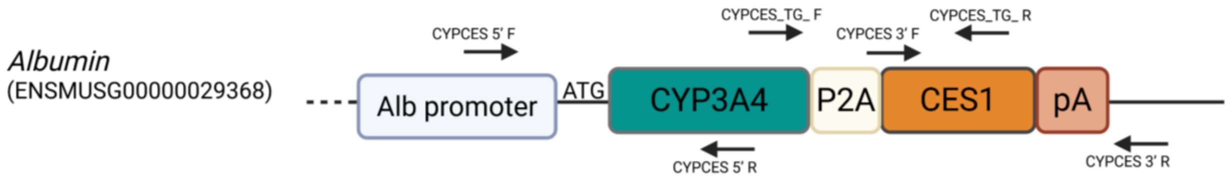


| 3‘reaction | Primers sequence (5‘>3‘) | Product size (bp) |
|---|---|---|
| CYPCES 3’F | | 836 |
| CYPCES 3’R | | |
| **5‘reaction** | **Primers sequence (5‘>3‘)** | **Product size (bp)** |
| CYPCES 5’F | CACCCCGAGAAAGAGGTTCA | 689 |
| CYPCES 5’R | GCCAAGTCTGGGATGAGAGC | |
| **transgene** | **Primers sequence (5‘>3‘)** | **Product size (bp)** |
| CYPCES_TG_F | CAAGGGATGGCACCGTAAGT | 1420 |
| CYPCES_TG_R | GCTCCAGCATCTCTGTGGTT | |

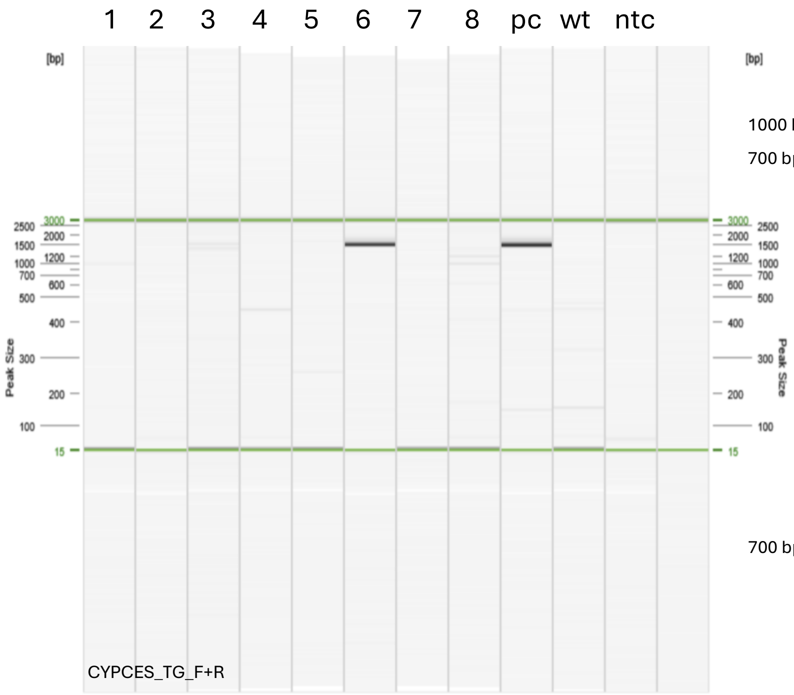


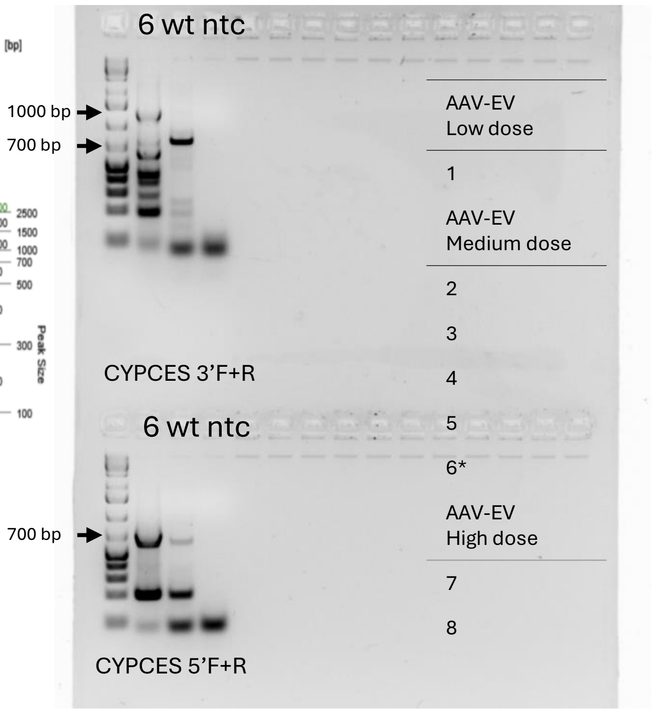


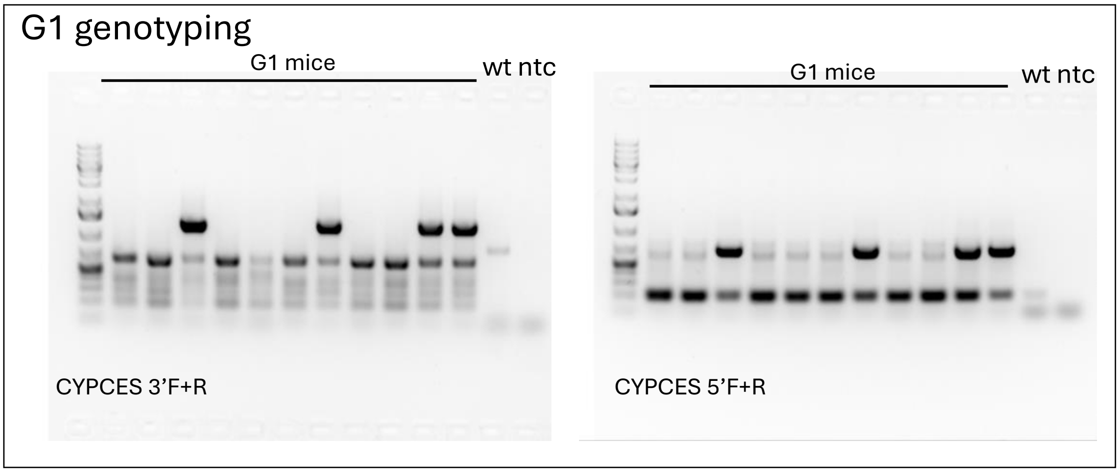

L) SD-Umod$^{em1(C126R)Ccpcz}$

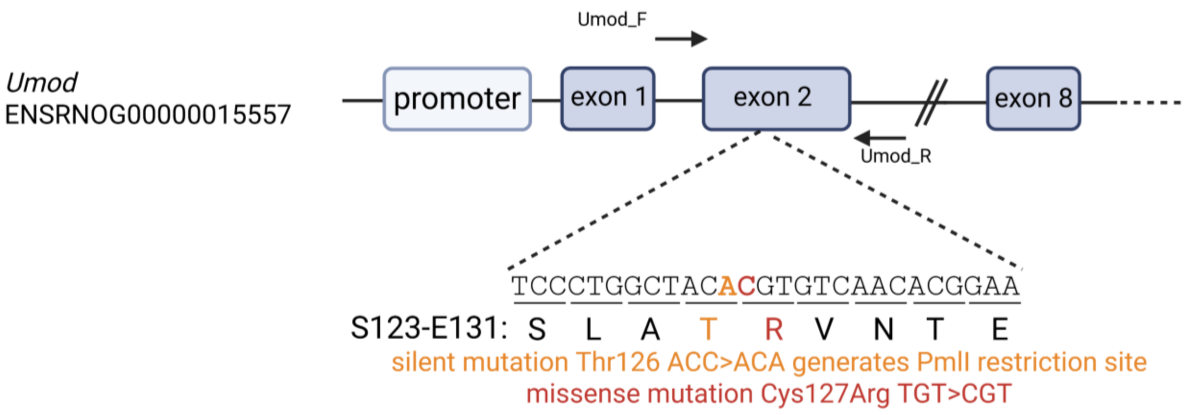


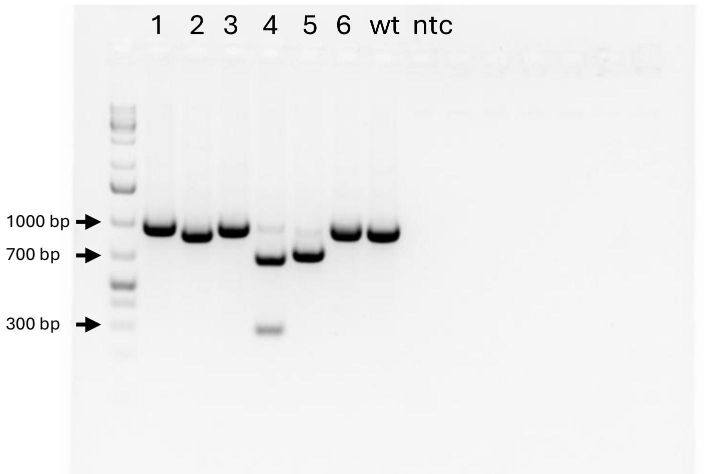

Umod C127R F+R PCR product digestion with PmlI

| AAV-EV Low dose |
|---|
| 1 |
| 2 |
| 3 |
| 4** |
| 5 |
| 6 |

** genotype confirmed by Sanger sequencing

| PCR reaction | Primers sequence (5‘>3‘) | Product size (bp) |
|---|---|---|
| Umod C127R F | AGGTGTTCTGAATGCCACGA | 993 |
| Umod C127R R | ATCTCCCTCTGAGTCTCACCT | |
| **PmlI digestion** | WT product – no digestion, C127R positive product – 293 + 710 bp fragments | |

#### M) SD-$Umod^{em1(DelY178-R186)Ccpcz}$

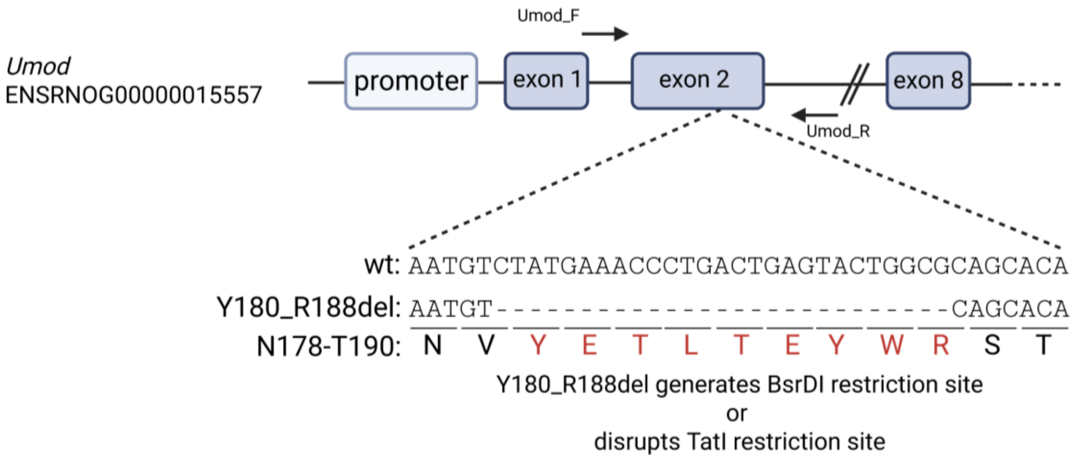


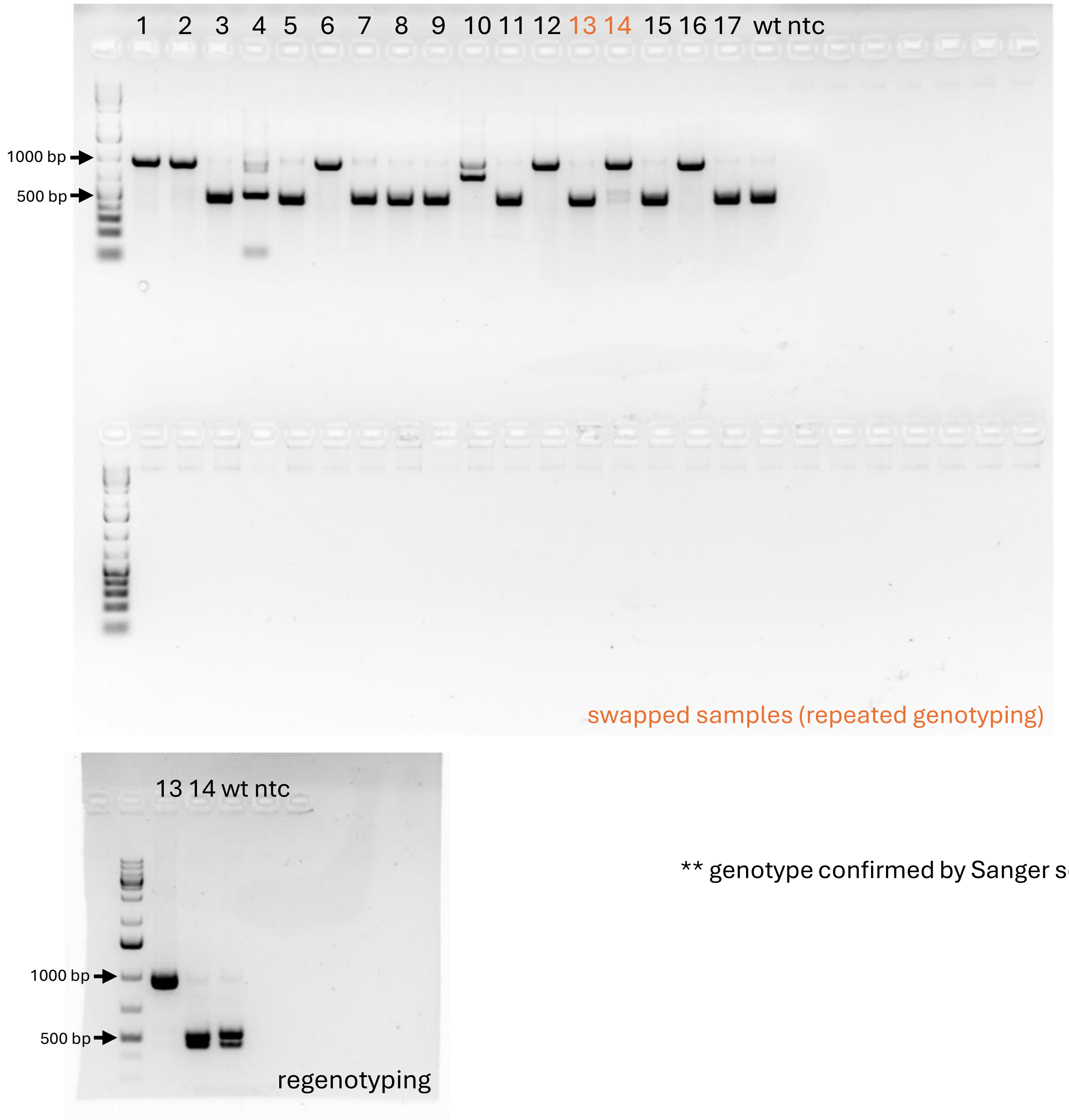


AAV-EV Low dose

16

17

AAV-EV Medium dose

1**

2

3

4

5

6**

7

8

9

10

11

12

13**

14

15

** genotype confirmed by Sanger sequencing

| PCR reaction | Primers sequence (5'>3') | Product size (bp) |
|---|---|---|
| Umod YR F | AGGTGTTCTGAATGCCACGA | 993 |
| Umod YR R | ATCTCCCTCTGAGTCTCACCT | |
| **PmlI digestion** | WT product – after TatI digestion: 458+532bp, Y180-R188 deletion positive product: 993 bp | |

#### N) $SD\text{-}Muc1^{em1(MUC1wt)Ccpcz}$

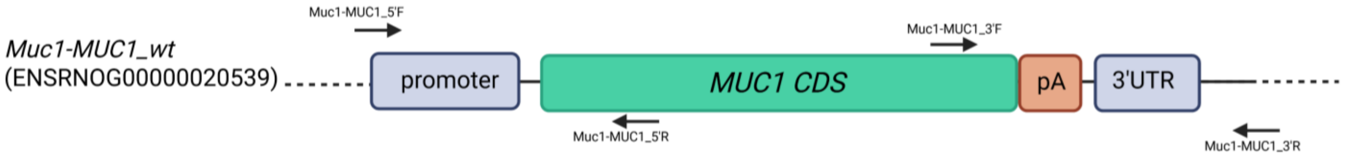


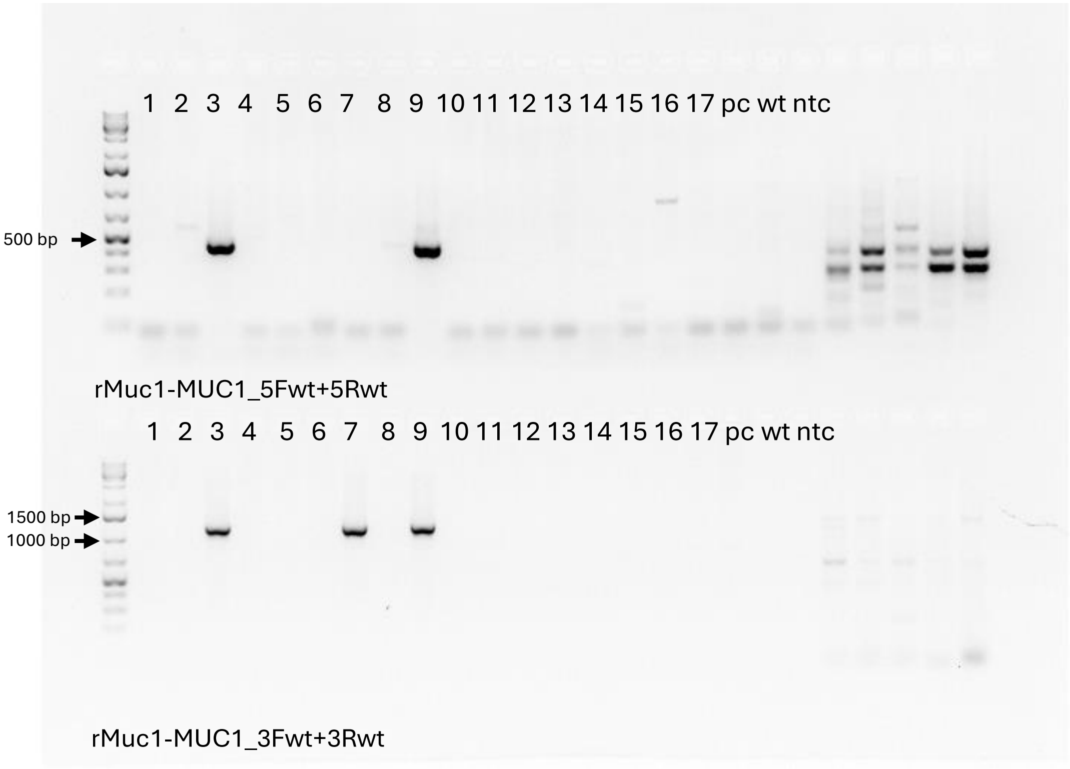


| AAV-EV Low dose |
|---|
| 1 |
| 2 |
| 3* |
| 4 |
| 5 |
| 6 |
| 7 |
| 8 |
| 9* |
| 10 |
| 11 |
| 12 |
| 13 |
| 14 |
| 15 |
| 16 |
| 17 |

| 3'reaction | Primers sequence (5'>3') | Product size (bp) |
|---|---|---|
| rMuc1-MUC1_3Fwt | TTTCCCATTTTCCGCCCAGT | 1160 |
| rMuc1-MUC1_3Rwt | ACAGTTATGGCGCGAGTTGA | |
| **5'reaction** | **Primers sequence (5'>3')** | **Product size (bp)** |
| rMuc1-MUC1 5Fwt | TGGGGTACTCAAAGAGTGGTTG | 446 |
| rMuc1-MUC1 5Rwt | CATTCTTCTCGGTAGAGGATGGC | |

O) *SD-Muc1*$^{em2(MUC1fs)Ccpcz}$

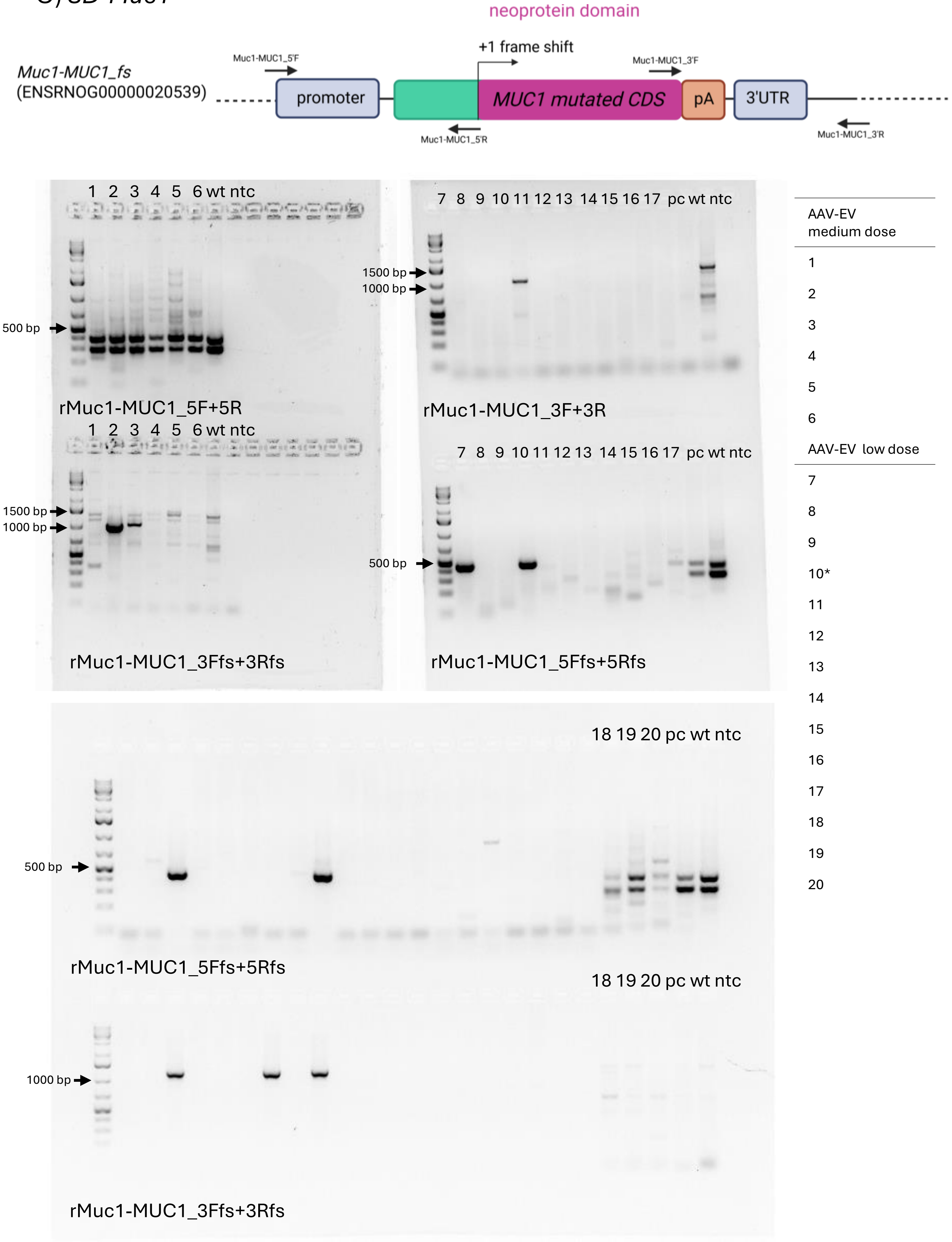


| 3'reaction | Primers sequence (5'>3') | Product size (bp) | 5'reaction | Primers sequence (5'>3') | Product size (bp) |
|---|---|---|---|---|---|
| rMuc1-MUC1_3Ffs | CTTTTATAGCGGCGCCCAGA | 1160 | rMuc1-MUC1 5Ffs | TGGGGTACTCAAAGAGTGGTTG | 446 |
| rMuc1-MUC1_3Rfs | CACACCTGGCCAGCTAAGAAT | | rMuc1-MUC1 5Rfs | CGTTCTTCTCGGTGGAGCTTGGC | |

**Additional file 8 figure: Detection of transgenic cassette in a corresponding rodent model**

(A) C57BL/6NCrl-Ube3a$^{em2(BioID2)Ccpcz}$ mouse line: site-specific insertion of Ube3a-BioID2 cassette in the genome detected at the 5' and 3' end. (B) C57BL/6NCrl-Sox2$^{em1(P2A-mStrawberry)Ccpcz}$ mouse line: site-specific insertion of Sox2-mStrawberry cassette in the genome detected at the 5' and 3' end.(C) C57BL/6NCrl-Lck$^{em2Ccpcz}$ mouse line: loxP insertion detected via XhoI digestion of the PCR product. (D) C57BL/6NCrl-Actn1$^{em1Ccpcz}$ mouse line: site-specific insertion of Actn1 cKO cassette in the genome detected at the 5' and 3' end; detection of loxP1 (loxP1 reaction) and of both loxP sites on the same allele (loxP1-2 reaction). (E) C57BL/6NCrl-Nes$^{em1(rtTA3,DTR,iRFP670)Ccpcz}$ mouse line: site-specific insertion of Nes-rtTA/DTR/iRFP670 cassette in the genome detected at the 5' and 3' end.(F) C57BL/6NCrl-Dpp4$^{em1(rtTA3,DTR,mKate)Ccpcz}$ mouse model: site-specific insertion of Dpp4-rtTA3-DTR-mKate2 cassette in the genome detected at the 5' and 3' end.(G) ) C57BL/6NCrl-Batf3$^{em1(iCre,eGFP)Ccpcz}$ mouse model: site-specific insertion of Batf3-iCre-eGFP cassette in the genome detected detection at the 5' and 3' end.(H) C57BL/6NCrl-Alox5ap$^{em1(CCre, CGFP)Ccpcz}$ mouse model: site-specific insertion of Alox5ap-CCreCGFP cassette in the genome detected detection at the 5' and 3' end.(I) C57BL/6NCrl-Lyve$^{em1(NCre, NGFP)Ccpcz}$ mouse line: site-specific of Lyve1-NCre-NGFP cassette insertion in the genome detected detection at the 5' and 3' end. (J) Genome-transgene junction genotyping of C57BL/6NCrl-Sox2$^{em1(P2A-mStrawberry)Ccpcz}$ animals targeted with Sox-mStr construct using AAVs and distinct doses of AAV-EV vector prepared with UCF-method. (K) Genome-transgene junction genotyping of C57BL/6NCrl-Alb$^{em6(Cyp3a4,P2A,Ces1)Ccpcz}$ animals targeted with Alb-CYP3A4-CES1 construct using AAVs and distinct doses of AAV-EV vector prepared with UCF-method. (L) SD-Umod$^{em1(C126R)Ccpcz}$ rat line: C127R insertion detected via PmlI digestion of the PCR product and Sanger sequencing. (M) SD-Umod$^{em1(DelY178-R186)Ccpcz}$ rat line: Y180-R188 deletion confirmed by loss of TatI restriction site and Sanger sequencing. (N) *SD-Muc1*$^{em1(MUC1wt)Ccpcz}$ rat line: site-specific insertion of Muc1-MUC1_wt (human MUC1 wild-type coding sequence) cassette in the genome detected cassette detection at the 5' and 3' end. (O) *SD-Muc1*$^{em2(MUC1fs)Ccpcz}$rat line: site-specific insertion of Muc1-MUC1_fs (human MUC1 coding sequence with +1 frame shift) cassette in the genome detected cassette detection at the 5' and 3' end.

Positive animals are marked by an asterisk (*) in the side column, which also summarizes the number of animals per treated group. Positive animals confirmed by sanger sequencing are marked by a double asterick (**). Tables summarize used proteins for corresponding reaction.

### Table: List of gRNA spacers used in rodent model generation

| AAV-EV | Model | Spacer (5'-3') |
|---|---|---|
| Ube3a-BioID2 | C57BL/6NCrl-Ube3a$^{\text{em2(BioID2)Ccpcz}}$ | ATCACATATGCCAAAGGATT |
| Sox2-mStrawberry | C57BL/6NCrl-Sox2$^{\text{em1(P2A-mStrawberry)Ccpcz}}$ | CTGCCCCTGTCGCACATGTG |
| Lck loxP2 | C57BL/6NCrl-Lck$^{\text{em2Ccpcz}}$ | GGAGTGCAAATCTTCCAGGA |
| Actn1 cKO | C57BL/6NCrl-Actn1$^{\text{em1Ccpcz}}$ | TAAGGCTTTAGTGAACGGGT |
| | | GCGTGCACATCCCGGGTCCC |
| Nes-rtTA3-DTR-IRES-iRFP670 | C57BL/6NCrl-Nes$^{\text{em1(rtTA3,DTR,iRFP670)Ccpcz}}$ | GACGCAACCCTCCATGTCGC |
| Dpp4-rtTA3-DTR-H1-mKate2 | C57BL/6NCrl-Dpp4$^{\text{em1(rtTA3,DTR,mKate)Ccpcz}}$ | AAGGAGCCGCCCGACCATGA |
| Batf3-iCre-eGFP | C57BL/6NCrl-Batf3$^{\text{em1(iCre,eGFP)Ccpcz}}$ | AGTGAGCTGGGGTGTCATCG |
| Alox5ap-CCre-CGFP | C57BL/6NCrl-Alox5ap$^{\text{em1(CCre, CGFP)Ccpcz}}$ | TGGCGGGCCTGAAGCAAGCA |
| Lyve1NCre-NGFP | C57BL/6NCrl-Lyve$^{\text{em1(NCre, NGFP)Ccpcz}}$ | GTAACACCAGGCTAGTGTGC |
| Alb-CYP3A4-CES1 | C57BL/6NCrl-Alb$^{\text{em6(Cyp3a4,P2A,Ces1)Ccpcz}}$ | AGCCTCTGGCAAAATGAAGT |
| | | GTTGTGATGTGTTTAGGCTA |
| Umod C127R | SD-Umod$^{\text{em1(C126R)Ccpcz}}$ | GCTGCCTTCCGTGTTGACAC |
| Umod Y180_R188del | SD-Umod$^{\text{em1(DelY178-R186)Ccpcz}}$ | GGGTTTCATAGACATTGCAG |
| | | TACTGGCGCAGCACAGACTA |
| Muc1-MUC1_wt/fs | *SD-Muc1*$^{\textit{em1(MUC1)Ccpcz}}$ *SD-Muc1*$^{\textit{em2(MUC1fs)Ccpcz}}$ | AACCCGTATGCCCGGGGTCA |
| | | CTCCTACAAGTTGGCCGAAG |
