## Additional file 9 for "Adeno-Associated Virus Co-Precipitation with Extracellular Vesicles for Genome Editing in Rodent Embryo": AAV-EGFP transposon.pdf

### NANOSIGHT

#### AAV MPRA GFP 1000x 5 60 120 2024-12-18 12-35-07

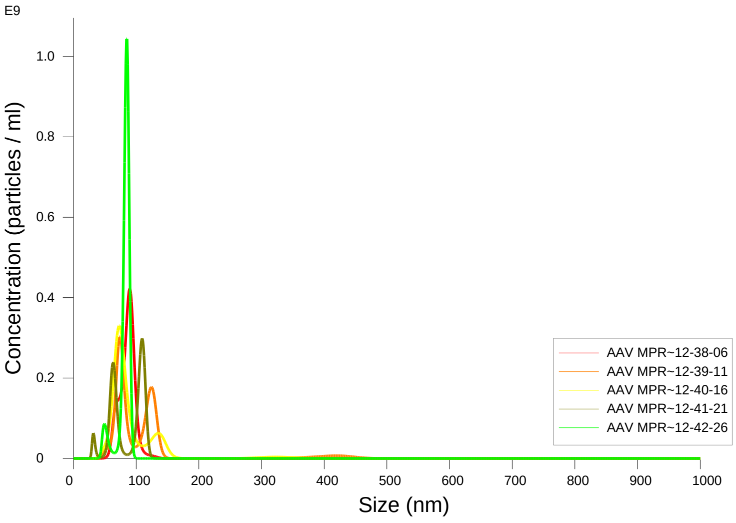


FTLA Concentration / Size graph for Experiment:
AAV MPRA GFP 1000x 5 60 120 2024-12-18 12-35-07

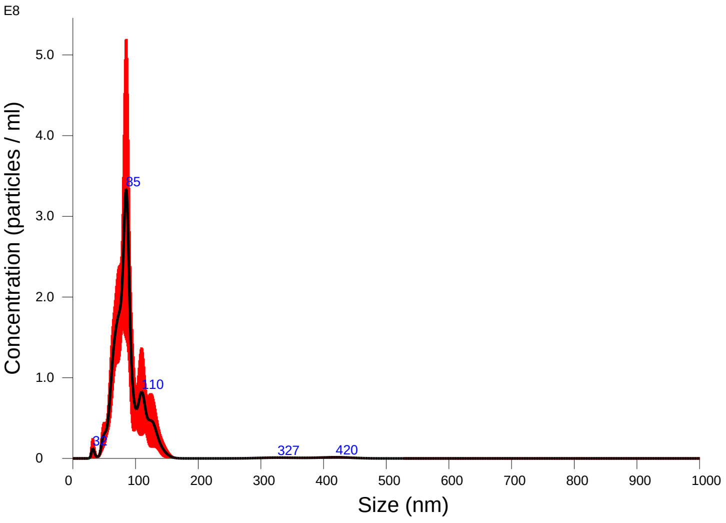


Averaged FTLA Concentration / Size for Experiment:
AAV MPRA GFP 1000x 5 60 120 2024-12-18 12-35-07
Error bars indicate + / -1 standard error of the mean

**Included Files**

AAV MPRA GFP 1000x 5 60 120 2024-12-18 12-38-06
AAV MPRA GFP 1000x 5 60 120 2024-12-18 12-39-11
AAV MPRA GFP 1000x 5 60 120 2024-12-18 12-40-16
AAV MPRA GFP 1000x 5 60 120 2024-12-18 12-41-21
AAV MPRA GFP 1000x 5 60 120 2024-12-18 12-42-26

**Details**

| | |
|---|---|
| NTA Version: | NTA 3.4 - Sample Assistant Build 3.4.4 - SA |
| Script Used: | SOP Standard Measurement 12-34-44PM 18~ |
| Time Captured: | 12:35:07 18/12/2024 |
| Operator: | Jitka |
| Pre-treatment: | |
| Sample Name: | |
| Diluent: | pbs |
| Remarks: | |

**Capture Settings**

| | |
|---|---|
| Camera Type: | sCMOS |
| Laser Type: | Blue405 |
| Camera Level: | 12 |
| Slider Shutter: | 1200 |
| Slider Gain: | 146 |
| FPS | 25.0 |
| Number of Frames: | 1498 |
| Temperature: | 25.0 - 25.0 ºC |
| Viscosity: | 1.0 cP |
| Dilution factor: | 1 x 10e3 |
| Syringe Pump Speed: | 120 |

**Analysis Settings**

| | |
|---|---|
| Detect Threshold: | 5 |
| Blur Size: | Auto |
| Max Jump Distance: | Auto: 14.6 pix |

**Results**

Stats: Merged Data

| | |
|---|---|
| Mean: | 91.4 nm |
| Mode: | 84.9 nm |
| SD: | 43.0 nm |
| D10: | 63.2 nm |
| D50: | 84.3 nm |
| D90: | 120.5 nm |

Stats: Mean +/- Standard Error

| | |
|---|---|
| Mean: | 91.4 +/- 5.4 nm |
| Mode: | 86.3 +/- 6.6 nm |
| SD: | 33.5 +/- 12.2 nm |
| D10: | 65.7 +/- 2.7 nm |
| D50: | 86.8 +/- 3.3 nm |
| D90: | 114.5 +/- 9.2 nm |
| Concentration (Upgrade): | 1.04e+10 +/- 6.65e+08 particles/ml |
| | 0.7 +/- 0.1 particles/frame |
| | 1.0 +/- 0.1 centres/frame |

Concentration measurements may be unreliable
See summary file for more info

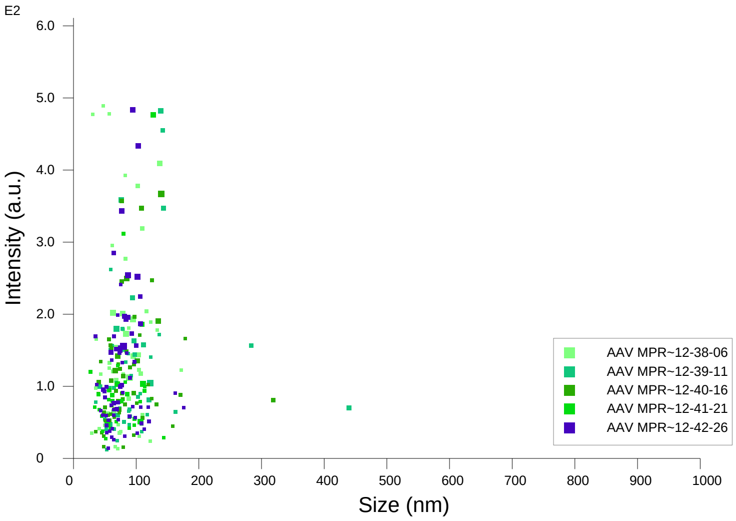


Intensity / Size graph for Experiment:
AAV MPRA GFP 1000x 5 60 120 2024-12-18 12-35-07

**Script Used: (Full Text):**

SOP Standard Measurement 12-34-44PM 18Dec2024.txt
