## Additional file 9 for "Adeno-Associated Virus Co-Precipitation with Extracellular Vesicles for Genome Editing in Rodent Embryo": AAV-Hypertransposase.pdf

NANOSIGHT

AAV Hypertransposase 1000x 5 60 120 2024-12-18 12-58-48

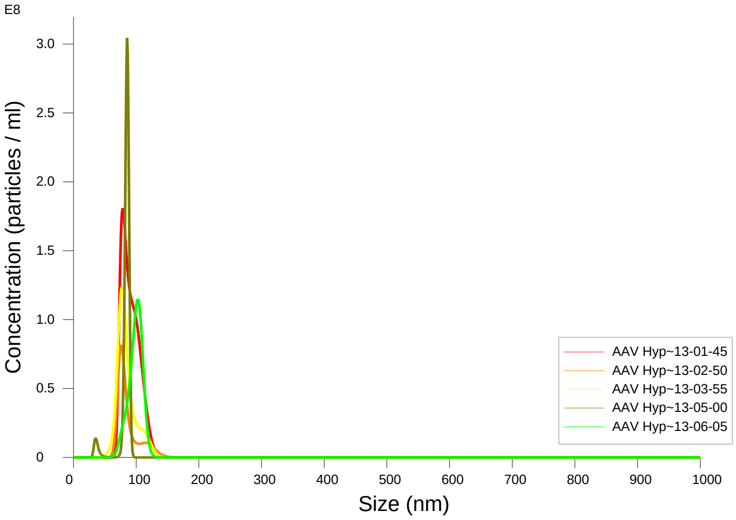


FTLA Concentration / Size graph for Experiment:
AAV Hypertransposase 1000x 5 60 120 2024-12-18 12-58-48

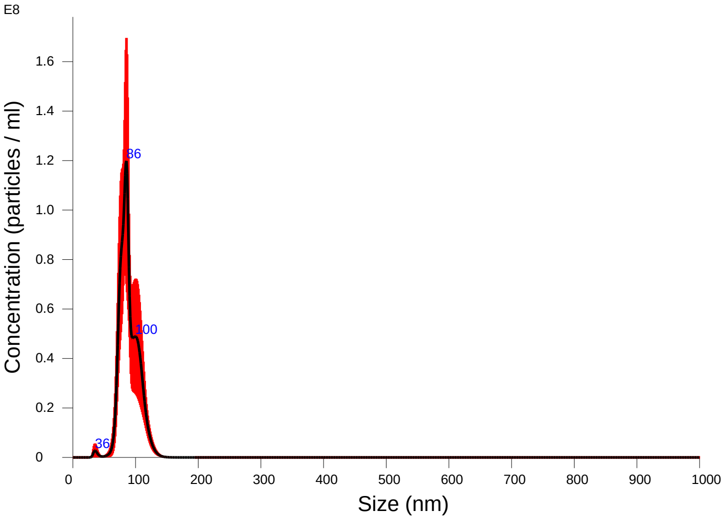


Averaged FTLA Concentration / Size for Experiment:
AAV Hypertransposase 1000x 5 60 120 2024-12-18 12-58-48
Error bars indicate + / -1 standard error of the mean

### Included Files

AAV Hypertransposase 1000x 5 60 120 2024-12-18 13-01-45
AAV Hypertransposase 1000x 5 60 120 2024-12-18 13-02-50
AAV Hypertransposase 1000x 5 60 120 2024-12-18 13-03-55
AAV Hypertransposase 1000x 5 60 120 2024-12-18 13-05-00
AAV Hypertransposase 1000x 5 60 120 2024-12-18 13-06-05

### Details

| | |
|---|---|
| NTA Version: | NTA 3.4 - Sample Assistant Build 3.4.4 - SA |
| Script Used: | SOP Standard Measurement 12-58-31PM 18~ |
| Time Captured: | 12:58:48 18/12/2024 |
| Operator: | Jitka |
| Pre-treatment: | |
| Sample Name: | |
| Diluent: | pbs |
| Remarks: | |

### Capture Settings

| | |
|---|---|
| Camera Type: | sCMOS |
| Laser Type: | Blue405 |
| Camera Level: | 12 |
| Slider Shutter: | 1200 |
| Slider Gain: | 146 |
| FPS | 25.0 |
| Number of Frames: | 1498 |
| Temperature: | 25.0 - 25.0 ºC |
| Viscosity: | 1.0 cP |
| Dilution factor: | 1 x 10e3 |
| Syringe Pump Speed: | 120 |

### Analysis Settings

| | |
|---|---|
| Detect Threshold: | 5 |
| Blur Size: | Auto |
| Max Jump Distance: | Auto: 14.6 pix |

### Results

Stats: Merged Data

| | |
|---|---|
| Mean: | 89.4 nm |
| Mode: | 85.1 nm |
| SD: | 14.8 nm |
| D10: | 73.4 nm |
| D50: | 86.7 nm |
| D90: | 109.8 nm |

Stats: Mean +/- Standard Error

| | |
|---|---|
| Mean: | 88.8 +/- 2.9 nm |
| Mode: | 84.1 +/- 4.9 nm |
| SD: | 13.8 +/- 1.3 nm |
| D10: | 75.5 +/- 2.7 nm |
| D50: | 87.0 +/- 3.7 nm |
| D90: | 107.4 +/- 4.7 nm |
| Concentration (Upgrade): | 3.14e+09 +/- 5.98e+08 particles/ml |
| | 0.3 +/- 0.0 particles/frame |
| | 0.4 +/- 0.0 centres/frame |

Concentration measurements may be unreliable
See summary file for more info

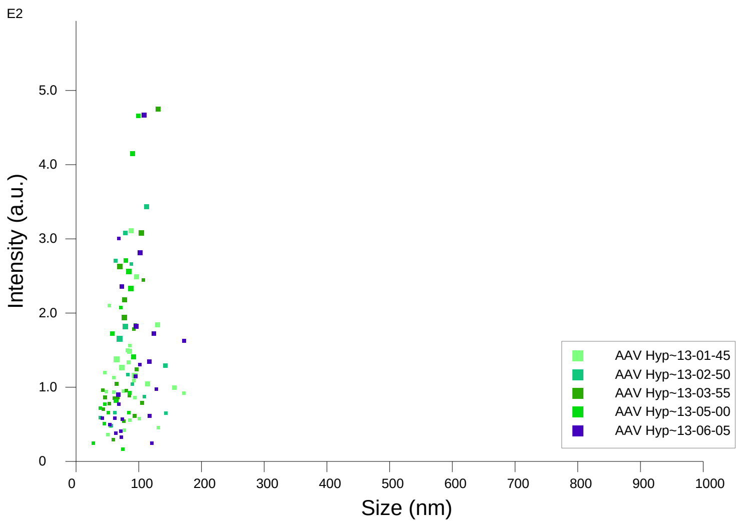


Intensity / Size graph for Experiment:
AAV Hypertransposase 1000x 5 60 120 2024-12-18 12-58-48

**Script Used: (Full Text):**

SOP Standard Measurement 12-58-31PM 18Dec2024.txt
