## Additional file 9 for "Adeno-Associated Virus Co-Precipitation with Extracellular Vesicles for Genome Editing in Rodent Embryo": AAV-Sox2-mStrawberry.pdf

NANOSIGHT

### AAV Sox2 mSTR 1000x 5 60 120 2024-12-18 13-21-30

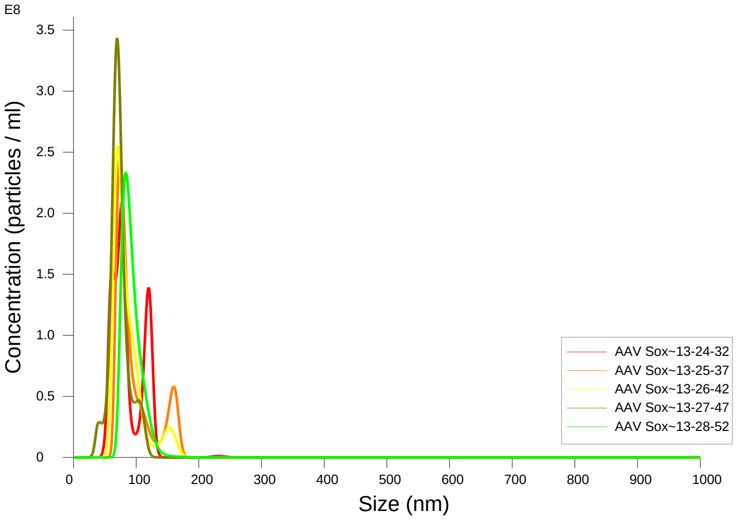


FTLA Concentration / Size graph for Experiment:
AAV Sox2 mSTR 1000x 5 60 120 2024-12-18 13-21-30

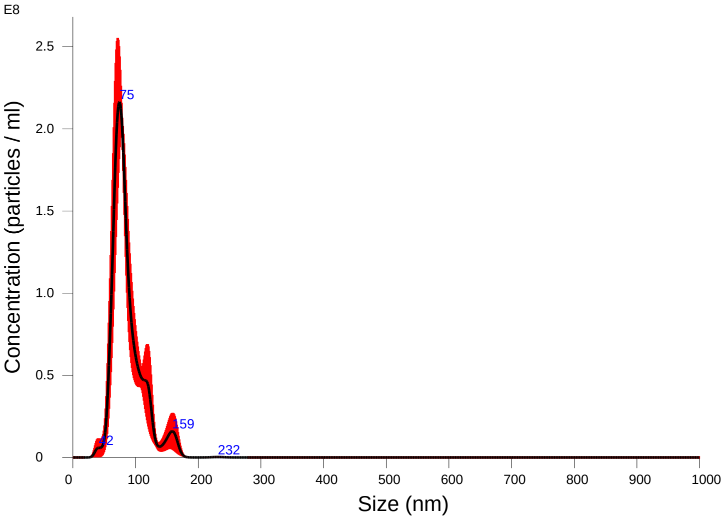


Averaged FTLA Concentration / Size for Experiment:
AAV Sox2 mSTR 1000x 5 60 120 2024-12-18 13-21-30
Error bars indicate + / -1 standard error of the mean

#### Included Files

AAV Sox2 mSTR 1000x 5 60 120 2024-12-18 13-24-32
AAV Sox2 mSTR 1000x 5 60 120 2024-12-18 13-25-37
AAV Sox2 mSTR 1000x 5 60 120 2024-12-18 13-26-42
AAV Sox2 mSTR 1000x 5 60 120 2024-12-18 13-27-47
AAV Sox2 mSTR 1000x 5 60 120 2024-12-18 13-28-52

#### Details

| | |
|---|---|
| NTA Version: | NTA 3.4 - Sample Assistant Build 3.4.4 - SA |
| Script Used: | SOP Standard Measurement 01-21-10PM 18~ |
| Time Captured: | 13:21:30 18/12/2024 |
| Operator: | Jitka |
| Pre-treatment: | |
| Sample Name: | |
| Diluent: | pbs |
| Remarks: | |

#### Capture Settings

| | |
|---|---|
| Camera Type: | sCMOS |
| Laser Type: | Blue405 |
| Camera Level: | 12 |
| Slider Shutter: | 1200 |
| Slider Gain: | 146 |
| FPS | 25.0 |
| Number of Frames: | 1498 |
| Temperature: | 25.0 - 25.0 ºC |
| Viscosity: | 1.0 cP |
| Dilution factor: | 1 x 10e3 |
| Syringe Pump Speed: | 120 |

#### Analysis Settings

| | |
|---|---|
| Detect Threshold: | 5 |
| Blur Size: | Auto |
| Max Jump Distance: | Auto: 14.6 - 18.7 pix |

#### Results

Stats: Merged Data

| | |
|---|---|
| Mean: | 86.3 nm |
| Mode: | 74.3 nm |
| SD: | 24.3 nm |
| D10: | 63.1 nm |
| D50: | 79.9 nm |
| D90: | 119.1 nm |

Stats: Mean +/- Standard Error

| | |
|---|---|
| Mean: | 86.9 +/- 3.9 nm |
| Mode: | 74.9 +/- 2.5 nm |
| SD: | 22.2 +/- 3.0 nm |
| D10: | 65.3 +/- 3.5 nm |
| D50: | 80.2 +/- 3.0 nm |
| D90: | 120.9 +/- 9.8 nm |
| Concentration (Upgrade): | 7.97e+09 +/- 3.14e+08 particles/ml |
| | 0.7 +/- 0.0 particles/frame |
| | 0.8 +/- 0.0 centres/frame |

Concentration measurements may be unreliable
See summary file for more info

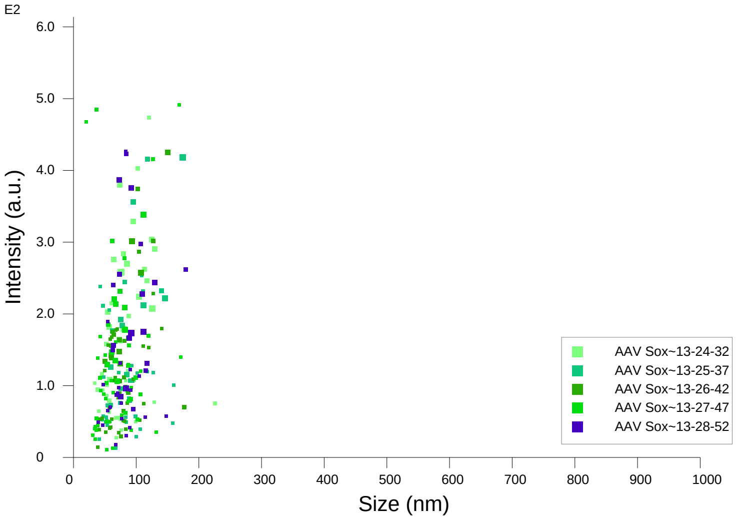


Intensity / Size graph for Experiment:
AAV Sox2 mSTR 1000x 5 60 120 2024-12-18 13-21-30

**Script Used: (Full Text):**

SOP Standard Measurement 01-21-10PM 18Dec2024.txt
