## Additional file 9 for "Adeno-Associated Virus Co-Precipitation with Extracellular Vesicles for Genome Editing in Rodent Embryo": EV-EGFP transposon.pdf

NANOSIGHT

### MPRA EV 1000x 5 60 120 2024-10-30 12-48-03

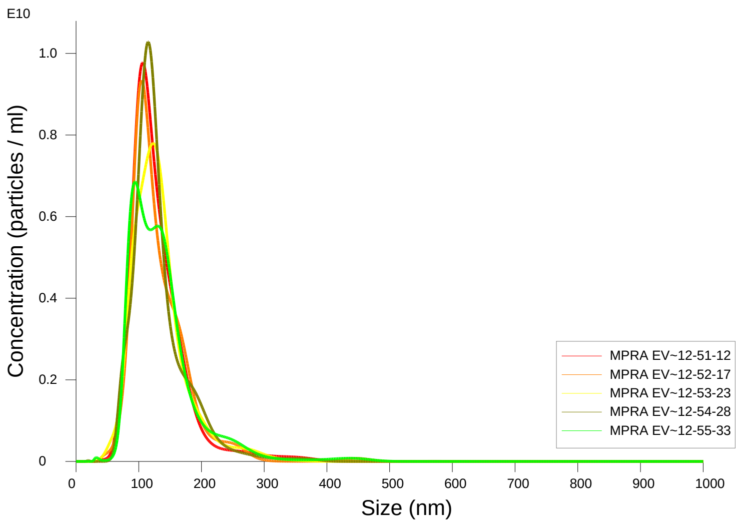


FTLA Concentration / Size graph for Experiment:
MPRA EV 1000x 5 60 120 2024-10-30 12-48-03

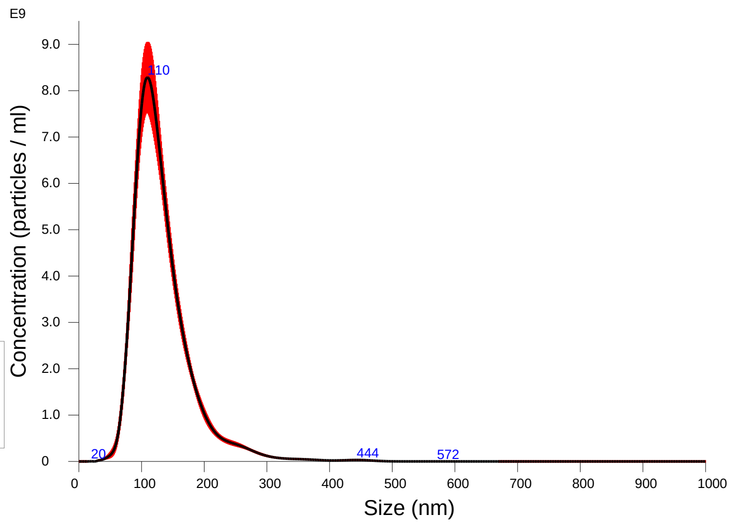


Averaged FTLA Concentration / Size for Experiment:
MPRA EV 1000x 5 60 120 2024-10-30 12-48-03
Error bars indicate + / -1 standard error of the mean

#### Included Files

MPRA EV 1000x 5 60 120 2024-10-30 12-51-12
MPRA EV 1000x 5 60 120 2024-10-30 12-52-17
MPRA EV 1000x 5 60 120 2024-10-30 12-53-23
MPRA EV 1000x 5 60 120 2024-10-30 12-54-28
MPRA EV 1000x 5 60 120 2024-10-30 12-55-33

#### Details

| | |
|---|---|
| NTA Version: | NTA 3.4 - Sample Assistant Build 3.4.4 - SA |
| Script Used: | SOP Standard Measurement 12-48-03PM 30O~ |
| Time Captured: | 12:48:03 30/10/2024 |
| Operator: | Jitka NeburkovÆ |
| Pre-treatment: | |
| Sample Name: | |
| Diluent: | |
| Remarks: | |

#### Capture Settings

| | |
|---|---|
| Camera Type: | sCMOS |
| Laser Type: | Blue405 |
| Camera Level: | 12 |
| Slider Shutter: | 1200 |
| Slider Gain: | 146 |
| FPS | 25.0 |
| Number of Frames: | 1498 |
| Temperature: | 25.0 - 25.0 $^{o}$C |
| Viscosity: | 1.0 cP |
| Dilution factor: | 1 x 10e3 |
| Syringe Pump Speed: | 120 |

#### Analysis Settings

| | |
|---|---|
| Detect Threshold: | 5 |
| Blur Size: | Auto |
| Max Jump Distance: | Auto: 16.7 - 17.2 pix |

#### Results

Stats: Merged Data

| | |
|---|---|
| Mean: | 132.5 nm |
| Mode: | 109.5 nm |
| SD: | 48.3 nm |
| D10: | 87.1 nm |
| D50: | 122.1 nm |
| D90: | 187.2 nm |

Stats: Mean +/- Standard Error

| | |
|---|---|
| Mean: | 132.5 +/- 1.1 nm |
| Mode: | 108.6 +/- 4.9 nm |
| SD: | 48.2 +/- 2.1 nm |
| D10: | 87.2 +/- 0.6 nm |
| D50: | 122.3 +/- 1.3 nm |
| D90: | 187.8 +/- 3.2 nm |
| Concentration (Upgrade): | 6.54e+11 +/- 1.12e+10 particles/ml |
| | 47.0 +/- 0.5 particles/frame |
| | 49.4 +/- 0.4 centres/frame |

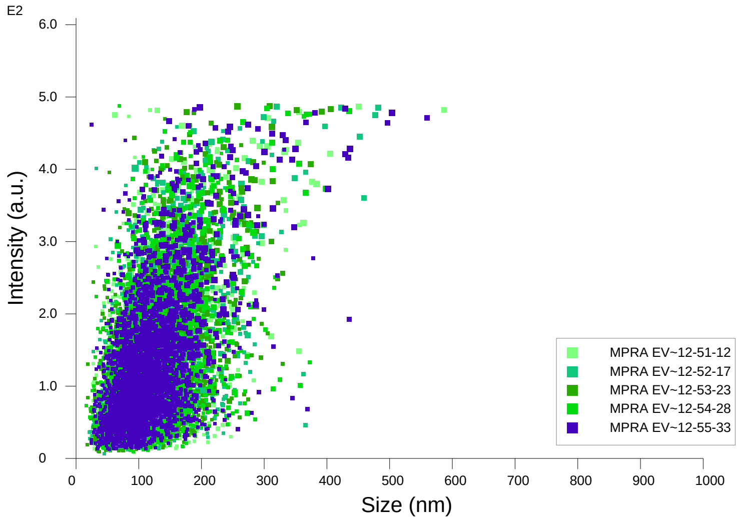


Intensity / Size graph for Experiment:
MPRA EV 1000x 5 60 120 2024-10-30 12-48-03

**Script Used: (Full Text):**

SOP Standard Measurement 12-48-03PM 30Oct2024.txt
