## Additional file 9 for "Adeno-Associated Virus Co-Precipitation with Extracellular Vesicles for Genome Editing in Rodent Embryo": EV-Hypertransposase.pdf

### NANOSIGHT

#### EV Hyp 1000x 5 60 120 2024-10-30 12-32-35

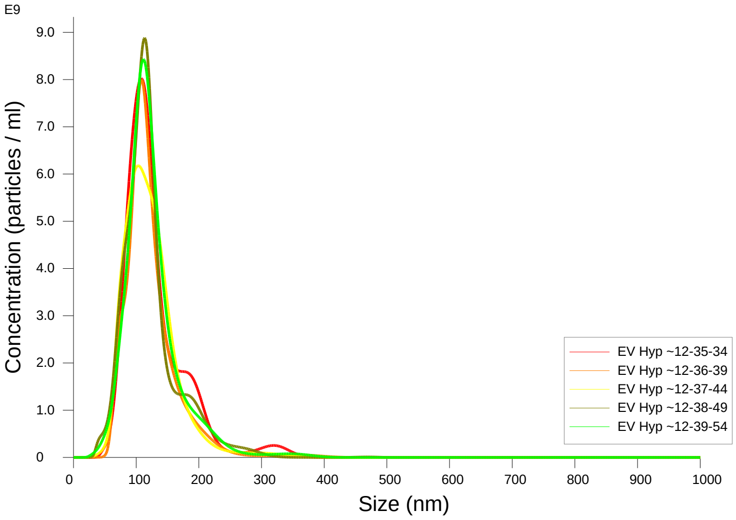


FTLA Concentration / Size graph for Experiment:
EV Hyp 1000x 5 60 120 2024-10-30 12-32-35

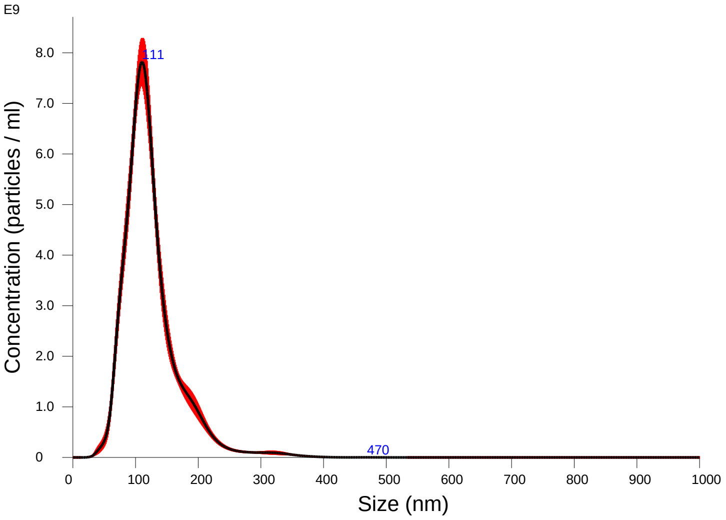


Averaged FTLA Concentration / Size for Experiment:
EV Hyp 1000x 5 60 120 2024-10-30 12-32-35
Error bars indicate + / -1 standard error of the mean

**Included Files**

EV Hyp 1000x 5 60 120 2024-10-30 12-35-34
EV Hyp 1000x 5 60 120 2024-10-30 12-36-39
EV Hyp 1000x 5 60 120 2024-10-30 12-37-44
EV Hyp 1000x 5 60 120 2024-10-30 12-38-49
EV Hyp 1000x 5 60 120 2024-10-30 12-39-54

**Details**

| | |
|---|---|
| NTA Version: | NTA 3.4 - Sample Assistant Build 3.4.4 - SA |
| Script Used: | SOP Standard Measurement 12-32-35PM 30O~ |
| Time Captured: | 12:32:35 30/10/2024 |
| Operator: | Jitka NeburkovÆ |
| Pre-treatment: | |
| Sample Name: | |
| Diluent: | |
| Remarks: | |

**Capture Settings**

| | |
|---|---|
| Camera Type: | sCMOS |
| Laser Type: | Blue405 |
| Camera Level: | 12 |
| Slider Shutter: | 1200 |
| Slider Gain: | 146 |
| FPS | 25.0 |
| Number of Frames: | 1498 |
| Temperature: | 25.0 - 25.0 ºC |
| Viscosity: | 1.0 cP |
| Dilution factor: | 1 x 10e3 |
| Syringe Pump Speed: | 120 |

**Analysis Settings**

| | |
|---|---|
| Detect Threshold: | 5 |
| Blur Size: | Auto |
| Max Jump Distance: | Auto: 17.1 - 20.9 pix |

**Results**

Stats: Merged Data

| | |
|---|---|
| Mean: | 124.1 nm |
| Mode: | 110.4 nm |
| SD: | 45.0 nm |
| D10: | 79.6 nm |
| D50: | 115.2 nm |
| D90: | 180.1 nm |

Stats: Mean +/- Standard Error

| | |
|---|---|
| Mean: | 124.0 +/- 1.3 nm |
| Mode: | 109.0 +/- 1.7 nm |
| SD: | 44.8 +/- 1.3 nm |
| D10: | 79.6 +/- 1.1 nm |
| D50: | 115.2 +/- 0.6 nm |
| D90: | 179.1 +/- 3.8 nm |
| Concentration (Upgrade): | 5.57e+11 +/- 1.69e+10 particles/ml |
| | 41.2 +/- 1.2 particles/frame |
| | 43.6 +/- 0.6 centres/frame |

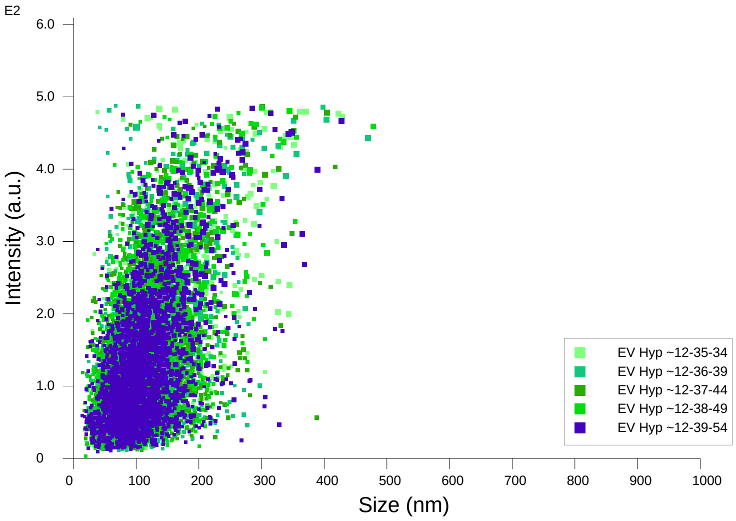


Intensity / Size graph for Experiment:
EV Hyp 1000x 5 60 120 2024-10-30 12-32-35

**Script Used: (Full Text):**

SOP Standard Measurement 12-32-35PM 30Oct2024.txt
