## Additional file 9 for "Adeno-Associated Virus Co-Precipitation with Extracellular Vesicles for Genome Editing in Rodent Embryo": EV-Sox2-mStrawberry.pdf

### NANOSIGHT

#### Sox EV 1000x 5 60 120 2024-10-30 11-50-19

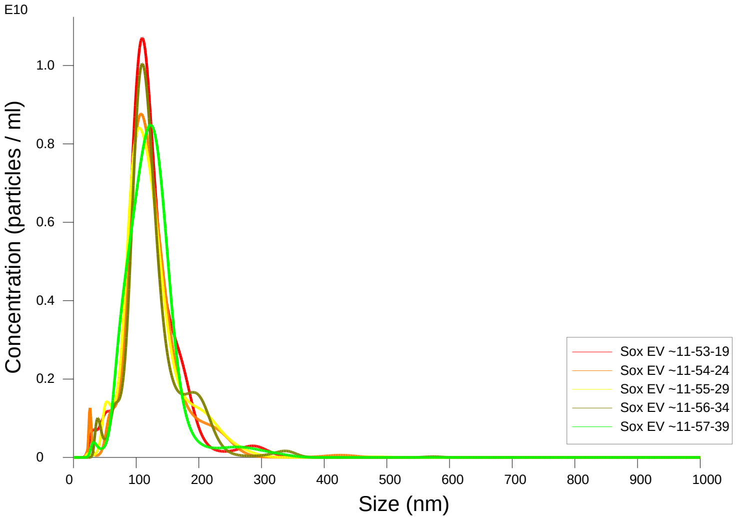


FTLA Concentration / Size graph for Experiment:
Sox EV 1000x 5 60 120 2024-10-30 11-50-19

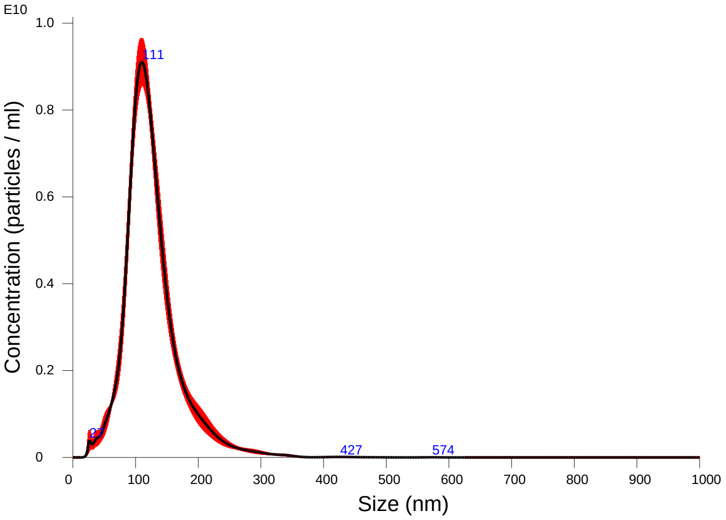


Averaged FTLA Concentration / Size for Experiment:
Sox EV 1000x 5 60 120 2024-10-30 11-50-19
Error bars indicate + / -1 standard error of the mean

##### Included Files

Sox EV 1000x 5 60 120 2024-10-30 11-53-19
Sox EV 1000x 5 60 120 2024-10-30 11-54-24
Sox EV 1000x 5 60 120 2024-10-30 11-55-29
Sox EV 1000x 5 60 120 2024-10-30 11-56-34
Sox EV 1000x 5 60 120 2024-10-30 11-57-39

##### Details

| | |
|---|---|
| NTA Version: | NTA 3.4 - Sample Assistant Build 3.4.4 - SA |
| Script Used: | SOP Standard Measurement 11-50-19AM 30O~ |
| Time Captured: | 11:50:19 30/10/2024 |
| Operator: | Jitka NeburkovÆ |
| Pre-treatment: | |
| Sample Name: | |
| Diluent: | |
| Remarks: | |

##### Capture Settings

| | |
|---|---|
| Camera Type: | sCMOS |
| Laser Type: | Blue405 |
| Camera Level: | 12 |
| Slider Shutter: | 1200 |
| Slider Gain: | 146 |
| FPS | 25.0 |
| Number of Frames: | 1498 |
| Temperature: | 25.0 - 25.0 ºC |
| Viscosity: | 1.0 cP |
| Dilution factor: | 1 x 10e3 |
| Syringe Pump Speed: | 120 |

##### Analysis Settings

| | |
|---|---|
| Detect Threshold: | 5 |
| Blur Size: | Auto |
| Max Jump Distance: | Auto: 20.3 - 22.3 pix |

##### Results

Stats: Merged Data

| | |
|---|---|
| Mean: | 126.0 nm |
| Mode: | 110.0 nm |
| SD: | 45.5 nm |
| D10: | 82.1 nm |
| D50: | 118.2 nm |
| D90: | 180.2 nm |

Stats: Mean +/- Standard Error

| | |
|---|---|
| Mean: | 126.1 +/- 0.8 nm |
| Mode: | 111.1 +/- 3.1 nm |
| SD: | 45.5 +/- 1.0 nm |
| D10: | 82.3 +/- 1.0 nm |
| D50: | 118.2 +/- 0.8 nm |
| D90: | 181.2 +/- 5.1 nm |
| Concentration (Upgrade): | 6.73e+11 +/- 1.60e+10 particles/ml |
| | 50.2 +/- 1.2 particles/frame |
| | 52.1 +/- 0.9 centres/frame |

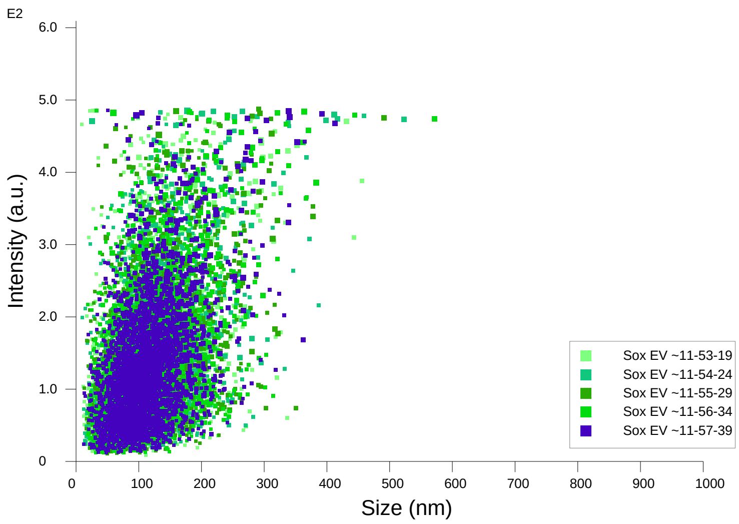


Intensity / Size graph for Experiment:
Sox EV 1000x 5 60 120 2024-10-30 11-50-19

**Script Used: (Full Text):**

SOP Standard Measurement 11-50-19AM 30Oct2024.txt
