## Additional file 9 for "Adeno-Associated Virus Co-Precipitation with Extracellular Vesicles for Genome Editing in Rodent Embryo": Actn1 cKO.pdf

NANOSIGHT

### Actn1 1000x 5 60 120 2024-10-30 11-06-49

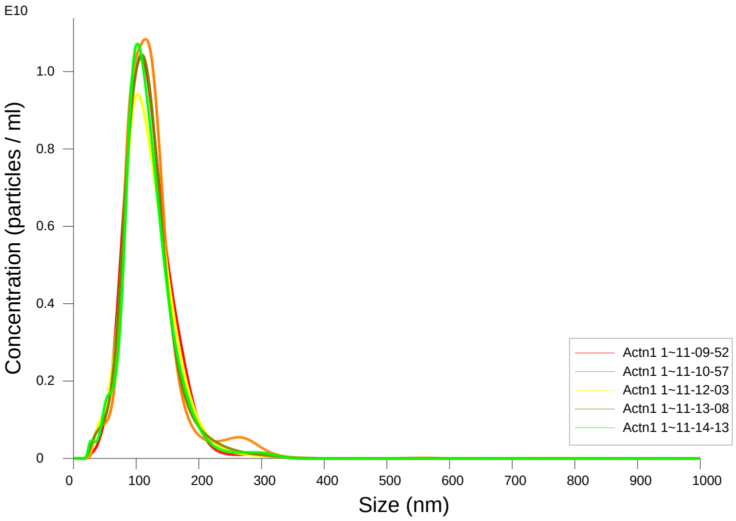


FTLA Concentration / Size graph for Experiment:
Actn1 1000x 5 60 120 2024-10-30 11-06-49

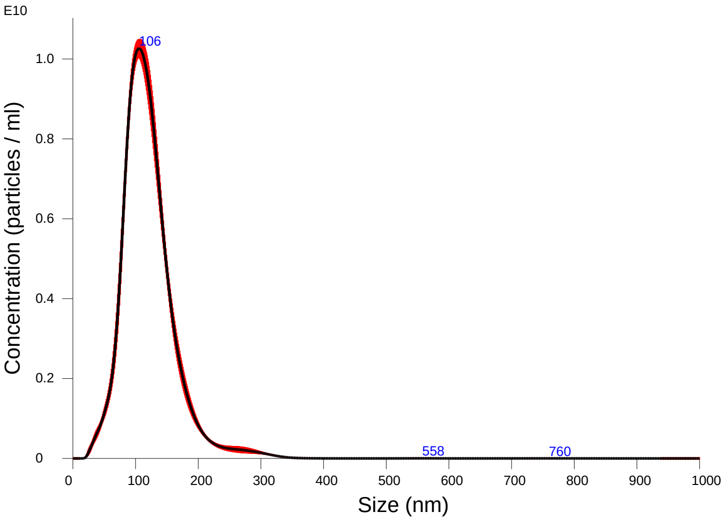


Averaged FTLA Concentration / Size for Experiment:
Actn1 1000x 5 60 120 2024-10-30 11-06-49
Error bars indicate + / -1 standard error of the mean

#### Included Files

Actn1 1000x 5 60 120 2024-10-30 11-09-52
Actn1 1000x 5 60 120 2024-10-30 11-10-57
Actn1 1000x 5 60 120 2024-10-30 11-12-03
Actn1 1000x 5 60 120 2024-10-30 11-13-08
Actn1 1000x 5 60 120 2024-10-30 11-14-13

#### Details

| | |
|---|---|
| NTA Version: | NTA 3.4 - Sample Assistant Build 3.4.4 - SA |
| Script Used: | SOP Standard Measurement 11-06-49AM 30O~ |
| Time Captured: | 11:06:49 30/10/2024 |
| Operator: | Jitka NeburkovÆ |
| Pre-treatment: | |
| Sample Name: | |
| Diluent: | |
| Remarks: | |

#### Capture Settings

| | |
|---|---|
| Camera Type: | sCMOS |
| Laser Type: | Blue405 |
| Camera Level: | 12 |
| Slider Shutter: | 1200 |
| Slider Gain: | 146 |
| FPS | 25.0 |
| Number of Frames: | 1498 |
| Temperature: | 25.0 - 25.0 °C |
| Viscosity: | 1.0 cP |
| Dilution factor: | 1 x 10e3 |
| Syringe Pump Speed: | 120 |

#### Analysis Settings

| | |
|---|---|
| Detect Threshold: | 5 |
| Blur Size: | Auto |
| Max Jump Distance: | Auto: 19.1 - 20.9 pix |

#### Results

Stats: Merged Data

| | |
|---|---|
| Mean: | 120.6 nm |
| Mode: | 105.6 nm |
| SD: | 43.2 nm |
| D10: | 77.1 nm |
| D50: | 114.6 nm |
| D90: | 168.8 nm |

Stats: Mean +/- Standard Error

| | |
|---|---|
| Mean: | 120.5 +/- 0.8 nm |
| Mode: | 107.2 +/- 2.4 nm |
| SD: | 43.0 +/- 1.7 nm |
| D10: | 77.0 +/- 0.9 nm |
| D50: | 114.6 +/- 0.5 nm |
| D90: | 168.7 +/- 0.8 nm |
| Concentration (Upgrade): | 8.36e+11 +/- 1.55e+10 particles/ml |
| | 63.5 +/- 0.9 particles/frame |
| | 65.2 +/- 0.8 centres/frame |

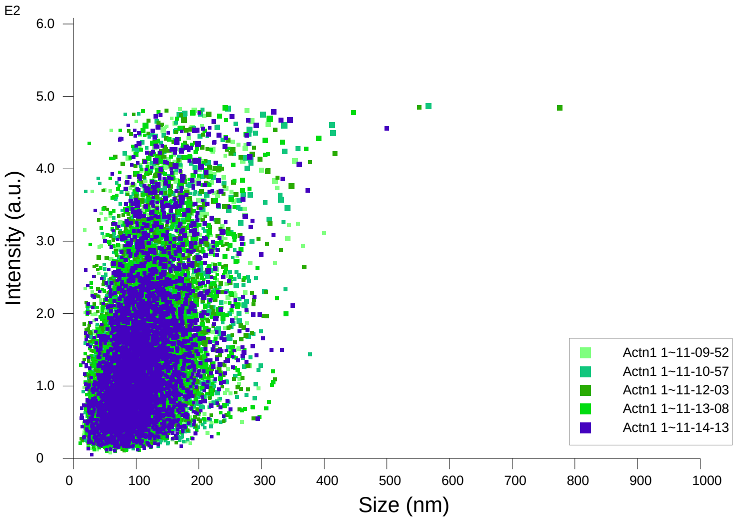


Intensity / Size graph for Experiment:
Actn1 1000x 5 60 120 2024-10-30 11-06-49

**Script Used: (Full Text):**

SOP Standard Measurement 11-06-49AM 30Oct2024.txt
