## Additional file 9 for "Adeno-Associated Virus Co-Precipitation with Extracellular Vesicles for Genome Editing in Rodent Embryo": Alox5ap-CCre-CGFP.pdf

### NANOSIGHT

### alox 1000x 5 60 120 2025-07-24 13-03-39

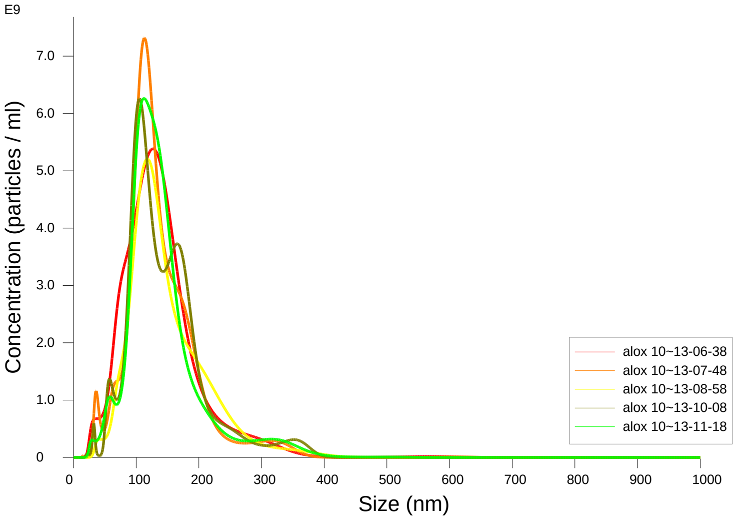


FTLA Concentration / Size graph for Experiment:
alox 1000x 5 60 120 2025-07-24 13-03-39

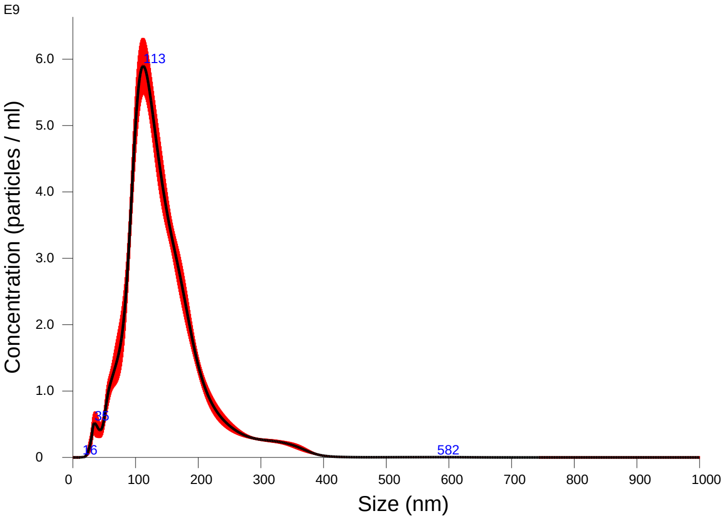


Averaged FTLA Concentration / Size for Experiment:
alox 1000x 5 60 120 2025-07-24 13-03-39
Error bars indicate + / -1 standard error of the mean

#### Included Files

alox 1000x 5 60 120 2025-07-24 13-06-38
alox 1000x 5 60 120 2025-07-24 13-07-48
alox 1000x 5 60 120 2025-07-24 13-08-58
alox 1000x 5 60 120 2025-07-24 13-10-08
alox 1000x 5 60 120 2025-07-24 13-11-18

#### Details

| | |
|---|---|
| NTA Version: | NTA 3.4 - Sample Assistant Build 3.4.4 - SA |
| Script Used: | SOP Standard Measurement 01-03-33PM 24J~ |
| Time Captured: | 13:03:39 24/07/2025 |
| Operator: | jitka |
| Pre-treatment: | |
| Sample Name: | |
| Diluent: | water |
| Remarks: | |

#### Capture Settings

| | |
|---|---|
| Camera Type: | sCMOS |
| Laser Type: | Blue405 |
| Camera Level: | 12 |
| Slider Shutter: | 1200 |
| Slider Gain: | 146 |
| FPS | 25.0 |
| Number of Frames: | 1498 |
| Temperature: | 25.0 - 25.0 °C |
| Viscosity: | 1.0 cP |
| Dilution factor: | 1 x 10e3 |
| Syringe Pump Speed: | 120 |

#### Analysis Settings

| | |
|---|---|
| Detect Threshold: | 5 |
| Blur Size: | Auto |
| Max Jump Distance: | Auto: 17.4 - 22.9 pix |

#### Results

Stats: Merged Data

| | |
|---|---|
| Mean: | 143.3 nm |
| Mode: | 112.3 nm |
| SD: | 61.0 nm |
| D10: | 83.8 nm |
| D50: | 130.6 nm |
| D90: | 216.6 nm |

Stats: Mean +/- Standard Error

| | |
|---|---|
| Mean: | 143.5 +/- 2.1 nm |
| Mode: | 115.1 +/- 3.6 nm |
| SD: | 60.9 +/- 1.3 nm |
| D10: | 84.5 +/- 2.7 nm |
| D50: | 130.9 +/- 1.4 nm |
| D90: | 216.1 +/- 4.6 nm |
| Concentration (Upgrade): | 5.72e+11 +/- 1.52e+10 particles/ml |
| | 43.0 +/- 1.2 particles/frame |
| | 46.9 +/- 1.1 centres/frame |

### NANOSIGHT

### alox 1000x 5 60 120 2025-07-24 13-03-39

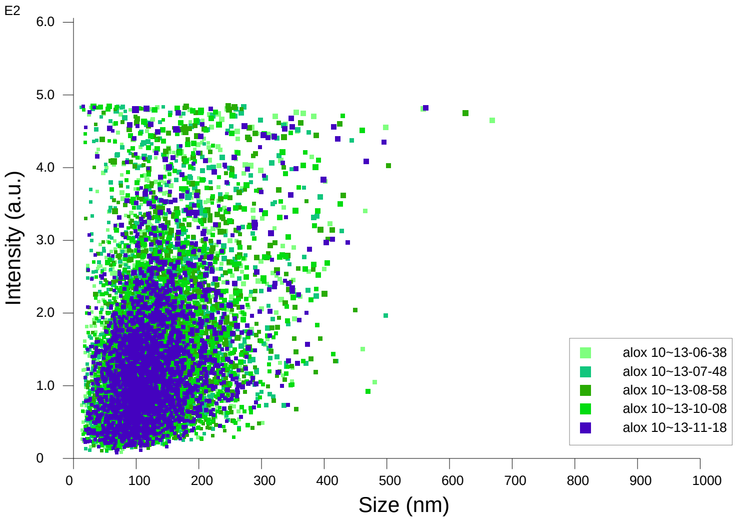


Intensity / Size graph for Experiment:
alox 1000x 5 60 120 2025-07-24 13-03-39

**Script Used: (Full Text):**

SOP Standard Measurement 01-03-33PM 24Jul2025.txt
