## Additional file 9 for "Adeno-Associated Virus Co-Precipitation with Extracellular Vesicles for Genome Editing in Rodent Embryo": Batf3-iCre-eGFP.pdf

### NANOSIGHT

#### EV Batf3 1000x 5 60 120 2024-12-18 09-56-29

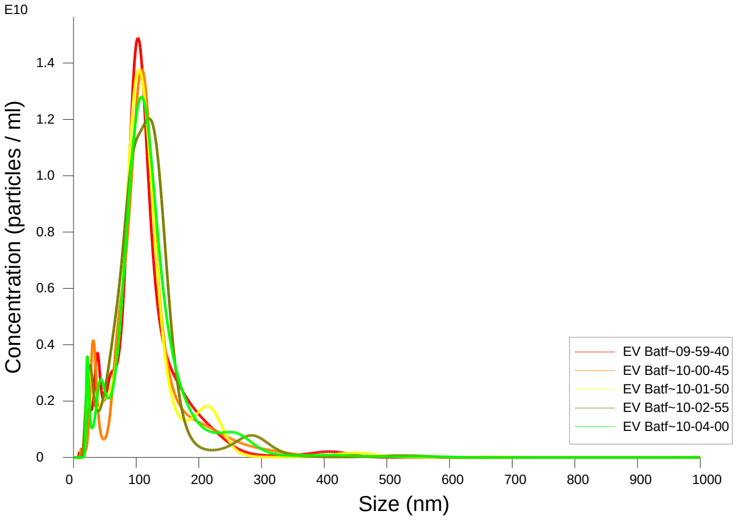


FTLA Concentration / Size graph for Experiment:
EV Batf3 1000x 5 60 120 2024-12-18 09-56-29

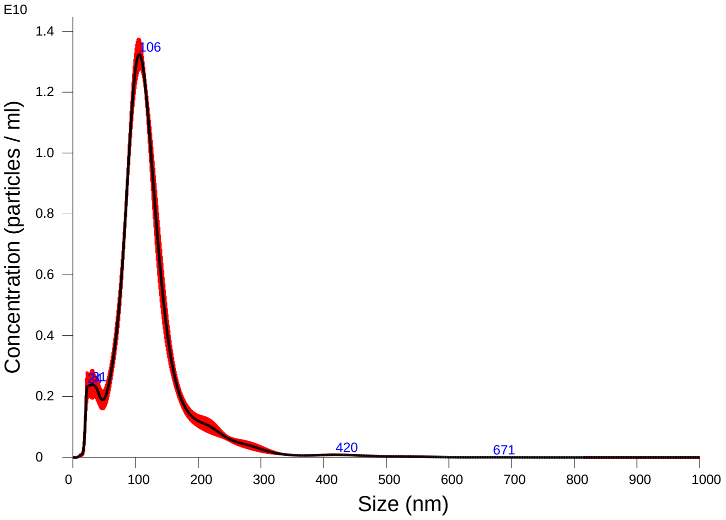


Averaged FTLA Concentration / Size for Experiment:
EV Batf3 1000x 5 60 120 2024-12-18 09-56-29
Error bars indicate + / -1 standard error of the mean

##### Included Files

EV Batf3 1000x 5 60 120 2024-12-18 09-59-40
EV Batf3 1000x 5 60 120 2024-12-18 10-00-45
EV Batf3 1000x 5 60 120 2024-12-18 10-01-50
EV Batf3 1000x 5 60 120 2024-12-18 10-02-55
EV Batf3 1000x 5 60 120 2024-12-18 10-04-00

##### Details

| | |
|---|---|
| NTA Version: | NTA 3.4 - Sample Assistant Build 3.4.4 - SA |
| Script Used: | SOP Standard Measurement 09-56-04AM 18~ |
| Time Captured: | 09:56:29 18/12/2024 |
| Operator: | Jitka |
| Pre-treatment: | |
| Sample Name: | |
| Diluent: | pbs |
| Remarks: | |

##### Capture Settings

| | |
|---|---|
| Camera Type: | sCMOS |
| Laser Type: | Blue405 |
| Camera Level: | 12 |
| Slider Shutter: | 1200 |
| Slider Gain: | 146 |
| FPS | 25.0 |
| Number of Frames: | 1498 |
| Temperature: | 25.0 - 25.0 ºC |
| Viscosity: | 1.0 cP |
| Dilution factor: | 1 x 10e3 |
| Syringe Pump Speed: | 120 |

##### Analysis Settings

| | |
|---|---|
| Detect Threshold: | 5 |
| Blur Size: | Auto |
| Max Jump Distance: | Auto: 23.9 - 26.2 pix |

##### Results

Stats: Merged Data

| | |
|---|---|
| Mean: | 121.9 nm |
| Mode: | 106.0 nm |
| SD: | 63.0 nm |
| D10: | 64.7 nm |
| D50: | 111.2 nm |
| D90: | 187.3 nm |

Stats: Mean +/- Standard Error

| | |
|---|---|
| Mean: | 121.9 +/- 1.1 nm |
| Mode: | 109.0 +/- 3.1 nm |
| SD: | 63.0 +/- 1.0 nm |
| D10: | 64.7 +/- 1.5 nm |
| D50: | 111.2 +/- 1.1 nm |
| D90: | 187.5 +/- 5.7 nm |
| Concentration (Upgrade): | 1.05e+12 +/- 2.43e+10 particles/ml |
| | 84.8 +/- 2.2 particles/frame |
| | 83.8 +/- 1.2 centres/frame |

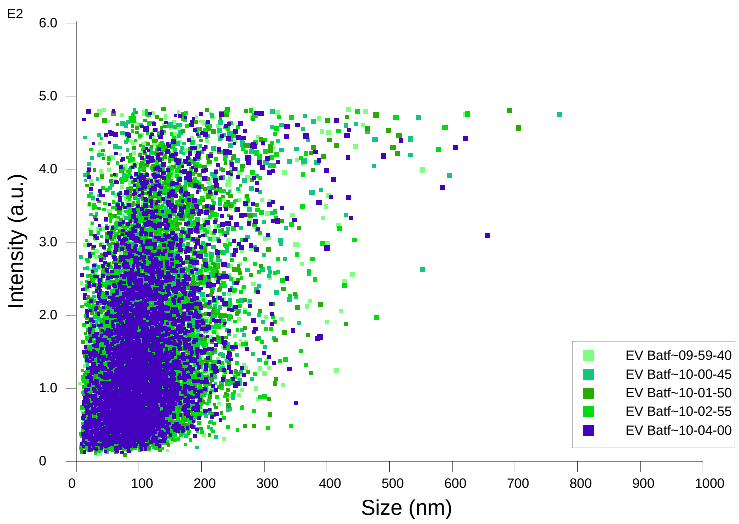


Intensity / Size graph for Experiment:
EV Batf3 1000x 5 60 120 2024-12-18 09-56-29

**Script Used: (Full Text):**

SOP Standard Measurement 09-56-04AM 18Dec2024.txt
