## Additional file 9 for "Adeno-Associated Virus Co-Precipitation with Extracellular Vesicles for Genome Editing in Rodent Embryo": Dpp4-rtTA3-DTR-H1-mKate2.pdf

### NANOSIGHT

#### Dpp4_1000x 5 60 120_2 2024-10-22 12-19-28

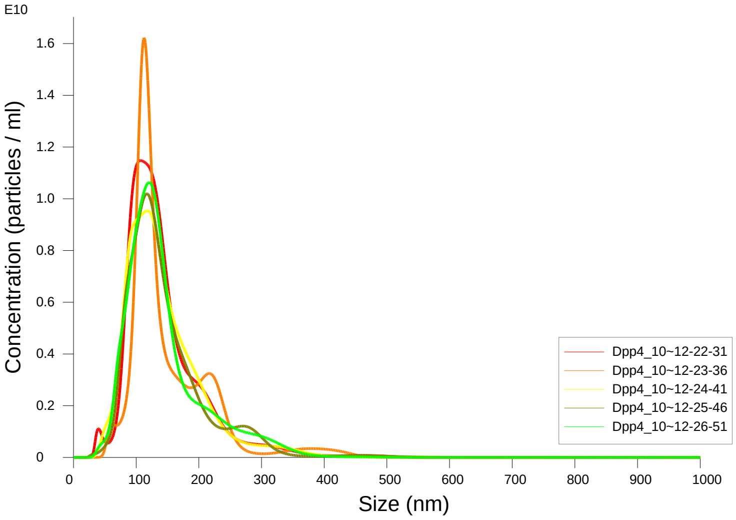


FTLA Concentration / Size graph for Experiment:
Dpp4_1000x 5 60 120_2 2024-10-22 12-19-28

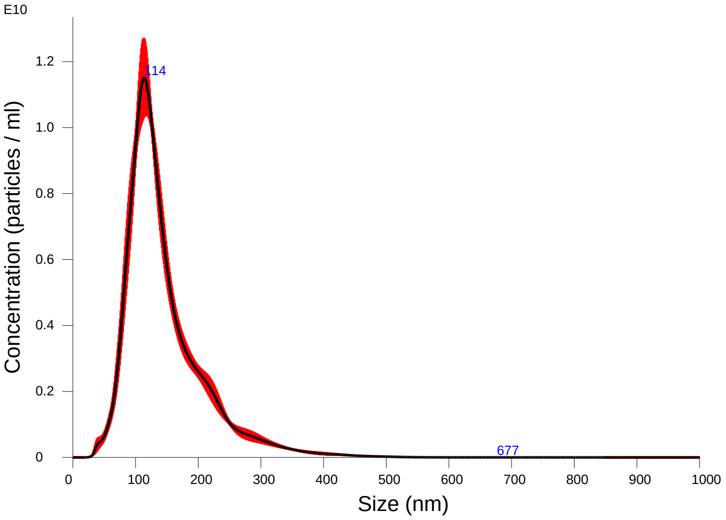


Averaged FTLA Concentration / Size for Experiment:
Dpp4_1000x 5 60 120_2 2024-10-22 12-19-28
Error bars indicate + / -1 standard error of the mean

**Included Files**

Dpp4_1000x 5 60 120_2 2024-10-22 12-22-31
Dpp4_1000x 5 60 120_2 2024-10-22 12-23-36
Dpp4_1000x 5 60 120_2 2024-10-22 12-24-41
Dpp4_1000x 5 60 120_2 2024-10-22 12-25-46
Dpp4_1000x 5 60 120_2 2024-10-22 12-26-51

**Details**

| | |
|---|---|
| NTA Version: | NTA 3.4 - Sample Assistant Build 3.4.4 - SA |
| Script Used: | SOP Standard Measurement 12-19-28PM 22O~ |
| Time Captured: | 12:19:28 22/10/2024 |
| Operator: | Jitka NeburkovÆ |
| Pre-treatment: | |
| Sample Name: | |
| Diluent: | |
| Remarks: | |

**Capture Settings**

| | |
|---|---|
| Camera Type: | sCMOS |
| Laser Type: | Blue405 |
| Camera Level: | 12 |
| Slider Shutter: | 1200 |
| Slider Gain: | 146 |
| FPS | 25.0 |
| Number of Frames: | 1498 |
| Temperature: | 25.0 - 25.0 ºC |
| Viscosity: | 1.0 cP |
| Dilution factor: | 1 x 10e3 |
| Syringe Pump Speed: | 120 |

**Analysis Settings**

| | |
|---|---|
| Detect Threshold: | 5 |
| Blur Size: | Auto |
| Max Jump Distance: | Auto: 16.0 - 18.0 pix |

**Results**

Stats: Merged Data

| | |
|---|---|
| Mean: | 145.6 nm |
| Mode: | 113.8 nm |
| SD: | 64.2 nm |
| D10: | 86.2 nm |
| D50: | 128.3 nm |
| D90: | 227.4 nm |

Stats: Mean +/- Standard Error

| | |
|---|---|
| Mean: | 145.8 +/- 1.6 nm |
| Mode: | 115.1 +/- 2.4 nm |
| SD: | 64.2 +/- 1.2 nm |
| D10: | 86.9 +/- 2.7 nm |
| D50: | 128.3 +/- 0.6 nm |
| D90: | 228.0 +/- 4.2 nm |
| Concentration (Upgrade): | 1.03e+12 +/- 2.46e+10 particles/ml |
| | 75.0 +/- 1.4 particles/frame |
| | 75.8 +/- 0.9 centres/frame |

NANOSIGHT

Dpp4_1000x 5 60 120_2 2024-10-22 12-19-28

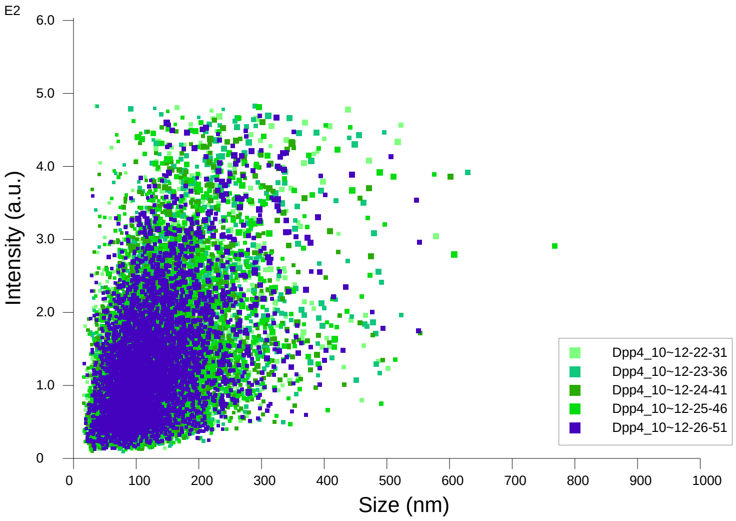


Intensity / Size graph for Experiment:
Dpp4_1000x 5 60 120_2 2024-10-22 12-19-28

**Script Used: (Full Text):**

SOP Standard Measurement 12-19-28PM 22Oct2024.txt
