## Additional file 9 for "Adeno-Associated Virus Co-Precipitation with Extracellular Vesicles for Genome Editing in Rodent Embryo": EGFP transposon.pdf

### NANOSIGHT

#### EV MPRA GFP 1000x 5 60 120 2024-12-18 10-48-30

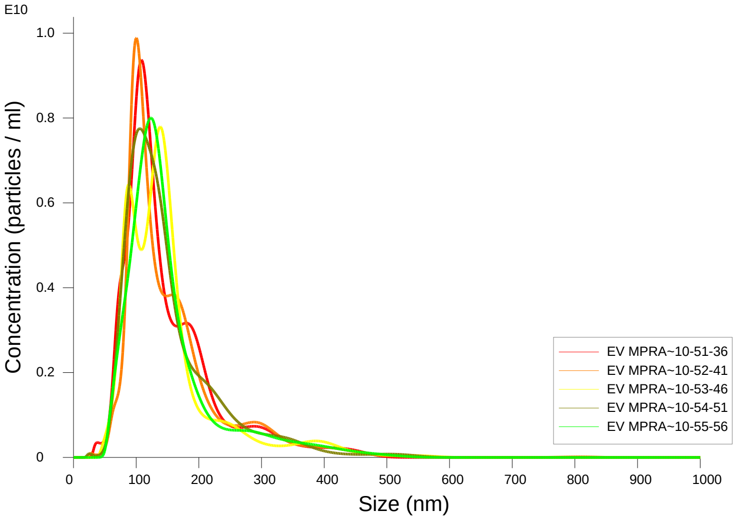


FTLA Concentration / Size graph for Experiment:
EV MPRA GFP 1000x 5 60 120 2024-12-18 10-48-30

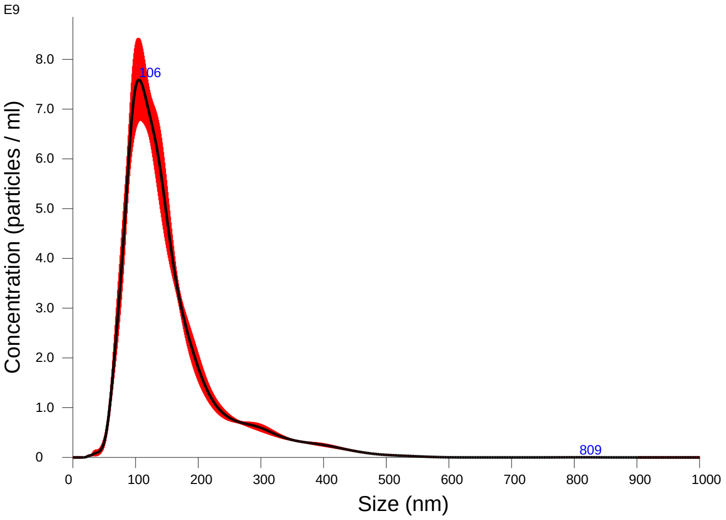


Averaged FTLA Concentration / Size for Experiment:
EV MPRA GFP 1000x 5 60 120 2024-12-18 10-48-30
Error bars indicate + / -1 standard error of the mean

##### Included Files

EV MPRA GFP 1000x 5 60 120 2024-12-18 10-51-36
EV MPRA GFP 1000x 5 60 120 2024-12-18 10-52-41
EV MPRA GFP 1000x 5 60 120 2024-12-18 10-53-46
EV MPRA GFP 1000x 5 60 120 2024-12-18 10-54-51
EV MPRA GFP 1000x 5 60 120 2024-12-18 10-55-56

##### Details

| | |
|---|---|
| NTA Version: | NTA 3.4 - Sample Assistant Build 3.4.4 - SA |
| Script Used: | SOP Standard Measurement 10-48-01AM 18~ |
| Time Captured: | 10:48:30 18/12/2024 |
| Operator: | Jitka |
| Pre-treatment: | |
| Sample Name: | |
| Diluent: | pbs |
| Remarks: | |

##### Capture Settings

| | |
|---|---|
| Camera Type: | sCMOS |
| Laser Type: | Blue405 |
| Camera Level: | 12 |
| Slider Shutter: | 1200 |
| Slider Gain: | 146 |
| FPS | 25.0 |
| Number of Frames: | 1498 |
| Temperature: | 25.0 - 25.0 $^{o}$C |
| Viscosity: | 1.0 cP |
| Dilution factor: | 1 x 10e3 |
| Syringe Pump Speed: | 120 |

##### Analysis Settings

| | |
|---|---|
| Detect Threshold: | 5 |
| Blur Size: | Auto |
| Max Jump Distance: | Auto: 16.4 - 18.3 pix |

##### Results

Stats: Merged Data

| | |
|---|---|
| Mean: | 153.6 nm |
| Mode: | 105.4 nm |
| SD: | 79.2 nm |
| D10: | 84.2 nm |
| D50: | 131.2 nm |
| D90: | 259.5 nm |

Stats: Mean +/- Standard Error

| | |
|---|---|
| Mean: | 153.7 +/- 1.0 nm |
| Mode: | 115.3 +/- 6.9 nm |
| SD: | 79.2 +/- 1.2 nm |
| D10: | 84.2 +/- 1.4 nm |
| D50: | 130.9 +/- 1.4 nm |
| D90: | 259.8 +/- 4.1 nm |
| Concentration (Upgrade): | 8.12e+11 +/- 1.62e+10 particles/ml |
| | 55.8 +/- 0.8 particles/frame |
| | 57.7 +/- 0.6 centres/frame |

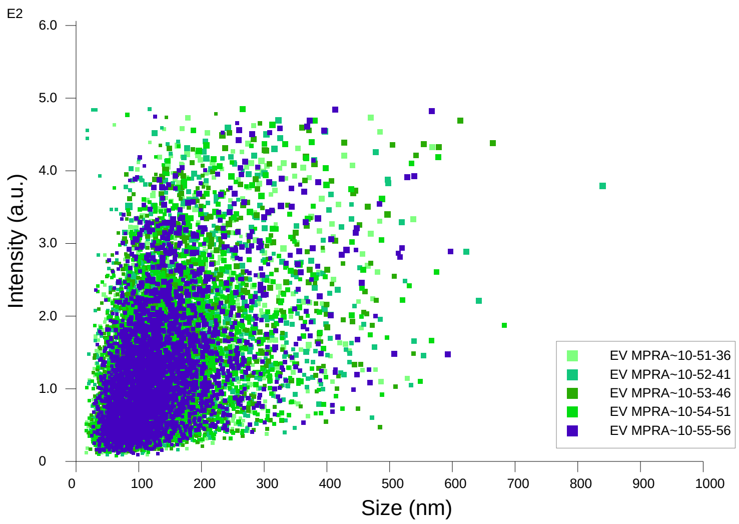


Intensity / Size graph for Experiment:
EV MPRA GFP 1000x 5 60 120 2024-12-18 10-48-30

**Script Used: (Full Text):**

SOP Standard Measurement 10-48-01AM 18Dec2024.txt
