## Additional file 9 for "Adeno-Associated Virus Co-Precipitation with Extracellular Vesicles for Genome Editing in Rodent Embryo": Hypertransposase.pdf

NANOSIGHT

### Hyper 1 1000x 5 60 120 2024-10-30 12-05-24

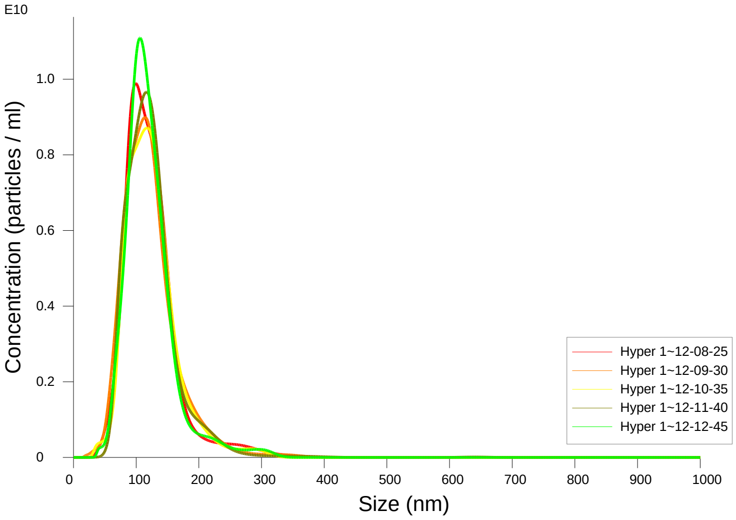


FTLA Concentration / Size graph for Experiment:
Hyper 1 1000x 5 60 120 2024-10-30 12-05-24

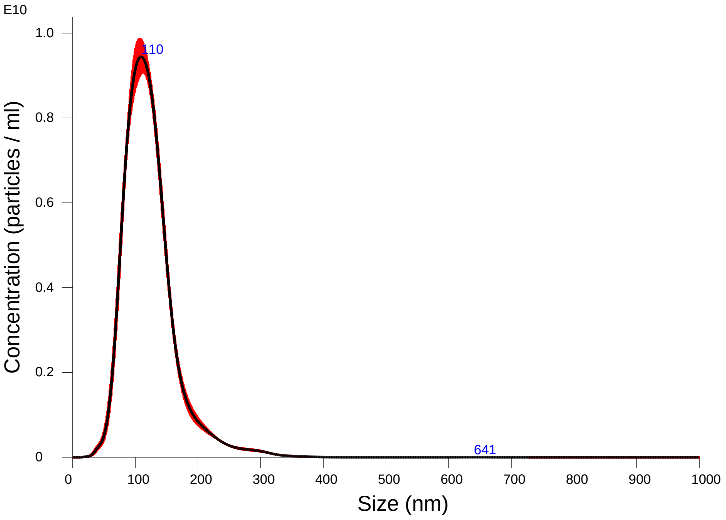


Averaged FTLA Concentration / Size for Experiment:
Hyper 1 1000x 5 60 120 2024-10-30 12-05-24
Error bars indicate + / -1 standard error of the mean

#### Included Files

Hyper 1 1000x 5 60 120 2024-10-30 12-08-25
Hyper 1 1000x 5 60 120 2024-10-30 12-09-30
Hyper 1 1000x 5 60 120 2024-10-30 12-10-35
Hyper 1 1000x 5 60 120 2024-10-30 12-11-40
Hyper 1 1000x 5 60 120 2024-10-30 12-12-45

#### Details

| | |
|---|---|
| NTA Version: | NTA 3.4 - Sample Assistant Build 3.4.4 - SA |
| Script Used: | SOP Standard Measurement 12-05-24PM 30O~ |
| Time Captured: | 12:05:24 30/10/2024 |
| Operator: | Jitka NeburkovÆ |
| Pre-treatment: | |
| Sample Name: | |
| Diluent: | |
| Remarks: | |

#### Capture Settings

| | |
|---|---|
| Camera Type: | sCMOS |
| Laser Type: | Blue405 |
| Camera Level: | 12 |
| Slider Shutter: | 1200 |
| Slider Gain: | 146 |
| FPS | 25.0 |
| Number of Frames: | 1498 |
| Temperature: | 25.0 - 25.0 $^{o}$C |
| Viscosity: | 1.0 cP |
| Dilution factor: | 1 x 10e3 |
| Syringe Pump Speed: | 120 |

#### Analysis Settings

| | |
|---|---|
| Detect Threshold: | 5 |
| Blur Size: | Auto |
| Max Jump Distance: | Auto: 16.8 - 19.9 pix |

#### Results

Stats: Merged Data

| | |
|---|---|
| Mean: | 122.7 nm |
| Mode: | 109.3 nm |
| SD: | 42.6 nm |
| D10: | 79.1 nm |
| D50: | 116.2 nm |
| D90: | 170.2 nm |

Stats: Mean +/- Standard Error

| | |
|---|---|
| Mean: | 122.7 +/- 0.5 nm |
| Mode: | 110.6 +/- 3.5 nm |
| SD: | 42.5 +/- 0.7 nm |
| D10: | 79.2 +/- 1.0 nm |
| D50: | 116.3 +/- 0.8 nm |
| D90: | 170.1 +/- 2.1 nm |
| Concentration (Upgrade): | 7.85e+11 +/- 4.90e+09 particles/ml |
| | 58.1 +/- 0.4 particles/frame |
| | 60.4 +/- 0.7 centres/frame |

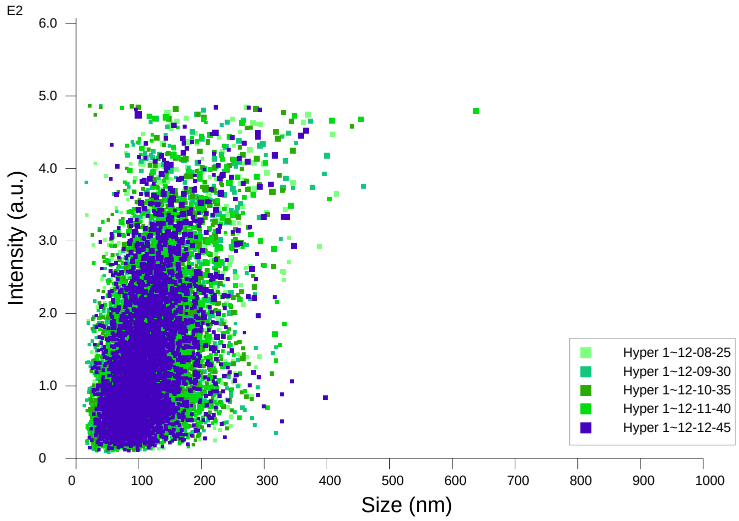


Intensity / Size graph for Experiment:
Hyper 1 1000x 5 60 120 2024-10-30 12-05-24

**Script Used: (Full Text):**

SOP Standard Measurement 12-05-24PM 30Oct2024.txt
