## Additional file 9 for "Adeno-Associated Virus Co-Precipitation with Extracellular Vesicles for Genome Editing in Rodent Embryo": Lck loxP2.pdf

### NANOSIGHT

#### Lck_H1 1000x 5 60 120_2 2024-10-22 11-18-18

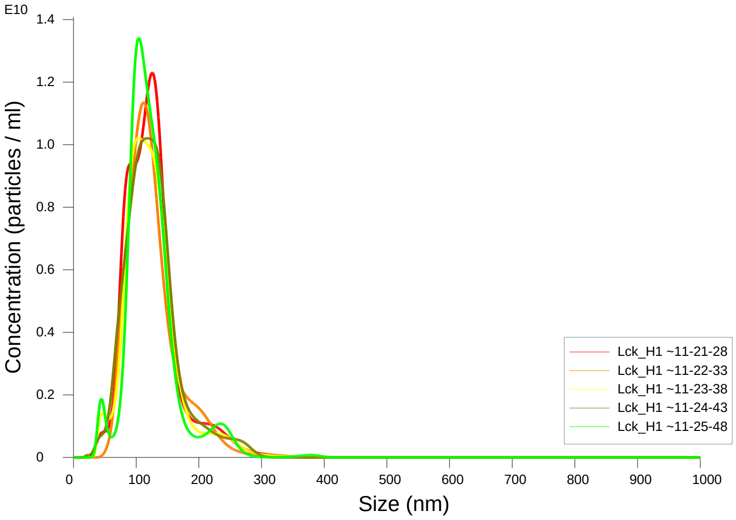


FTLA Concentration / Size graph for Experiment:
Lck_H1 1000x 5 60 120_2 2024-10-22 11-18-18

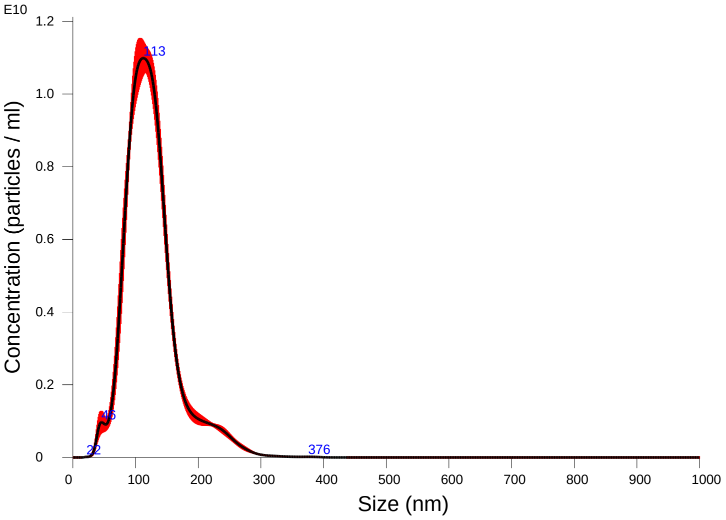


Averaged FTLA Concentration / Size for Experiment:
Lck_H1 1000x 5 60 120_2 2024-10-22 11-18-18
Error bars indicate + / -1 standard error of the mean

**Included Files**

Lck_H1 1000x 5 60 120_2 2024-10-22 11-21-28
Lck_H1 1000x 5 60 120_2 2024-10-22 11-22-33
Lck_H1 1000x 5 60 120_2 2024-10-22 11-23-38
Lck_H1 1000x 5 60 120_2 2024-10-22 11-24-43
Lck_H1 1000x 5 60 120_2 2024-10-22 11-25-48

**Details**

| | |
|---|---|
| NTA Version: | NTA 3.4 - Sample Assistant Build 3.4.4 - SA |
| Script Used: | SOP Standard Measurement 11-18-18AM 22O~ |
| Time Captured: | 11:18:18 22/10/2024 |
| Operator: | Jitka NeburkovÆ |
| Pre-treatment: | |
| Sample Name: | |
| Diluent: | |
| Remarks: | |

**Capture Settings**

| | |
|---|---|
| Camera Type: | sCMOS |
| Laser Type: | Blue405 |
| Camera Level: | 12 |
| Slider Shutter: | 1200 |
| Slider Gain: | 146 |
| FPS | 25.0 |
| Number of Frames: | 1498 |
| Temperature: | 25.0 - 25.0 $^{o}$C |
| Viscosity: | 1.0 cP |
| Dilution factor: | 1 x 10e3 |
| Syringe Pump Speed: | 120 |

**Analysis Settings**

| | |
|---|---|
| Detect Threshold: | 5 |
| Blur Size: | Auto |
| Max Jump Distance: | Auto: 16.8 - 19.3 pix |

**Results**

Stats: Merged Data

| | |
|---|---|
| Mean: | 124.9 nm |
| Mode: | 112.4 nm |
| SD: | 42.6 nm |
| D10: | 81.1 nm |
| D50: | 118.6 nm |
| D90: | 175.5 nm |

Stats: Mean +/- Standard Error

| | |
|---|---|
| Mean: | 124.9 +/- 0.3 nm |
| Mode: | 112.7 +/- 4.1 nm |
| SD: | 42.5 +/- 0.3 nm |
| D10: | 81.3 +/- 1.5 nm |
| D50: | 118.7 +/- 0.6 nm |
| D90: | 175.1 +/- 2.4 nm |
| Concentration (Upgrade): | 9.12e+11 +/- 2.13e+10 particles/ml |
| | 63.2 +/- 1.6 particles/frame |
| | 64.5 +/- 1.2 centres/frame |

### NANOSIGHT



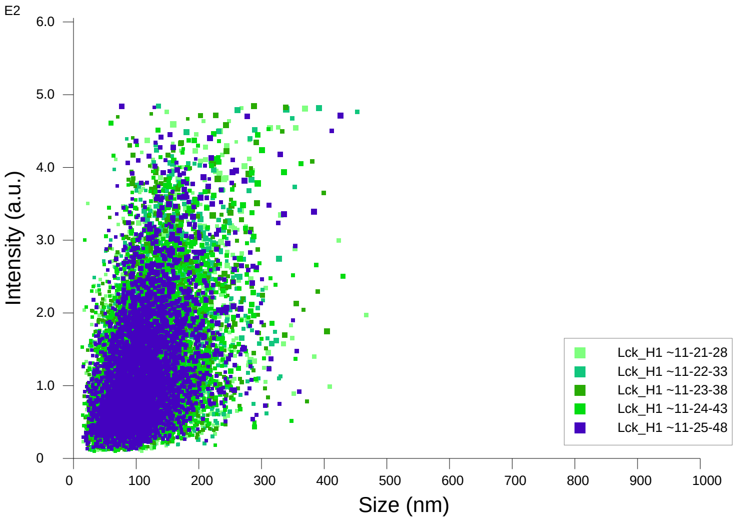


Intensity / Size graph for Experiment:
Lck_H1 1000x 5 60 120_2 2024-10-22 11-18-18

**Script Used: (Full Text):**

SOP Standard Measurement 11-18-18AM 22Oct2024.txt
