## Additional file 9 for "Adeno-Associated Virus Co-Precipitation with Extracellular Vesicles for Genome Editing in Rodent Embryo": Lyve1NCre-NGFP.pdf

### NANOSIGHT Lyve 1000x 5 60 120 2025-07-24 13-22-11

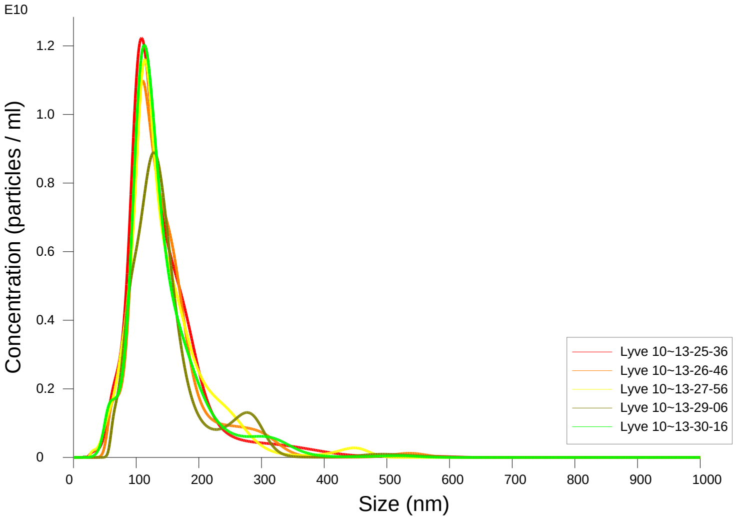


FTLA Concentration / Size graph for Experiment:
Lyve 1000x 5 60 120 2025-07-24 13-22-11

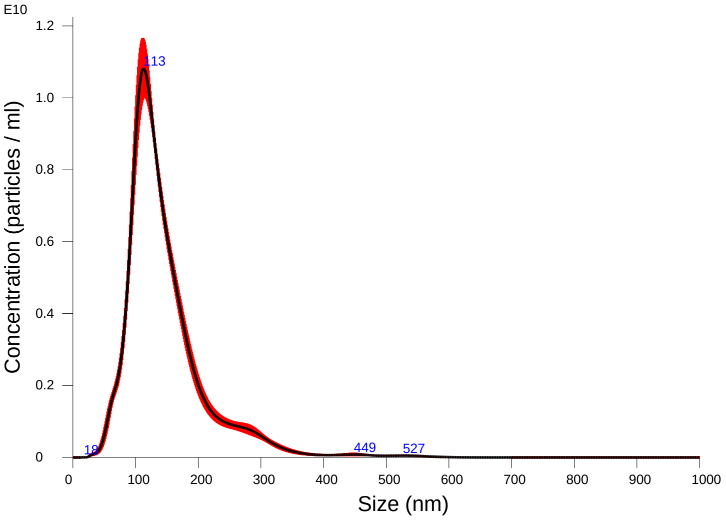


Averaged FTLA Concentration / Size for Experiment:
Lyve 1000x 5 60 120 2025-07-24 13-22-11
Error bars indicate + / -1 standard error of the mean

#### Included Files

Lyve 1000x 5 60 120 2025-07-24 13-25-36
Lyve 1000x 5 60 120 2025-07-24 13-26-46
Lyve 1000x 5 60 120 2025-07-24 13-27-56
Lyve 1000x 5 60 120 2025-07-24 13-29-06
Lyve 1000x 5 60 120 2025-07-24 13-30-16

#### Details

| | |
|---|---|
| NTA Version: | NTA 3.4 - Sample Assistant Build 3.4.4 - SA |
| Script Used: | SOP Standard Measurement 01-21-53PM 24J~ |
| Time Captured: | 13:22:11 24/07/2025 |
| Operator: | jitka |
| Pre-treatment: | |
| Sample Name: | |
| Diluent: | water |
| Remarks: | |

#### Capture Settings

| | |
|---|---|
| Camera Type: | sCMOS |
| Laser Type: | Blue405 |
| Camera Level: | 12 |
| Slider Shutter: | 1200 |
| Slider Gain: | 146 |
| FPS | 25.0 |
| Number of Frames: | 1498 |
| Temperature: | 25.0 $^{o}$C |
| Viscosity: | 1.0 cP |
| Dilution factor: | 1 x 10e3 |
| Syringe Pump Speed: | 120 |

#### Analysis Settings

| | |
|---|---|
| Detect Threshold: | 5 |
| Blur Size: | Auto |
| Max Jump Distance: | Auto: 15.6 - 18.8 pix |

#### Results

Stats: Merged Data

| | |
|---|---|
| Mean: | 147.2 nm |
| Mode: | 112.8 nm |
| SD: | 66.5 nm |
| D10: | 89.0 nm |
| D50: | 130.3 nm |
| D90: | 227.6 nm |

Stats: Mean +/- Standard Error

| | |
|---|---|
| Mean: | 147.4 +/- 1.3 nm |
| Mode: | 114.8 +/- 3.5 nm |
| SD: | 66.5 +/- 0.8 nm |
| D10: | 89.1 +/- 1.0 nm |
| D50: | 130.4 +/- 1.4 nm |
| D90: | 229.1 +/- 6.5 nm |
| Concentration (Upgrade): | 9.56e+11 +/- 3.83e+10 particles/ml |
| | 68.0 +/- 2.4 particles/frame |
| | 70.3 +/- 1.5 centres/frame |

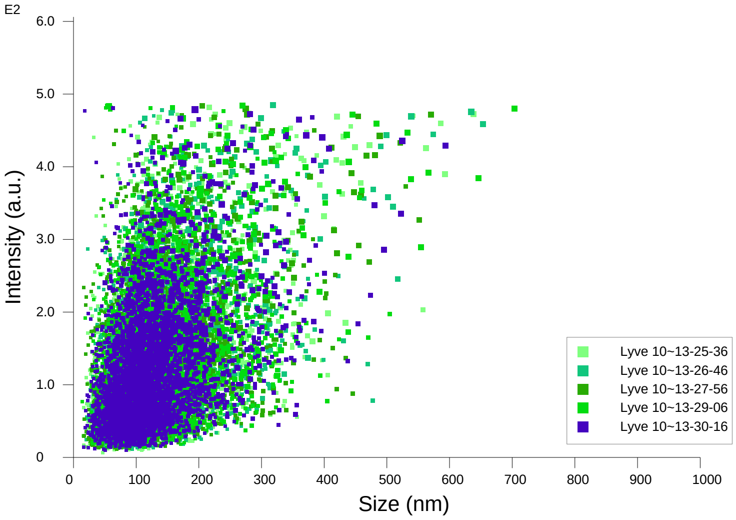


Intensity / Size graph for Experiment:
Lyve 1000x 5 60 120 2025-07-24 13-22-11

**Script Used: (Full Text):**

SOP Standard Measurement 01-21-53PM 24Jul2025.txt
