## Additional file 9 for "Adeno-Associated Virus Co-Precipitation with Extracellular Vesicles for Genome Editing in Rodent Embryo": Muc1_Muc1_FS 5 60 120 500x 2026-03-09 12-41-52-ExperimentReport.pdf

### NANOSIGHT

#### Muc1_Muc1_FS 5 60 120 500x 2026-03-09 12-41-52

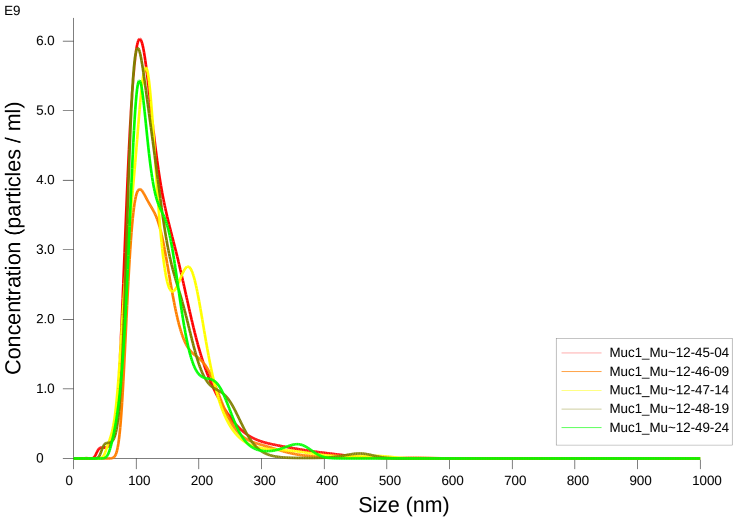


FTLA Concentration / Size graph for Experiment:
Muc1_Muc1_FS 5 60 120 500x 2026-03-09 12-41-52

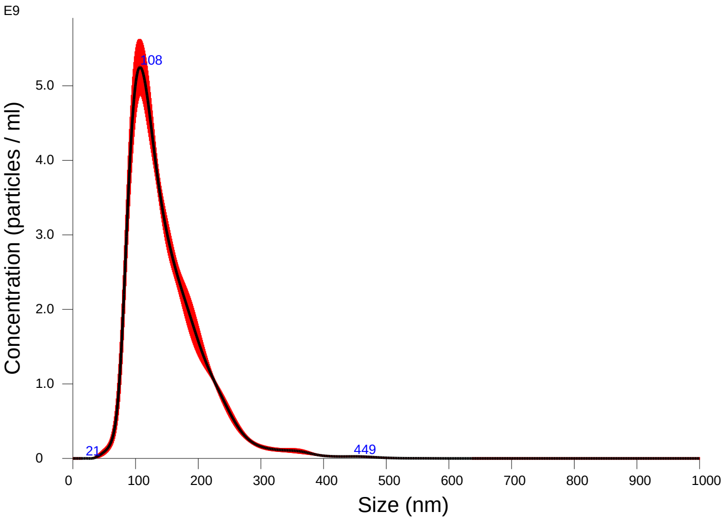


Averaged FTLA Concentration / Size for Experiment:
Muc1_Muc1_FS 5 60 120 500x 2026-03-09 12-41-52
Error bars indicate + / -1 standard error of the mean

##### Included Files

Muc1_Muc1_FS 5 60 120 500x 2026-03-09 12-45-04
Muc1_Muc1_FS 5 60 120 500x 2026-03-09 12-46-09
Muc1_Muc1_FS 5 60 120 500x 2026-03-09 12-47-14
Muc1_Muc1_FS 5 60 120 500x 2026-03-09 12-48-19
Muc1_Muc1_FS 5 60 120 500x 2026-03-09 12-49-24

##### Details

| | |
|---|---|
| NTA Version: | NTA 3.4 - Sample Assistant Build 3.4.4 - SA |
| Script Used: | SOP Standard Measurement 12-41-49PM 09~ |
| Time Captured: | 12:41:52 09/03/2026 |
| Operator: | jitka |
| Pre-treatment: | 500x |
| Sample Name: | FS |
| Diluent: | pbs |
| Remarks: | |

##### Capture Settings

| | |
|---|---|
| Camera Type: | sCMOS |
| Laser Type: | Blue405 |
| Camera Level: | 12 |
| Slider Shutter: | 1200 |
| Slider Gain: | 146 |
| FPS | 25.0 |
| Number of Frames: | 1498 |
| Temperature: | 25.0 - 25.0 $^{o}$C |
| Viscosity: | 1.0 cP |
| Dilution factor: | 5 x 10e2 |
| Syringe Pump Speed: | 120 |

##### Analysis Settings

| | |
|---|---|
| Detect Threshold: | 5 |
| Blur Size: | Auto |
| Max Jump Distance: | Auto: 14.8 - 16.6 pix |

##### Results

Stats: Merged Data

| | |
|---|---|
| Mean: | 148.6 nm |
| Mode: | 107.2 nm |
| SD: | 59.3 nm |
| D10: | 91.1 nm |
| D50: | 133.4 nm |
| D90: | 224.4 nm |

Stats: Mean +/- Standard Error

| | |
|---|---|
| Mean: | 148.8 +/- 1.4 nm |
| Mode: | 106.9 +/- 2.2 nm |
| SD: | 59.1 +/- 0.8 nm |
| D10: | 91.2 +/- 0.9 nm |
| D50: | 133.7 +/- 1.6 nm |
| D90: | 224.5 +/- 1.9 nm |
| Concentration (Upgrade): | 5.03e+11 +/- 2.75e+10 particles/ml |
| | 79.1 +/- 4.5 particles/frame |
| | 79.5 +/- 3.6 centres/frame |

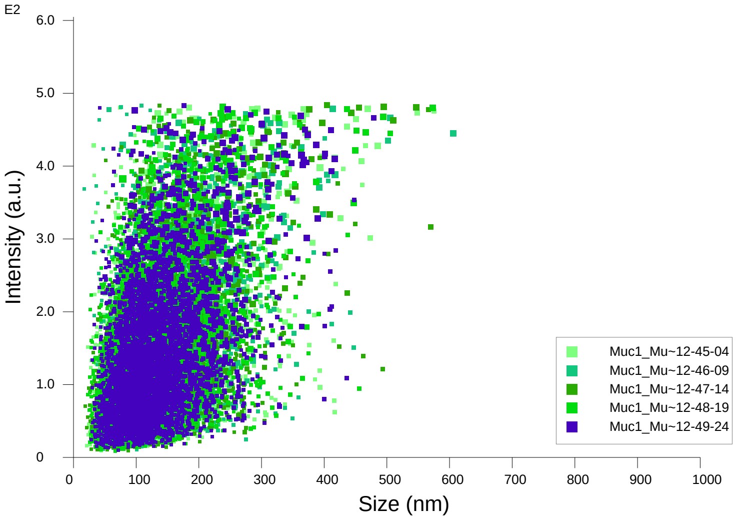


Intensity / Size graph for Experiment:
Muc1_Muc1_FS 5 60 120 500x 2026-03-09 12-41-52

**Script Used: (Full Text):**

SOP Standard Measurement 12-41-49PM 09Mar2026.txt
