## Additional file 9 for "Adeno-Associated Virus Co-Precipitation with Extracellular Vesicles for Genome Editing in Rodent Embryo": Muc1_Muc1_WT 5 60 120 1000x 2026-03-09 12-56-12-ExperimentReport.pdf

### NANOSIGHT

#### Muc1_Muc1_WT 5 60 120 1000x 2026-03-09 12-56-12

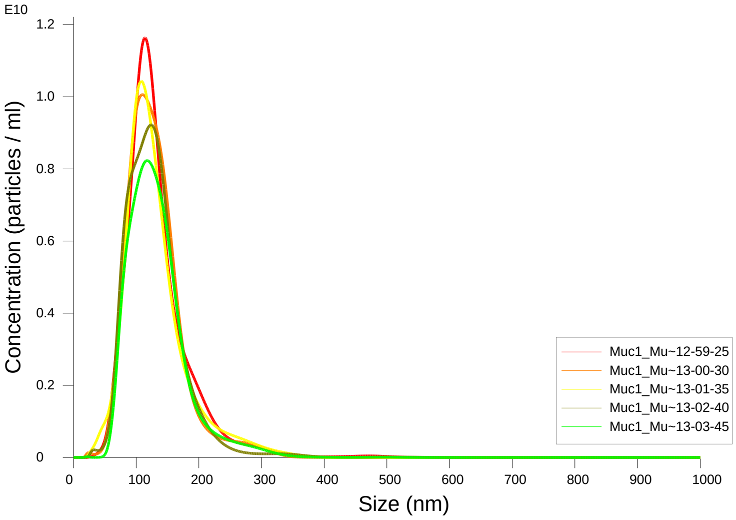


FTLA Concentration / Size graph for Experiment:
Muc1_Muc1_WT 5 60 120 1000x 2026-03-09 12-56-12

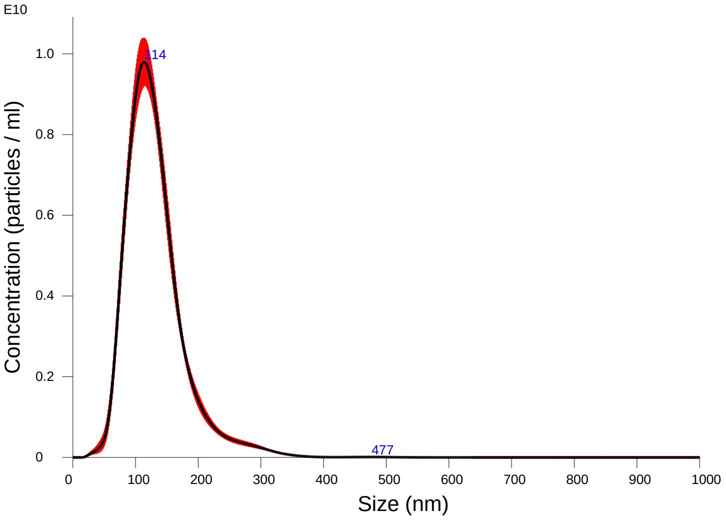


Averaged FTLA Concentration / Size for Experiment:
Muc1_Muc1_WT 5 60 120 1000x 2026-03-09 12-56-12
Error bars indicate + / -1 standard error of the mean

**Included Files**

Muc1_Muc1_WT 5 60 120 1000x 2026-03-09 12-59-25
Muc1_Muc1_WT 5 60 120 1000x 2026-03-09 13-00-30
Muc1_Muc1_WT 5 60 120 1000x 2026-03-09 13-01-35
Muc1_Muc1_WT 5 60 120 1000x 2026-03-09 13-02-40
Muc1_Muc1_WT 5 60 120 1000x 2026-03-09 13-03-45

**Details**

| | |
|---|---|
| NTA Version: | NTA 3.4 - Sample Assistant Build 3.4.4 - SA |
| Script Used: | SOP Standard Measurement 12-56-10PM 09~ |
| Time Captured: | 12:56:12 09/03/2026 |
| Operator: | jitka |
| Pre-treatment: | 1000x |
| Sample Name: | WT |
| Diluent: | pbs |
| Remarks: | |

**Capture Settings**

| | |
|---|---|
| Camera Type: | sCMOS |
| Laser Type: | Blue405 |
| Camera Level: | 12 |
| Slider Shutter: | 1200 |
| Slider Gain: | 146 |
| FPS | 25.0 |
| Number of Frames: | 1498 |
| Temperature: | 25.0 - 25.0 ºC |
| Viscosity: | 1.0 cP |
| Dilution factor: | 1 x 10e3 |
| Syringe Pump Speed: | 120 |

**Analysis Settings**

| | |
|---|---|
| Detect Threshold: | 5 |
| Blur Size: | Auto |
| Max Jump Distance: | Auto: 16.3 - 19.9 pix |

**Results**

Stats: Merged Data

| | |
|---|---|
| Mean: | 131.0 nm |
| Mode: | 113.5 nm |
| SD: | 48.3 nm |
| D10: | 81.7 nm |
| D50: | 122.8 nm |
| D90: | 187.3 nm |

Stats: Mean +/- Standard Error

| | |
|---|---|
| Mean: | 131.0 +/- 0.9 nm |
| Mode: | 114.7 +/- 2.8 nm |
| SD: | 48.1 +/- 1.4 nm |
| D10: | 81.8 +/- 0.7 nm |
| D50: | 122.9 +/- 0.9 nm |
| D90: | 187.6 +/- 3.3 nm |
| Concentration (Upgrade): | 8.88e+11 +/- 2.59e+10 particles/ml |
| | 65.6 +/- 1.8 particles/frame |
| | 66.0 +/- 1.0 centres/frame |

### NANOSIGHT

#### Muc1_Muc1_WT 5 60 120 1000x 2026-03-09 12-56-12

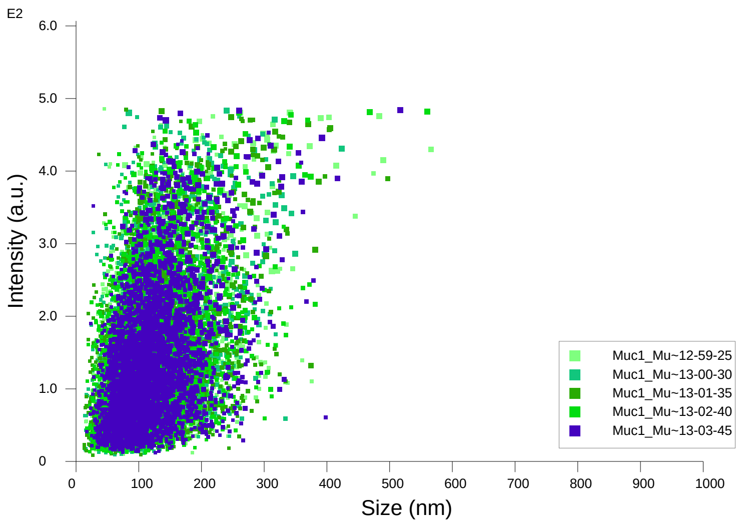


Intensity / Size graph for Experiment:
Muc1_Muc1_WT 5 60 120 1000x 2026-03-09 12-56-12

**Script Used: (Full Text):**

SOP Standard Measurement 12-56-10PM 09Mar2026.txt
