## Additional file 9 for "Adeno-Associated Virus Co-Precipitation with Extracellular Vesicles for Genome Editing in Rodent Embryo": Nes-rtTA3-DTR-IRES-iRFP670.pdf

NANOSIGHT

### Nes_1000x 5 60120 2024-10-22 11-40-17

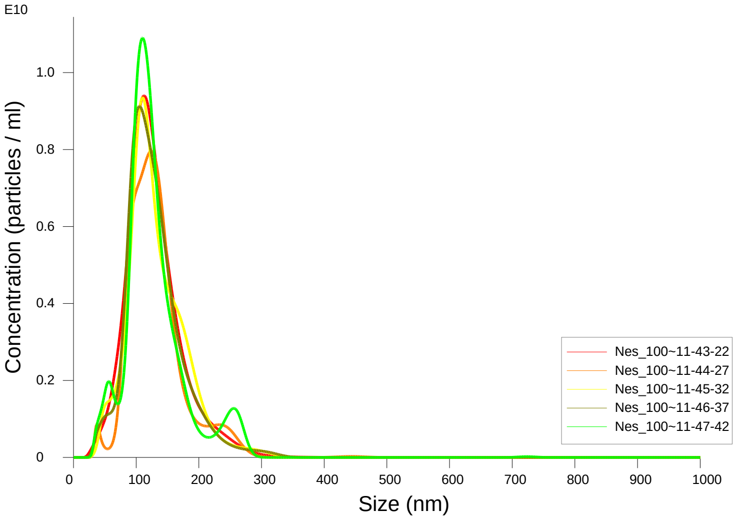


FTLA Concentration / Size graph for Experiment:
Nes_1000x 5 60120 2024-10-22 11-40-17

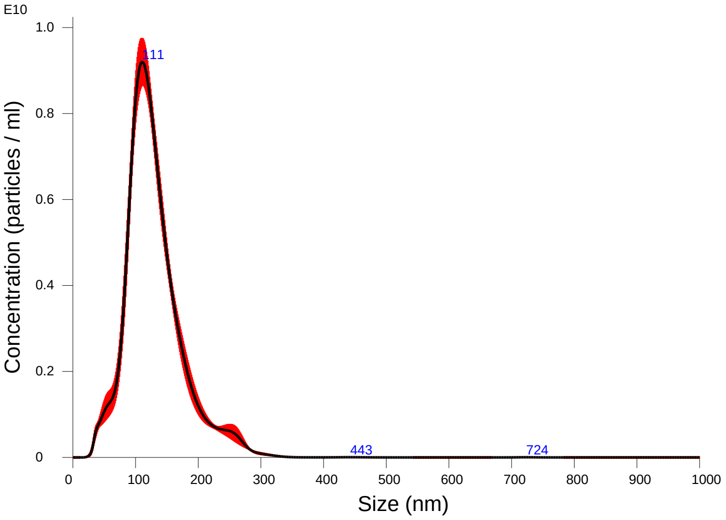


Averaged FTLA Concentration / Size for Experiment:
Nes_1000x 5 60120 2024-10-22 11-40-17
Error bars indicate + / -1 standard error of the mean

#### Included Files

Nes_1000x 5 60120 2024-10-22 11-43-22
Nes_1000x 5 60120 2024-10-22 11-44-27
Nes_1000x 5 60120 2024-10-22 11-45-32
Nes_1000x 5 60120 2024-10-22 11-46-37
Nes_1000x 5 60120 2024-10-22 11-47-42

#### Details

| | |
|---|---|
| NTA Version: | NTA 3.4 - Sample Assistant Build 3.4.4 - SA |
| Script Used: | SOP Standard Measurement 11-40-17AM 22O~ |
| Time Captured: | 11:40:17 22/10/2024 |
| Operator: | Jitka NeburkovÆ |
| Pre-treatment: | |
| Sample Name: | |
| Diluent: | |
| Remarks: | |

#### Capture Settings

| | |
|---|---|
| Camera Type: | sCMOS |
| Laser Type: | Blue405 |
| Camera Level: | 12 |
| Slider Shutter: | 1200 |
| Slider Gain: | 146 |
| FPS | 25.0 |
| Number of Frames: | 1498 |
| Temperature: | 25.0 - 25.0 $^{o}$C |
| Viscosity: | 1.0 cP |
| Dilution factor: | 1 x 10e3 |
| Syringe Pump Speed: | 120 |

#### Analysis Settings

| | |
|---|---|
| Detect Threshold: | 5 |
| Blur Size: | Auto |
| Max Jump Distance: | Auto: 18.9 - 20.4 pix |

#### Results

Stats: Merged Data

| | |
|---|---|
| Mean: | 129.7 nm |
| Mode: | 110.7 nm |
| SD: | 46.4 nm |
| D10: | 82.8 nm |
| D50: | 121.7 nm |
| D90: | 187.8 nm |

Stats: Mean +/- Standard Error

| | |
|---|---|
| Mean: | 129.7 +/- 0.8 nm |
| Mode: | 112.5 +/- 3.1 nm |
| SD: | 46.3 +/- 1.2 nm |
| D10: | 82.7 +/- 1.0 nm |
| D50: | 121.8 +/- 0.9 nm |
| D90: | 187.8 +/- 1.2 nm |
| Concentration (Upgrade): | 7.52e+11 +/- 1.97e+10 particles/ml |
| | 54.6 +/- 1.5 particles/frame |
| | 56.4 +/- 1.2 centres/frame |

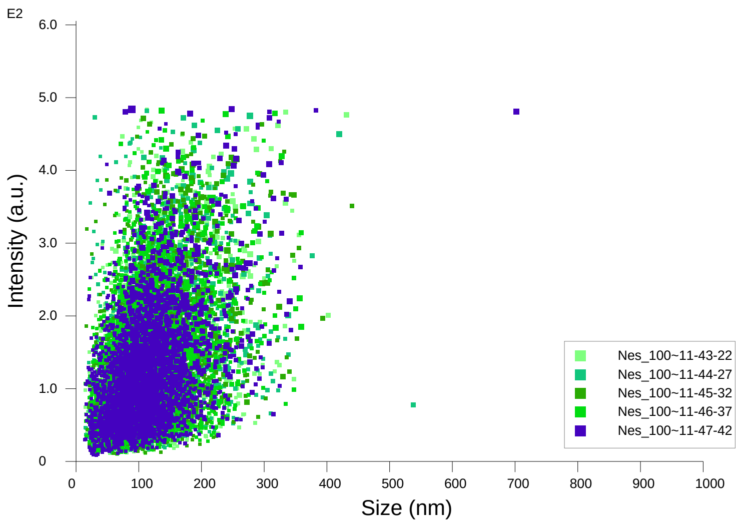


Intensity / Size graph for Experiment:
Nes_1000x 5 60120 2024-10-22 11-40-17

**Script Used: (Full Text):**

SOP Standard Measurement 11-40-17AM 22Oct2024.txt
