## Additional file 9 for "Adeno-Associated Virus Co-Precipitation with Extracellular Vesicles for Genome Editing in Rodent Embryo": pbs 5 60 120 2026-03-09 14-08-15-ExperimentReport.pdf

### NANOSIGHT

### pbs 5 60 120 2026-03-09 14-08-15

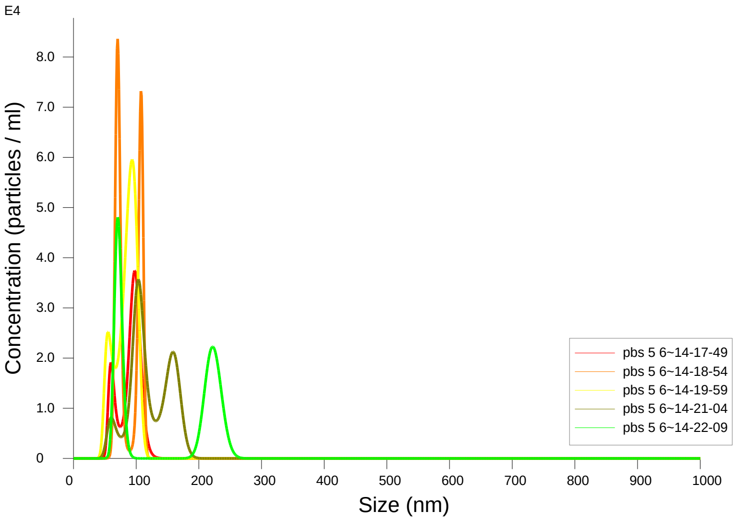


FTLA Concentration / Size graph for Experiment:
pbs 5 60 120 2026-03-09 14-08-15

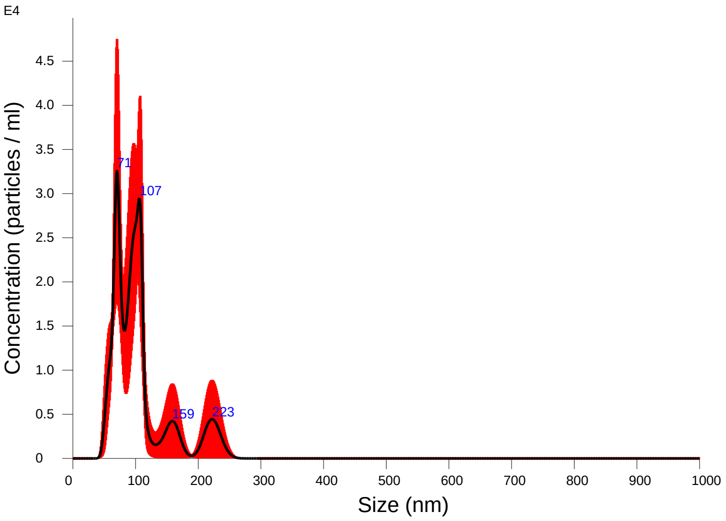


Averaged FTLA Concentration / Size for Experiment:
pbs 5 60 120 2026-03-09 14-08-15
Error bars indicate + / -1 standard error of the mean

#### Included Files

pbs 5 60 120 2026-03-09 14-17-49
pbs 5 60 120 2026-03-09 14-18-54
pbs 5 60 120 2026-03-09 14-19-59
pbs 5 60 120 2026-03-09 14-21-04
pbs 5 60 120 2026-03-09 14-22-09

#### Details

| | |
|---|---|
| NTA Version: | NTA 3.4 - Sample Assistant Build 3.4.4 - SA |
| Script Used: | SOP Standard Measurement 02-08-13PM 09~ |
| Time Captured: | 14:08:15 09/03/2026 |
| Operator: | jitka |
| Pre-treatment: | 1x |
| Sample Name: | pbs |
| Diluent: | pbs |
| Remarks: | |

#### Capture Settings

| | |
|---|---|
| Camera Type: | sCMOS |
| Laser Type: | Blue405 |
| Camera Level: | 12 |
| Slider Shutter: | 1200 |
| Slider Gain: | 146 |
| FPS | 25.0 |
| Number of Frames: | 1498 |
| Temperature: | 25.0 - 25.0 ºC |
| Viscosity: | 1.0 cP |
| Dilution factor: | 1 x 10e0 |
| Syringe Pump Speed: | 120 |

#### Analysis Settings

| | |
|---|---|
| Detect Threshold: | 5 |
| Blur Size: | Auto |
| Max Jump Distance: | Auto: 14.6 pix |

#### Results

Stats: Merged Data

| | |
|---|---|
| Mean: | 104.9 nm |
| Mode: | 70.4 nm |
| SD: | 45.6 nm |
| D10: | 64.6 nm |
| D50: | 94.8 nm |
| D90: | 172.6 nm |

Stats: Mean +/- Standard Error

| | |
|---|---|
| Mean: | 105.6 +/- 12.4 nm |
| Mode: | 87.2 +/- 7.0 nm |
| SD: | 32.4 +/- 11.4 nm |
| D10: | 66.1 +/- 4.1 nm |
| D50: | 93.6 +/- 5.9 nm |
| D90: | 144.1 +/- 25.2 nm |
| Concentration (Upgrade): | 1.63e+06 +/- 1.55e+05 particles/ml |
| | 0.1 +/- 0.0 particles/frame |
| | 0.2 +/- 0.0 centres/frame |

Concentration measurements may be unreliable
See summary file for more info

NANOSIGHT

### pbs 5 60 120 2026-03-09 14-08-15

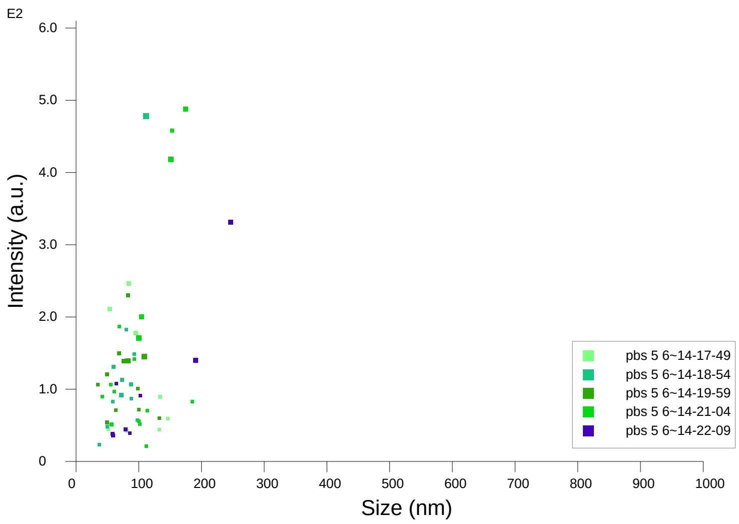


Intensity / Size graph for Experiment:
pbs 5 60 120 2026-03-09 14-08-15

**Script Used: (Full Text):**

SOP Standard Measurement 02-08-13PM 09Mar2026.txt
