## Additional file 9 for "Adeno-Associated Virus Co-Precipitation with Extracellular Vesicles for Genome Editing in Rodent Embryo": PBS-HAT control.pdf

NANOSIGHT

### PBS-HAT 1000x 5 60 120 2024-10-22 12-54-16

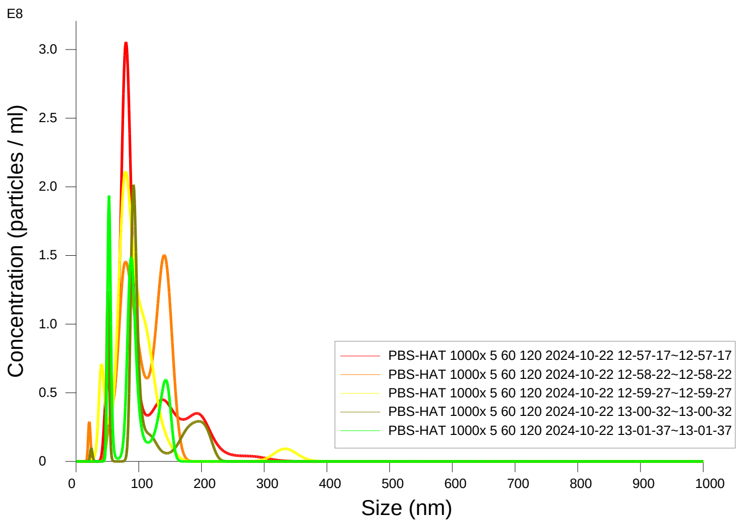


FTLA Concentration / Size graph for Experiment:
PBS-HAT 1000x 5 60 120 2024-10-22 12-54-16

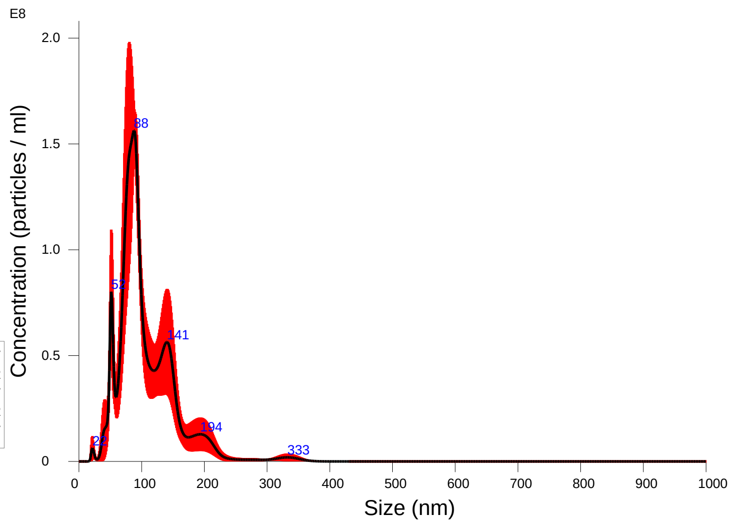


Averaged FTLA Concentration / Size for Experiment:
PBS-HAT 1000x 5 60 120 2024-10-22 12-54-16
Error bars indicate + / -1 standard error of the mean

**Included Files**

PBS-HAT 1000x 5 60 120 2024-10-22 12-57-17
PBS-HAT 1000x 5 60 120 2024-10-22 12-58-22
PBS-HAT 1000x 5 60 120 2024-10-22 12-59-27
PBS-HAT 1000x 5 60 120 2024-10-22 13-00-32
PBS-HAT 1000x 5 60 120 2024-10-22 13-01-37

**Details**

| | |
|---|---|
| NTA Version: | NTA 3.4 - Sample Assistant Build 3.4.4 - SA |
| Script Used: | SOP Standard Measurement 12-54-16PM 22O~ |
| Time Captured: | 12:54:16 22/10/2024 |
| Operator: | Jitka NeburkovÆ |
| Pre-treatment: | |
| Sample Name: | |
| Diluent: | |
| Remarks: | |

**Capture Settings**

| | |
|---|---|
| Camera Type: | sCMOS |
| Laser Type: | Blue405 |
| Camera Level: | 12 |
| Slider Shutter: | 1200 |
| Slider Gain: | 146 |
| FPS | 25.0 |
| Number of Frames: | 1498 |
| Temperature: | 25.0 - 25.0 ºC |
| Viscosity: | 1.0 cP |
| Dilution factor: | 1 x 10e3 |
| Syringe Pump Speed: | 120 |

**Analysis Settings**

| | |
|---|---|
| Detect Threshold: | 5 |
| Blur Size: | Auto |
| Max Jump Distance: | Auto: 14.6 - 22.5 pix |

**Results**

Stats: Merged Data

| | |
|---|---|
| Mean: | 106.1 nm |
| Mode: | 88.0 nm |
| SD: | 46.8 nm |
| D10: | 60.2 nm |
| D50: | 92.6 nm |
| D90: | 159.8 nm |

Stats: Mean +/- Standard Error

| | |
|---|---|
| Mean: | 106.1 +/- 4.2 nm |
| Mode: | 88.7 +/- 14.5 nm |
| SD: | 44.1 +/- 4.9 nm |
| D10: | 59.4 +/- 3.4 nm |
| D50: | 93.8 +/- 3.7 nm |
| D90: | 162.8 +/- 13.9 nm |
| Concentration (Upgrade): | 8.97e+09 +/- 1.49e+09 particles/ml |
| | 0.8 +/- 0.1 particles/frame |
| | 1.0 +/- 0.2 centres/frame |

Concentration measurements may be unreliable
See summary file for more info

### **NANO**SIGHT



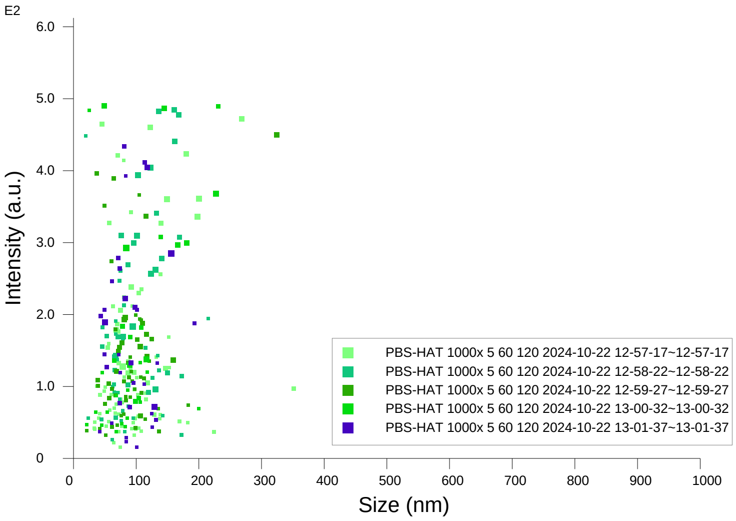


Intensity / Size graph for Experiment:
PBS-HAT 1000x 5 60 120 2024-10-22 12-54-16

**Script Used: (Full Text):**

SOP Standard Measurement 12-54-16PM 22Oct2024.txt
