## Additional file 9 for "Adeno-Associated Virus Co-Precipitation with Extracellular Vesicles for Genome Editing in Rodent Embryo": Sox2-mStrawberry.pdf

### NANOSIGHT

#### Sox2 1000x 5 60 120 2025-07-24 12-33-01

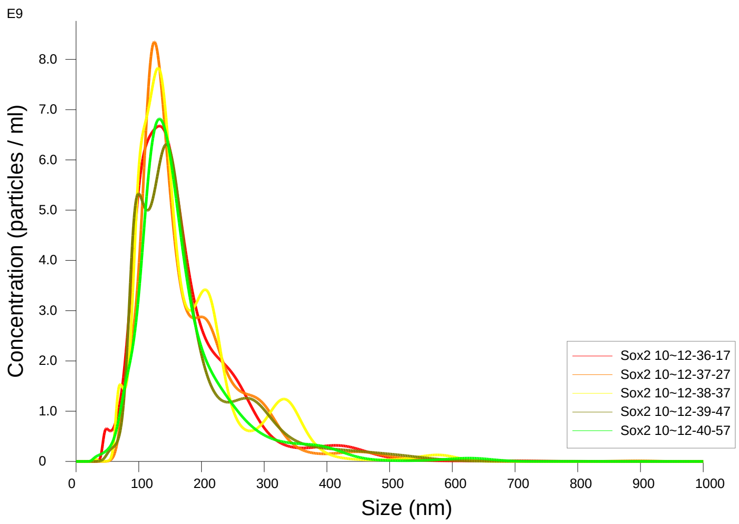


FTLA Concentration / Size graph for Experiment:
Sox2 1000x 5 60 120 2025-07-24 12-33-01

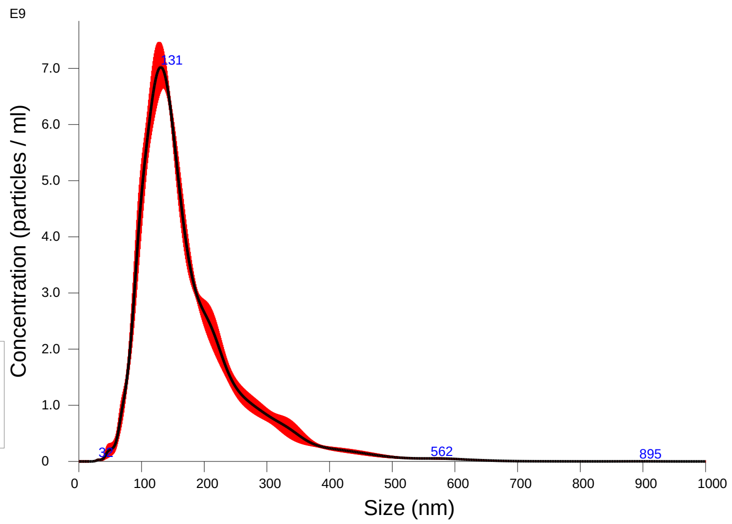


Averaged FTLA Concentration / Size for Experiment:
Sox2 1000x 5 60 120 2025-07-24 12-33-01
Error bars indicate + / -1 standard error of the mean

**Included Files**

Sox2 1000x 5 60 120 2025-07-24 12-36-17
Sox2 1000x 5 60 120 2025-07-24 12-37-27
Sox2 1000x 5 60 120 2025-07-24 12-38-37
Sox2 1000x 5 60 120 2025-07-24 12-39-47
Sox2 1000x 5 60 120 2025-07-24 12-40-57

**Details**

| | |
|---|---|
| NTA Version: | NTA 3.4 - Sample Assistant Build 3.4.4 - SA |
| Script Used: | SOP Standard Measurement 12-32-42PM 24J~ |
| Time Captured: | 12:33:01 24/07/2025 |
| Operator: | jitka |
| Pre-treatment: | |
| Sample Name: | Sox2 |
| Diluent: | water |
| Remarks: | |

**Capture Settings**

| | |
|---|---|
| Camera Type: | sCMOS |
| Laser Type: | Blue405 |
| Camera Level: | 12 |
| Slider Shutter: | 1200 |
| Slider Gain: | 146 |
| FPS | 25.0 |
| Number of Frames: | 1498 |
| Temperature: | 25.0 °C |
| Viscosity: | 1.0 cP |
| Dilution factor: | 1 x 10e3 |
| Syringe Pump Speed: | 120 |

**Analysis Settings**

| | |
|---|---|
| Detect Threshold: | 5 |
| Blur Size: | Auto |
| Max Jump Distance: | Auto: 15.0 - 17.3 pix |

**Results**

Stats: Merged Data

| | |
|---|---|
| Mean: | 176.0 nm |
| Mode: | 130.7 nm |
| SD: | 86.8 nm |
| D10: | 98.4 nm |
| D50: | 150.8 nm |
| D90: | 288.6 nm |

Stats: Mean +/- Standard Error

| | |
|---|---|
| Mean: | 176.0 +/- 1.5 nm |
| Mode: | 133.2 +/- 3.1 nm |
| SD: | 86.7 +/- 1.4 nm |
| D10: | 98.7 +/- 1.6 nm |
| D50: | 150.8 +/- 0.6 nm |
| D90: | 288.4 +/- 7.4 nm |
| Concentration (Upgrade): | 8.25e+11 +/- 2.68e+10 particles/ml |
| | 55.3 +/- 1.6 particles/frame |
| | 58.1 +/- 1.7 centres/frame |

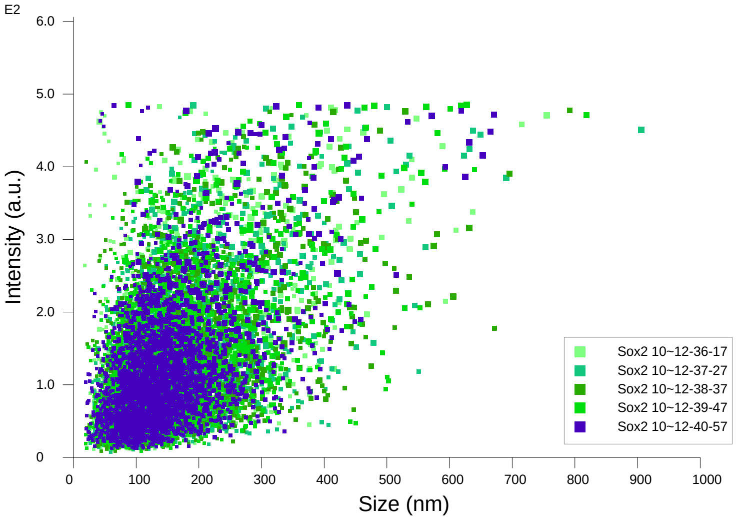


Intensity / Size graph for Experiment:
Sox2 1000x 5 60 120 2025-07-24 12-33-01

**Script Used: (Full Text):**

SOP Standard Measurement 12-32-42PM 24Jul2025.txt
