## Additional file 9 for "Adeno-Associated Virus Co-Precipitation with Extracellular Vesicles for Genome Editing in Rodent Embryo": Ube3a-BioID2.pdf

### NANOSIGHT

#### XO_Ube3a_BioID2_H2_ 2022-10-11 11-46-58

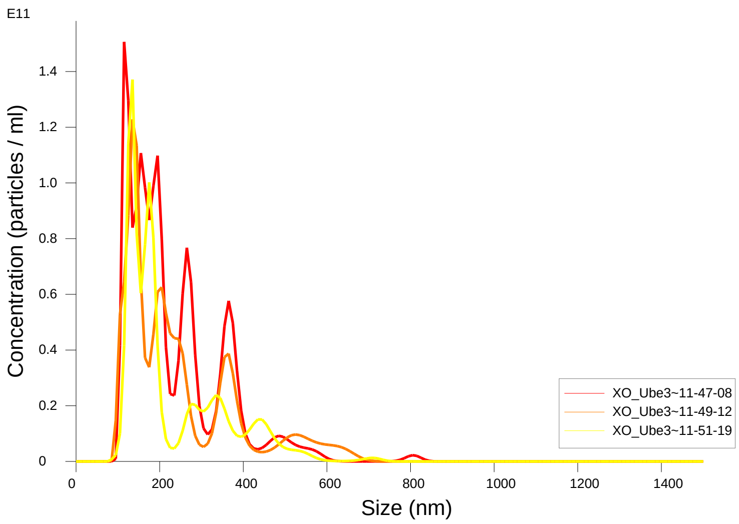


FTLA Concentration / Size graph for Experiment:
XO_Ube3a_BioID2_H2_ 2022-10-11 11-46-58

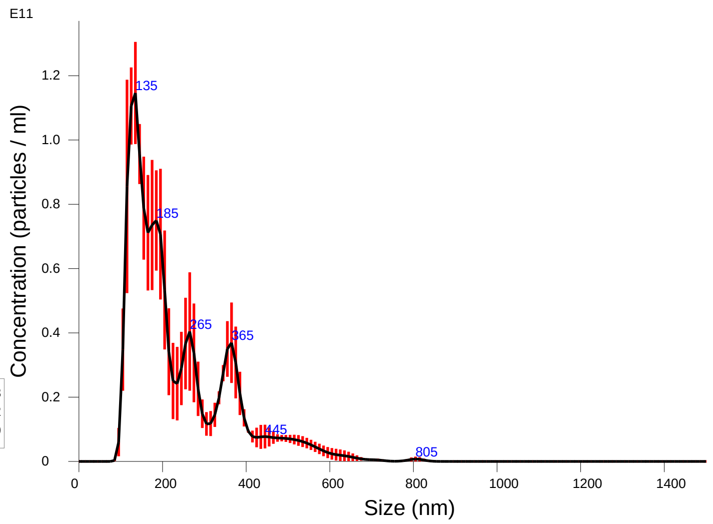


Averaged FTLA Concentration / Size for Experiment:
XO_Ube3a_BioID2_H2_ 2022-10-11 11-46-58
Error bars indicate + / -1 standard error of the mean

##### Included Files

XO_Ube3a_BioID2_H2_ 2022-10-11 11-47-08
XO_Ube3a_BioID2_H2_ 2022-10-11 11-49-12
XO_Ube3a_BioID2_H2_ 2022-10-11 11-51-19

##### Details

| | |
|---|---|
| NTA Version: | NTA 3.4 Build 3.4.4 |
| Script Used: | SOP Standard Measurement 11-46-58AM 11O~ |
| Time Captured: | 11:46:58 11/10/2022 |
| Operator: | Jacopo |
| Pre-treatment: | |
| Sample Name: | |
| Diluent: | 1000 |
| Remarks: | Some aggregate |

##### Capture Settings

| | |
|---|---|
| Camera Type: | sCMOS |
| Laser Type: | Blue405 |
| Camera Level: | 15 |
| Slider Shutter: | 1206 |
| Slider Gain: | 366 |
| FPS | 25.0 |
| Number of Frames: | 1498 |
| Temperature: | 19.3 - 19.4 $^{o}$C |
| Viscosity: | (Water) 1.014 - 1.016 cP |
| Dilution factor: | 1 x 10e3 |

##### Analysis Settings

| | |
|---|---|
| Detect Threshold: | 5 |
| Blur Size: | Auto |
| Max Jump Distance: | Auto: 6.7 - 6.9 pix |

##### Results

Stats: Merged Data

| | |
|---|---|
| Mean: | 235.2 nm |
| Mode: | 131.8 nm |
| SD: | 122.5 nm |
| D10: | 122.1 nm |
| D50: | 190.6 nm |
| D90: | 398.2 nm |

Stats: Mean +/- Standard Error

| | |
|---|---|
| Mean: | 235.3 +/- 4.7 nm |
| Mode: | 129.7 +/- 5.8 nm |
| SD: | 122.7 +/- 6.3 nm |
| D10: | 122.2 +/- 1.8 nm |
| D50: | 191.2 +/- 7.5 nm |
| D90: | 418.6 +/- 22.1 nm |
| Concentration: | 1.50e+12 +/- 2.05e+11 particles/ml |
| | 82.0 +/- 11.2 particles/frame |
| | 93.6 +/- 15.4 centres/frame |

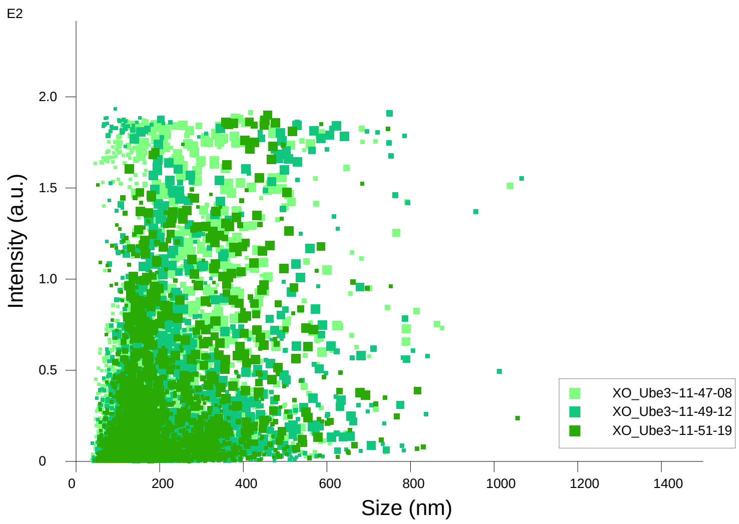


Intensity / Size graph for Experiment:
XO_Ube3a_BioID2_H2_ 2022-10-11 11-46-58

**Script Used: (Full Text):**

SOP Standard Measurement 11-46-58AM 11Oct2022.txt
