## Additional file 9 for "Adeno-Associated Virus Co-Precipitation with Extracellular Vesicles for Genome Editing in Rodent Embryo": Umod Y178 5 60 120 2000x 2026-03-09 13-30-48-ExperimentReport.pdf

**NANOSIGHT**

### Umod 4178 5 60 120 2000x 2026-03-09 13-30-48

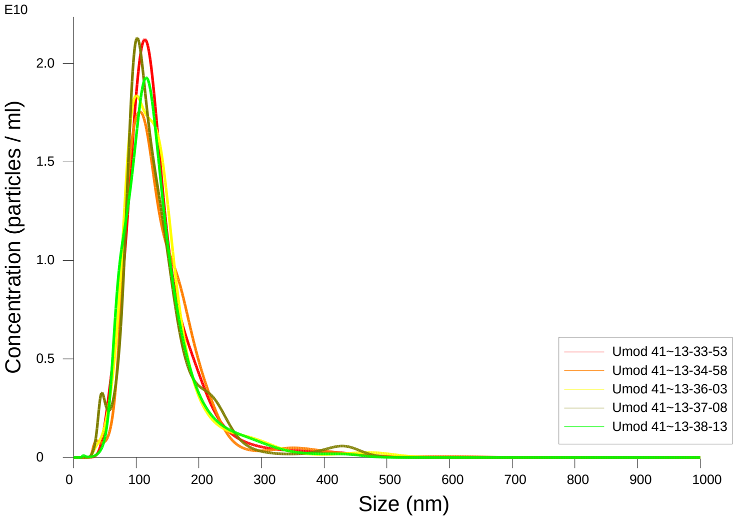


FTLA Concentration / Size graph for Experiment:
Umod 4178 5 60 120 2000x 2026-03-09 13-30-48

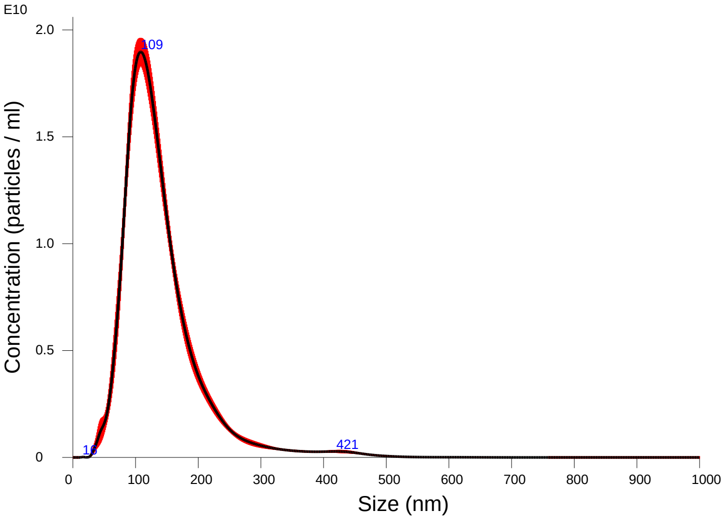


Averaged FTLA Concentration / Size for Experiment:
Umod 4178 5 60 120 2000x 2026-03-09 13-30-48
Error bars indicate + / -1 standard error of the mean

#### Included Files

Umod 4178 5 60 120 2000x 2026-03-09 13-33-53
Umod 4178 5 60 120 2000x 2026-03-09 13-34-58
Umod 4178 5 60 120 2000x 2026-03-09 13-36-03
Umod 4178 5 60 120 2000x 2026-03-09 13-37-08
Umod 4178 5 60 120 2000x 2026-03-09 13-38-13

#### Details

| | |
|---|---|
| NTA Version: | NTA 3.4 - Sample Assistant Build 3.4.4 - SA |
| Script Used: | SOP Standard Measurement 01-30-46PM 09~ |
| Time Captured: | 13:30:48 09/03/2026 |
| Operator: | jitka |
| Pre-treatment: | 2000x |
| Sample Name: | 4178 |
| Diluent: | pbs |
| Remarks: | |

#### Capture Settings

| | |
|---|---|
| Camera Type: | sCMOS |
| Laser Type: | Blue405 |
| Camera Level: | 12 |
| Slider Shutter: | 1200 |
| Slider Gain: | 146 |
| FPS | 25.0 |
| Number of Frames: | 1498 |
| Temperature: | 25.0 - 25.0 °C |
| Viscosity: | 1.0 cP |
| Dilution factor: | 2 x 10e3 |
| Syringe Pump Speed: | 120 |

#### Analysis Settings

| | |
|---|---|
| Detect Threshold: | 5 |
| Blur Size: | Auto |
| Max Jump Distance: | Auto: 18.2 - 19.8 pix |

#### Results

Stats: Merged Data

| | |
|---|---|
| Mean: | 139.4 nm |
| Mode: | 108.3 nm |
| SD: | 64.8 nm |
| D10: | 81.5 nm |
| D50: | 124.7 nm |
| D90: | 210.1 nm |

Stats: Mean +/- Standard Error

| | |
|---|---|
| Mean: | 139.4 +/- 1.2 nm |
| Mode: | 107.6 +/- 3.2 nm |
| SD: | 64.7 +/- 1.7 nm |
| D10: | 81.5 +/- 0.9 nm |
| D50: | 124.8 +/- 1.0 nm |
| D90: | 209.9 +/- 1.6 nm |
| Concentration (Upgrade): | 1.85e+12 +/- 2.62e+10 particles/ml |
| | 77.2 +/- 0.8 particles/frame |
| | 75.6 +/- 0.8 centres/frame |

### Umod 4178 5 60 120 2000x 2026-03-09 13-30-48

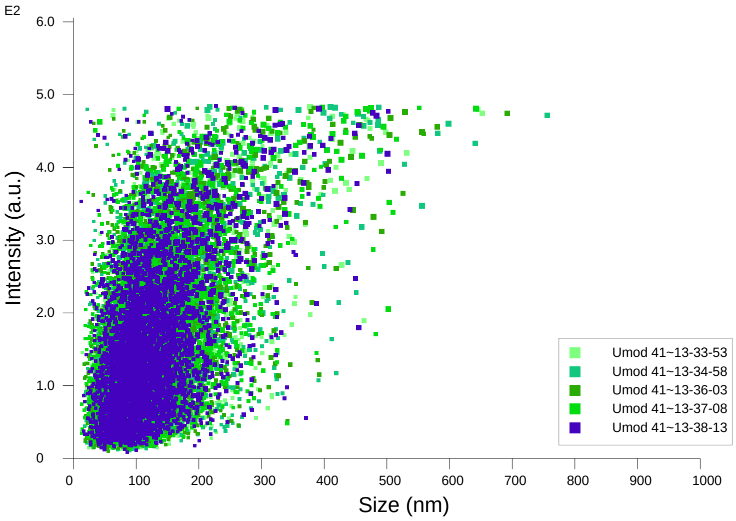


Intensity / Size graph for Experiment:
Umod 4178 5 60 120 2000x 2026-03-09 13-30-48

**Script Used: (Full Text):**

SOP Standard Measurement 01-30-46PM 09Mar2026.txt
