## Additional file 9 for "Adeno-Associated Virus Co-Precipitation with Extracellular Vesicles for Genome Editing in Rodent Embryo": Cypces 1000x 5 60 120 2025-07-24 13-35-08-ExperimentReport.pdf

NANOSIGHT

### Cypces 1000x 5 60 120 2025-07-24 13-35-08

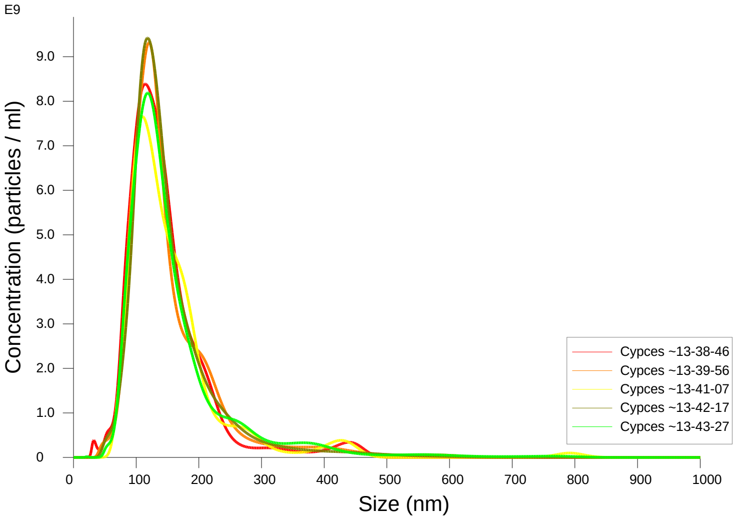


FTLA Concentration / Size graph for Experiment:
Cypces 1000x 5 60 120 2025-07-24 13-35-08

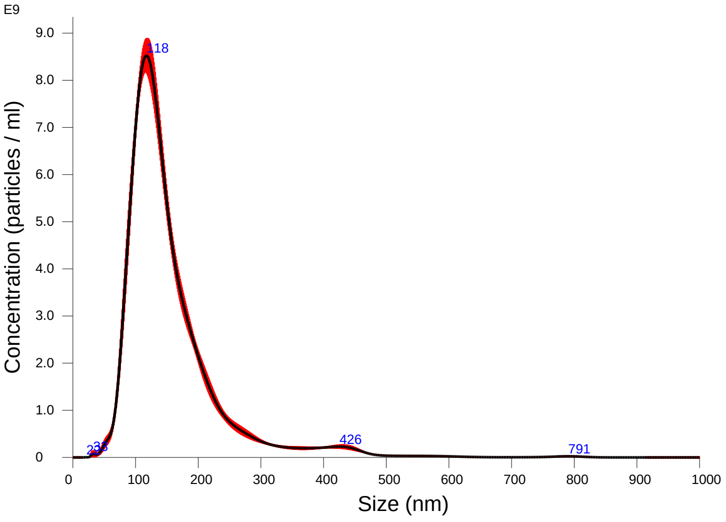


Averaged FTLA Concentration / Size for Experiment:
Cypces 1000x 5 60 120 2025-07-24 13-35-08
Error bars indicate + / -1 standard error of the mean

#### Included Files

Cypces 1000x 5 60 120 2025-07-24 13-38-46
Cypces 1000x 5 60 120 2025-07-24 13-39-56
Cypces 1000x 5 60 120 2025-07-24 13-41-07
Cypces 1000x 5 60 120 2025-07-24 13-42-17
Cypces 1000x 5 60 120 2025-07-24 13-43-27

#### Details

| | |
|---|---|
| NTA Version: | NTA 3.4 - Sample Assistant Build 3.4.4 - SA |
| Script Used: | SOP Standard Measurement 01-34-53PM 24J~ |
| Time Captured: | 13:35:08 24/07/2025 |
| Operator: | jitka |
| Pre-treatment: | |
| Sample Name: | |
| Diluent: | pbs |
| Remarks: | |

#### Capture Settings

| | |
|---|---|
| Camera Type: | sCMOS |
| Laser Type: | Blue405 |
| Camera Level: | 12 |
| Slider Shutter: | 1200 |
| Slider Gain: | 146 |
| FPS | 25.0 |
| Number of Frames: | 1498 |
| Temperature: | 25.0 - 25.0 °C |
| Viscosity: | 1.0 cP |
| Dilution factor: | 1 x 10e3 |
| Syringe Pump Speed: | 120 |

#### Analysis Settings

| | |
|---|---|
| Detect Threshold: | 5 |
| Blur Size: | Auto |
| Max Jump Distance: | Auto: 15.2 - 18.3 pix |

#### Results

Stats: Merged Data

| | |
|---|---|
| Mean: | 155.5 nm |
| Mode: | 117.4 nm |
| SD: | 80.4 nm |
| D10: | 91.3 nm |
| D50: | 134.1 nm |
| D90: | 235.9 nm |

Stats: Mean +/- Standard Error

| | |
|---|---|
| Mean: | 155.6 +/- 2.4 nm |
| Mode: | 116.3 +/- 1.9 nm |
| SD: | 79.9 +/- 4.6 nm |
| D10: | 91.4 +/- 1.0 nm |
| D50: | 134.3 +/- 0.9 nm |
| D90: | 238.2 +/- 7.3 nm |
| Concentration (Upgrade): | 8.22e+11 +/- 1.23e+10 particles/ml |
| | 54.8 +/- 0.7 particles/frame |
| | 58.6 +/- 0.4 centres/frame |

### NANOSIGHT

Cypces 1000x 5 60 120 2025-07-24 13-35-08

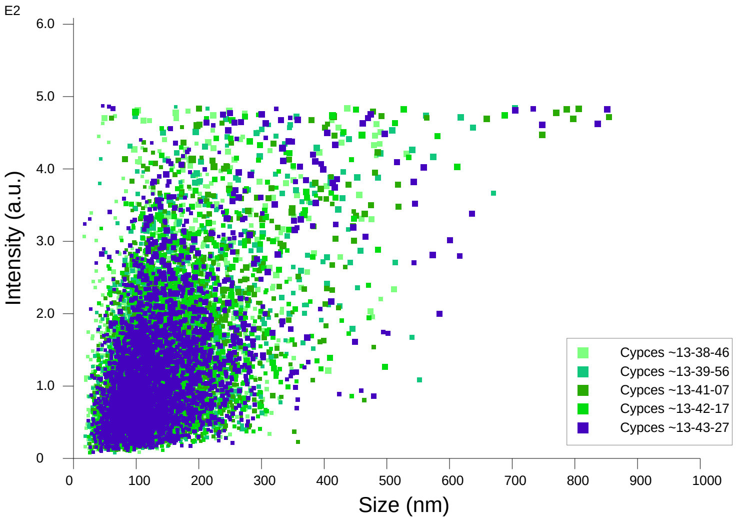


Intensity / Size graph for Experiment:
Cypces 1000x 5 60 120 2025-07-24 13-35-08

**Script Used: (Full Text):**

SOP Standard Measurement 01-34-53PM 24Jul2025.txt
