## Additional file 9 for "Adeno-Associated Virus Co-Precipitation with Extracellular Vesicles for Genome Editing in Rodent Embryo": EV Sox2-mStr 6h 1000x 5 60 120 2024-12-18 11-42-32-ExperimentReport.pdf

### NANOSIGHT

#### EV Sox2-mStr 6h 1000x 5 60 120 2024-12-18 11-42-32

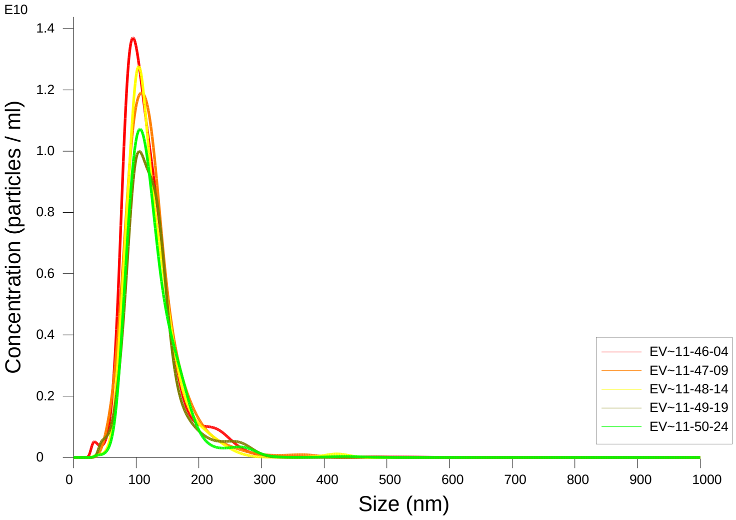


FTLA Concentration / Size graph for Experiment:
EV Sox2-mStr 6h 1000x 5 60 120 2024-12-18 11-42-32

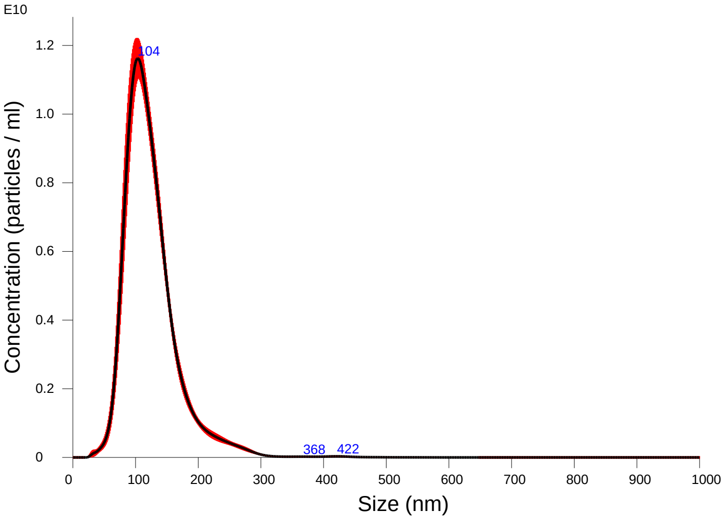


Averaged FTLA Concentration / Size for Experiment:
EV Sox2-mStr 6h 1000x 5 60 120 2024-12-18 11-42-32
Error bars indicate + / -1 standard error of the mean

##### Included Files

EV Sox2-mStr 6h 1000x 5 60 120 2024-12-18 11-46-04
EV Sox2-mStr 6h 1000x 5 60 120 2024-12-18 11-47-09
EV Sox2-mStr 6h 1000x 5 60 120 2024-12-18 11-48-14
EV Sox2-mStr 6h 1000x 5 60 120 2024-12-18 11-49-19
EV Sox2-mStr 6h 1000x 5 60 120 2024-12-18 11-50-24

##### Details

| | |
|---|---|
| NTA Version: | NTA 3.4 - Sample Assistant Build 3.4.4 - SA |
| Script Used: | SOP Standard Measurement 11-42-13AM 18~ |
| Time Captured: | 11:42:32 18/12/2024 |
| Operator: | Jitka |
| Pre-treatment: | |
| Sample Name: | |
| Diluent: | pbs |
| Remarks: | |

##### Capture Settings

| | |
|---|---|
| Camera Type: | sCMOS |
| Laser Type: | Blue405 |
| Camera Level: | 12 |
| Slider Shutter: | 1200 |
| Slider Gain: | 146 |
| FPS | 25.0 |
| Number of Frames: | 1498 |
| Temperature: | 25.0 - 25.0 °C |
| Viscosity: | 1.0 cP |
| Dilution factor: | 1 x 10e3 |
| Syringe Pump Speed: | 120 |

##### Analysis Settings

| | |
|---|---|
| Detect Threshold: | 5 |
| Blur Size: | Auto |
| Max Jump Distance: | Auto: 16.4 - 19.2 pix |

##### Results

Stats: Merged Data

| | |
|---|---|
| Mean: | 124.1 nm |
| Mode: | 103.6 nm |
| SD: | 44.2 nm |
| D10: | 81.4 nm |
| D50: | 115.7 nm |
| D90: | 174.3 nm |

Stats: Mean +/- Standard Error

| | |
|---|---|
| Mean: | 124.3 +/- 1.0 nm |
| Mode: | 103.8 +/- 2.3 nm |
| SD: | 44.0 +/- 0.5 nm |
| D10: | 81.9 +/- 1.2 nm |
| D50: | 115.8 +/- 1.3 nm |
| D90: | 174.3 +/- 0.8 nm |
| Concentration (Upgrade): | 8.95e+11 +/- 4.82e+10 particles/ml |
| | 62.1 +/- 3.2 particles/frame |
| | 63.7 +/- 2.7 centres/frame |

NANOSIGHT

EV Sox2-mStr 6h 1000x 5 60 120 2024-12-18 11-42-32

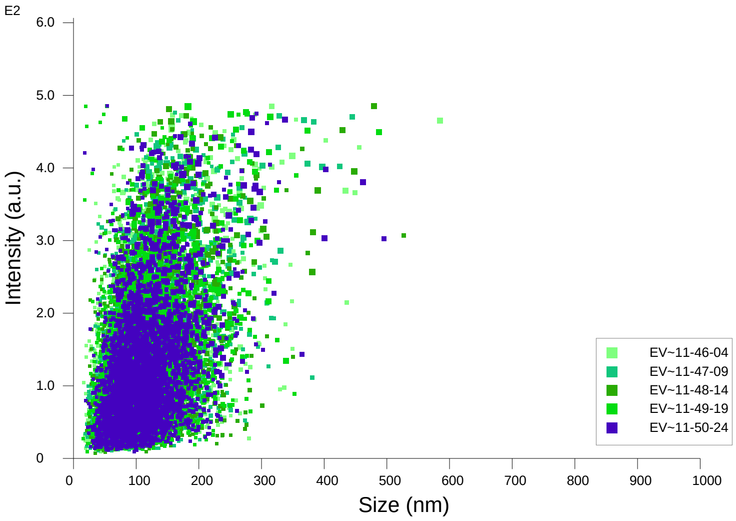


Intensity / Size graph for Experiment:
EV Sox2-mStr 6h 1000x 5 60 120 2024-12-18 11-42-32

**Script Used: (Full Text):**

SOP Standard Measurement 11-42-13AM 18Dec2024.txt
