## Additional file 9 for "Adeno-Associated Virus Co-Precipitation with Extracellular Vesicles for Genome Editing in Rodent Embryo": Umod C126 5 60 120 2000x 2026-03-09 13-47-44-ExperimentReport.pdf

NANOSIGHT

### Umod C126 5 60 120 2000x 2026-03-09 13-47-44

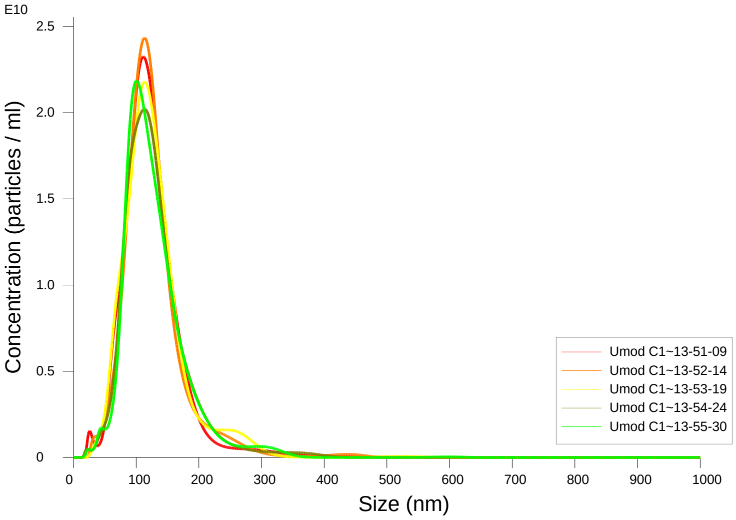


FTLA Concentration / Size graph for Experiment:
Umod C126 5 60 120 2000x 2026-03-09 13-47-44

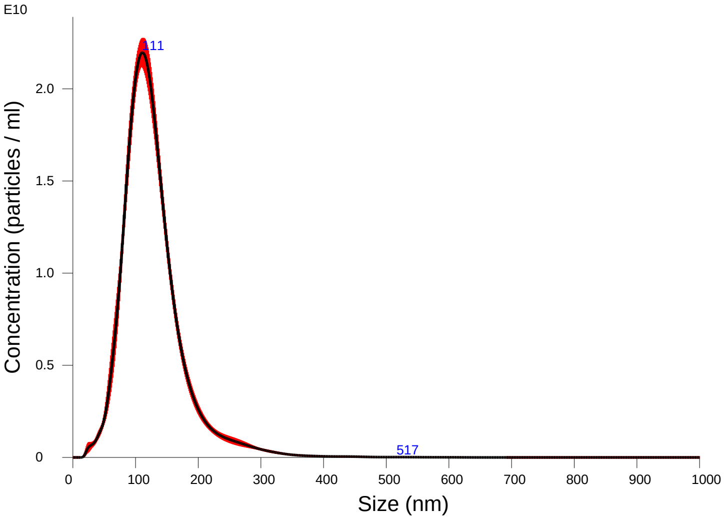


Averaged FTLA Concentration / Size for Experiment:
Umod C126 5 60 120 2000x 2026-03-09 13-47-44
Error bars indicate + / -1 standard error of the mean

#### Included Files

Umod C126 5 60 120 2000x 2026-03-09 13-51-09
Umod C126 5 60 120 2000x 2026-03-09 13-52-14
Umod C126 5 60 120 2000x 2026-03-09 13-53-19
Umod C126 5 60 120 2000x 2026-03-09 13-54-24
Umod C126 5 60 120 2000x 2026-03-09 13-55-30

#### Details

NTA Version: NTA 3.4 - Sample Assistant Build 3.4.4 - SA
Script Used: SOP Standard Measurement 01-47-42PM 09~
Time Captured: 13:47:44 09/03/2026
Operator: jitka
Pre-treatment: 2000x
Sample Name: C126
Diluent: pbs
Remarks:

#### Capture Settings

Camera Type: sCMOS
Laser Type: Blue405
Camera Level: 12
Slider Shutter: 1200
Slider Gain: 146
FPS 25.0
Number of Frames: 1498
Temperature: 25.0 - 25.0 $^{o}$C
Viscosity: 1.0 cP
Dilution factor: 2 x 10e3
Syringe Pump Speed: 120

#### Analysis Settings

Detect Threshold: 5
Blur Size: Auto
Max Jump Distance: Auto: 19.1 - 21.6 pix

#### Results

Stats: Merged Data
Mean: 127.5 nm
Mode: 111.0 nm
SD: 51.0 nm
D10: 77.8 nm
D50: 118.9 nm
D90: 183.2 nm

Stats: Mean +/- Standard Error
Mean: 127.5 +/- 0.7 nm
Mode: 110.5 +/- 2.5 nm
SD: 51.0 +/- 0.3 nm
D10: 77.7 +/- 1.2 nm
D50: 118.9 +/- 0.4 nm
D90: 183.4 +/- 2.4 nm
Concentration (Upgrade): 1.92e+12 +/- 2.48e+10 particles/ml
73.7 +/- 0.9 particles/frame
70.9 +/- 0.5 centres/frame

### NANOSIGHT



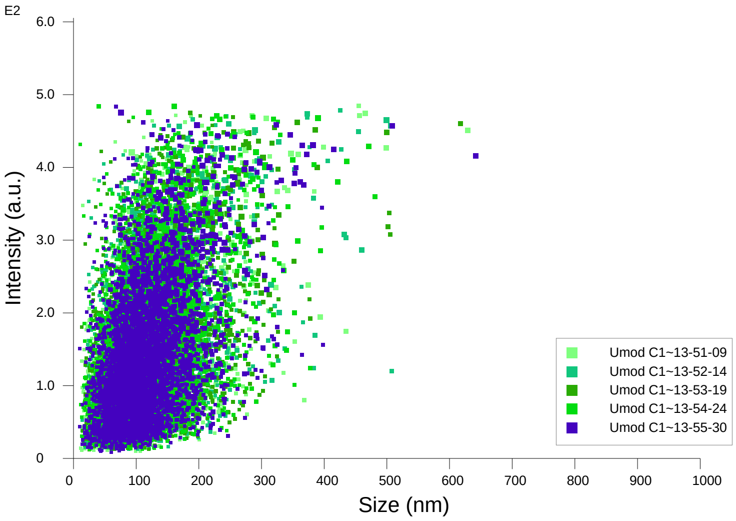


Intensity / Size graph for Experiment:
Umod C126 5 60 120 2000x 2026-03-09 13-47-44

**Script Used: (Full Text):**

SOP Standard Measurement 01-47-42PM 09Mar2026.txt
